# Haplogenome assembly reveals interspecific structural variation in *Eucalyptus* hybrids

**DOI:** 10.1101/2022.08.17.501336

**Authors:** Anneri Lötter, Tuan A. Duong, Julia Candotti, Eshchar Mizrachi, Jill L. Wegrzyn, Alexander A. Myburg

## Abstract

*De novo* phased (haplo)genome assembly using long-read DNA sequencing data has improved the detection and characterization of structural variants (SVs) in plant and animal genomes. Able to span across haplotypes, long reads allow phased, haplogenome assembly in highly outbred organisms such as forest trees. *Eucalyptus* tree species and interspecific hybrids are the most widely planted hardwood trees with F_1_ hybrids of *Eucalyptus grandis* and *E. urophylla* forming the bulk of fast-growing pulpwood plantations in subtropical regions. The extent of structural variation and its effect on interspecific hybridization is unknown in these trees. As a first step towards elucidating the extent of structural variation between the genomes of *E. grandis* and *E. urophylla*, we sequenced and assembled the haplogenomes contained in an F_1_ hybrid of the two species. Using Nanopore sequencing and a trio-binning approach, we assembled the separate haplogenomes (567 Mb and 545 Mb) to 98.8% BUSCO completion. High-density SNP genetic linkage maps of both parents allowed scaffolding of 88% of the haplogenome contigs into 11 pseudo-chromosomes (scaffold N_50_ of 43.82 Mb and 42.45 Mb for the *E. grandis* and *E. urophylla* haplogenomes, respectively). We identify 48,729 SVs between the two haplogenomes providing the first detailed insight into genome structural rearrangement in these species. The two haplogenomes have similar gene content, 35,572 and 33,915 functionally annotated genes, of which 34% are contained in genome rearrangements. Knowledge of SV and haplotype diversity in the two species will form the basis for understanding the genetic basis of hybrid superiority in these trees.

**Significance statement:** We have produced phased, haplogenome assemblies of an interspecific F1 hybrid using a trio-binning approach and performed the first genome-wide analysis of genome synteny between a subtropical *Eucalyptus* tree species, *E. grandis,* and a tropical eucalypt, *E. urophylla.* This revealed a large number of previously undescribed genome structural variants as a step towards understanding genome structural evolution in this iconic genus of fast-growing woody perennials.

## Introduction

There is considerable pressure to improve crop yields to provide food, fibre, shelter and renewable energy for the growing human population (Grierson *et al*., 2011) in a sustainable manner. Fast-growing *Eucalyptus* tree species provide an important renewable feedstock for biomaterial (timber, fibre and lignocellulosics) and bioenergy production, relieving pressure on native forests (Grattapaglia and Kirst, 2008). These species, commonly referred to as eucalypts, constitute the most widely planted hardwood fibre crop globally. The most productive plantation areas are planted with interspecific F_1_ hybrid clones that combine favourable characteristics of parental species and generally lead to increased forest productivity and product quality, and reduced production costs (de Assis, 2000, Grattapaglia and Kirst, 2008). The most widely planted hybrid combination in subtropical regions, *E. grandis* x *E. urophylla,* is primarily bred to combine the disease resistance of the tropical species *E. urophylla* with the fast growth of the subtropical species *E. grandis*. To further improve plantation productivity, wood quality and resilience, more efficient breeding strategies have been pursued in the past decade, primarily through genomic selection using genome-wide SNP markers (Rezende *et al*., 2014, Grattapaglia *et al*., 2018).

Discriminating the maternal and paternal chromosome copies (defined by haplotypes or blocks of allelic variants that are inherited together; Zheng *et al*., 2016) allows for identification of haplotype and structural variants that may be associated with crop productivity and resilience (Jiao and Schneeberger, 2017, Alonge *et al*., 2020). Haplotype-based molecular breeding has been shown to be a more accurate and effective breeding strategy (Ogawa *et al*., 2018a, Ogawa *et al*., 2018b) compared to SNP based strategies. Haplotypes can often be inferred accurately in offspring by using the parental genomes and previously defined SNP tag-markers and haplotype imputation with statistical methods (Motazedi *et al*., 2018). SNP tag-markers can then be used in accelerated breeding strategies by aiding the selection of progeny for propagation and deployment, or identification of parents for further breeding (Bevan *et al*., 2017).

Access to multiple high-quality reference genome assemblies facilitates the identification of haplotypes and structural variants, both of which underlie pan-genome variation in plants. Genome assembly in highly outbred organisms such as forest trees, is often hampered by high levels of heterozygosity and the frequent occurrence of non-syntenic DNA sequences in intergenic regions leading to mixed phase contigs. As a consequence, many of the available reference sequences of outbred plants do not accurately reflect the haplogenomes carried by the reference individuals (Kyriakidou *et al*., 2018). Long-read sequencing (LRS) technologies such as Oxford Nanopore (ONT) and Pacific Biosciences (PacBio) can mitigate the challenges associated with assembling outbred plant genomes. Long reads can span across multiple syntenic (gene) regions and connect intergenic allelic variants between them, allowing separate, phased assembly of haplotype and structural variant alternatives. The growing number of phased genome assemblies, especially those assembled with LRS data, has revealed that a single flat reference genome misses a substantial portion of the genotypic diversity in outbred species (Sherman and Salzberg, 2020). As such, there is a movement towards assembly of a pan-reference genome, which incorporates variants from multiple individuals as has been reported in humans (reviews by Sherman and Salzberg, 2020) and plants (reviewed by Bayer *et al*., 2020).

Studies on pan-genomic (including haplotype and structural) variation are still lacking in *Eucalyptus*, with most information on genome synteny still derived from genetic linkage mapping. These studies have suggested that there is high collinearity between eucalypt species, including *E. grandis* and *E. urophylla* (Brondani *et al*., 1998, Marques *et al*., 2002, Hudson *et al*., 2011, Bartholome *et al*., 2015). However, the degree of fine scale synteny between *E. grandis* and *E. urophylla* is unknown as there is not a *de novo* reference assembly available for *E. urophylla*, one of the most important hybrid parent partners. The current reference genome, *E. grandis* (Myburg *et al*., 2014), was sequenced with a combination of Sanger and SRS. These technologies have limited capability to identify haplotype and structural variants (reviewed by Ho *et al*., 2020). The lack of available LRS based genome assemblies for *E. grandis* and *E. urophylla* have precluded studies of pan-genome variation in these species and their F_1_ hybrids.

Combining SRS and LRS data with a parent-offspring trio-sequencing approach has been demonstrated to allow assembly of high-quality haplo-reference genomes for the two parents, at a lower cost than generating two independent reference quality genomes (Koren *et al*., 2017, Shirasawa *et al*., 2019, Zhu *et al*., 2019). Similarly, trio-sequencing of an interspecific F_1_ hybrid of *E. grandis* and *E. urophylla,* paired with LRS technologies will generate high-quality assemblies of the haplogenomes contained in the F_1_ hybrid. These high-quality phased genome assemblies will ultimately provide a basis for pursuing haplotype-based molecular breeding of eucalypt trees and will provide preliminary insights into the abundance and distribution of structural variants (SVs) of consequence to breeding. Thus, the aim of this study was to create a starting point for defining pan-genome, haplotype and structural variation in *E. grandis* and *E. urophylla*.

## Results

### Genome sequencing

Illumina sequencing of an F_1_ hybrid individual (SAP_F1_FK118) and its pure-species *E. grandis* (SAP_GRA_FK1758) and *E. urophylla* (SAP_URO_FK1756) parents (Sappi Forest Research, South Africa) resulted in more than 116 Gb of PE150 data per individual (Supplementary Table 1). Using GenomeScope2.0, we estimated the genome size to be 443.19 Mb, 482.27 Mb and 477.76 Mb for the *E. urophylla, E. grandis* parents and the F_1_ hybrid respectively (Supplementary Figure 1). These short read-based estimates were substantially smaller than previous estimates based on flow cytometry (Grattapaglia and Bradshaw Jr, 1994) and that reported for the *E. grandis* reference genome (Myburg *et al*., 2014). Marks *et al*., (2021) recently reported a lower flow cytometry size estimate (497.7 Mb) for *E. grandis* supporting our findings. Levels of heterozygosity in the short-read data were 2.14%, 2.63% and 3.46% for the *E. grandis*, *E. urophylla* and the F_1_ hybrid (Supplementary Figure 1) providing ample genetic diversity for trio-binning of the long-reads (see below).

A total of 75.32 Gb of Nanopore sequencing data was generated (read N_50_ ∼27 kb), of which 68.15 Gb (90.48%) passed QC (Q-value > 7, Supplementary Table 2) and was used for trio-binning corresponding to ∼105X coverage of the F_1_ hybrid genome and ∼50X coverage per haplogenome (Figure 1, Supplementary Table 2).

**Figure 1.**
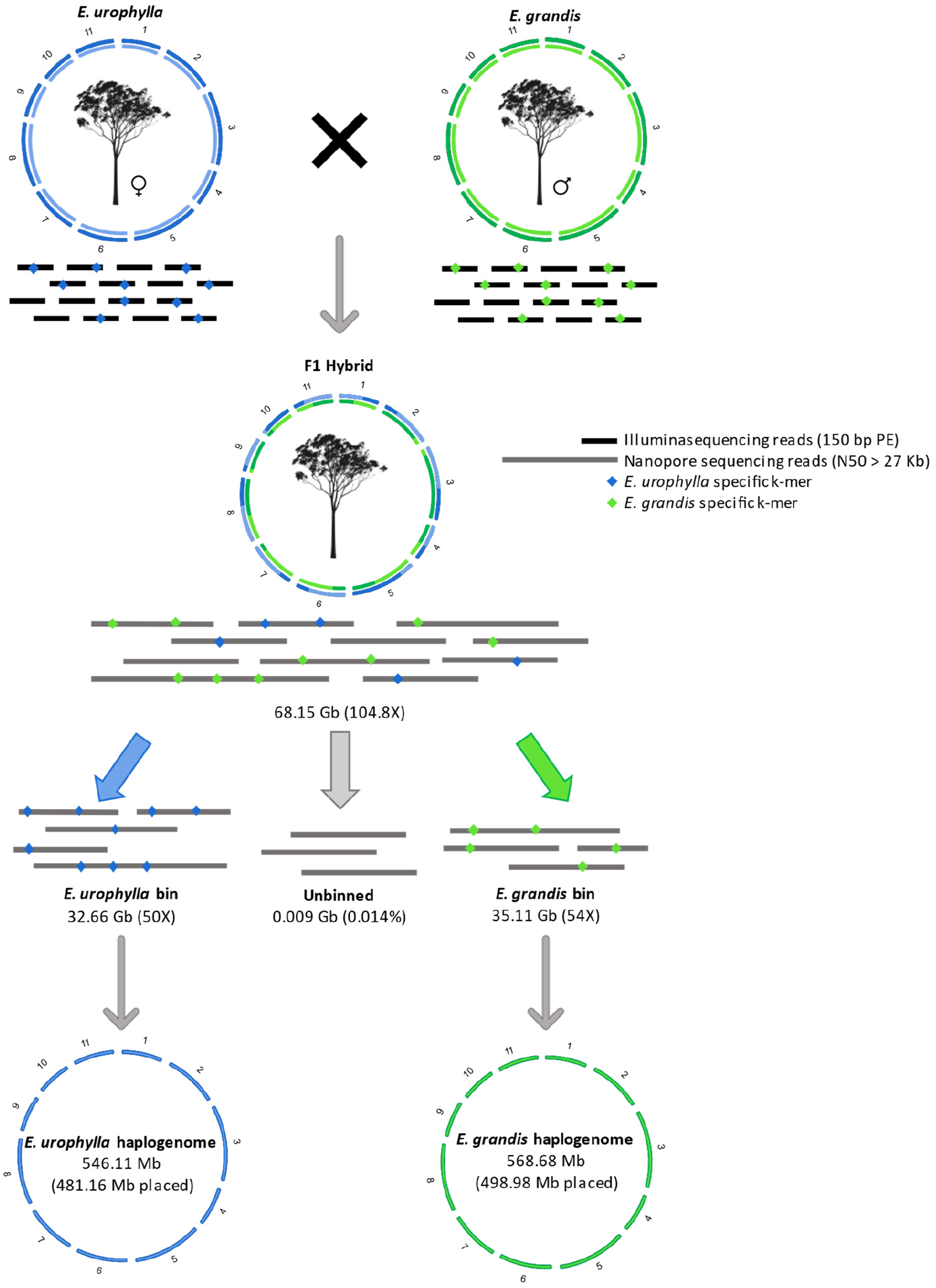
Separation of *E. urophylla* and *E. grandis* haplogenomes in the F_1_ hybrid using a trio-binning strategy. Using whole-genome Illumina short-read sequencing data of the parental genomes, long-read sequencing data of the F_1_ hybrid offspring is separated based on unique parental k-mers into *E. urophylla* and *E. grandis* haplotype bins (amount of Nanopore sequencing data is indicated in gigabases (Gb) below each bin, as well as the estimated genome coverage). Reads that contain no unique k-mers were unbinned and kept in their own bin. Long reads were subsequently assembled independently, resulting in fully assembled *E. urophylla* and *E. grandis* haplogenome (total assembly size is shown below the relevant haplogenome and size of assembly scaffolded into eleven chromosomes are indicated in brackets). This figure is adapted from Koren *et al*., (2018), and tree images are from https://rooweb.com.au/.

### Genome assembly

#### Phased hybrid genome assembly using trio-binning

To separately assemble the long reads originating from the two haplogenomes in the F_1_ hybrid, we performed trio-binning using the Illumina short-read data for the parents and the long-read data for the F_1_ individual. We were able to bin 1,876,816 reads (32.66 Gb) for the *E. urophylla* haplogenome and 1,998,860 reads (35.11 Gb) for the *E. grandis* haplogenome corresponding to 50X and 54X coverage of the two haplotypes, respectively (Figure 1, Supplementary Table 3). Only 6,693 reads (0.014%) could not be binned and were excluded from further analyses.

Assembly of the binned reads for the *E. urophylla* haplogenome resulted in 654 contigs and a total size of 546.1 Mb, with a contig N_50_ of 4.41 Mb (Table 1). A BUSCO completeness score of 99.2% was obtained of which 95.2% were single-copy genes and only 4.0% were duplicate-copy genes (Supplementary Figure 2). The reads binned for the *E. grandis* haplogenome assembled into 793 contigs with a total size of 568.5 Mb and a contig N_50_ of 3.91 Mb (Table 1). For this assembly we obtained a BUSCO completeness score of 99.0%, of which 94.4% were single copy genes and 4.6% were duplicate genes (Supplementary Figure 2). The low duplicate percentages reflected efficient trio-binning and haplogenome assembly.

**Table 1.** Assembly and annotation statistics for the *E. urophylla* and *E. grandis* haplogenomess compared to the previous *E. grandis* genome assembly.

|  | *E. grandis* v2.0 | *E. grandis* | *E. urophylla* |
| --- | --- | --- | --- |
| Type of sequencing | BAC end cloning (ABI) | Illumina + ONP | Illumina + ONP |
| Genome coverage<sup>a</sup> | 6.73x | 54.01x (ONP) | 50.25x (ONP) |
| Primary assembly: |  |  |  |
| Number of contigs | 32,724 | 793 | 654 |
| Total number bases in contigs | 691.43 Mb | 568.46 Mb | 546.11 Mb |
| Contig N50 length | 67.25 kb | 3.91 Mb | 4.41 Mb |
| Contig L50 | 2,261 | 38 | 36 |
| Total contigs > 50 kb | 288 | 387 | 368 |
| Validated contigs (Polar_Star): |  |  |  |
| Number of contigs | - | 1,579 | 1,418 |
| Total number bases in contigs | - | 566.72 Mb | 544.51 Mb |
| Contig N50 length | - | 2.42 Mb | 1.93 Mb |
| Contig L50 | - | 74 | 83 |
| Total contigs > 50 kb | - | 522 | 547 |
| Assembly BUSCO completeness<sup>b</sup> | 98.80% | 98.70% | 99.20% |
| Number scaffolds | 4,951 | 1,279 | 1,078 |
| Total number of bases scaffolded<sup>c</sup> | 612.60 Mb | 498.98 Mb | 481.16 Mb |
| Scaffold N50 | 53.80 Mb | 43.82 Mb | 42.45 Mb |
| Scaffold L50 | 5 | 6 | 6 |
| BUSCO completeness<sup>d</sup> | 98.80% | 98.80% | 99.20% |
| GC content | 39.99% | 39.46% | 39.44% |
| Repeat content | 44.50% | 49.06% | 48.34% |
| Number of genes | 36,376 | 39,849 | 37,942 |
| Annotation BUSCO completeness<sup>e</sup> | 97.20% | 94.6% | 95.8% |

<sup>a</sup> Coverage based on 650 Mb genome size for *E. grandis* and *E. urophylla*.
<sup>b</sup> BUSCO completeness scores of contig level assembly.
<sup>c</sup> Total number of bases scaffolded onto one of the eleven chromosomes.
<sup>d</sup> BUSCO completeness scores of all scaffolds (including unplaced scaffolds).
<sup>e</sup> BUSCO completeness of gene annotation

Next, we mapped the parental Illumina reads to the corresponding haplogenome to investigate whether the smaller than expected haplogenome assembly size might be due to unassembled genomic regions. We observed mapping rates of 98.73% and 99.10% (93.79% and 92.91% properly paired), respectively (Supplementary Table 3), indicating that it is unlikely that major genomic regions are missing in the haplogenome assemblies.

#### Genome scaffolding

To curate incorrectly assembled contigs, contig breakpoints were inferred based on long-read depth support and used to split suspicious contigs before scaffolding. The parental genetic linkage maps yielded a set of 3,125 (for the *E. urophylla* haplogenome) and 3,129 (*E. grandis* haplogenome) unique SNP markers to anchor contigs into pseudo-chromosome level scaffolds. The anchoring rate for both haplogenome assemblies was greater than 88.0% (Table 2) and a BUSCO completeness score of at least 96.3% was obtained for contigs anchored to one of the eleven chromosomes. Dot-plot visualization of the haplogenome alignment confirmed high levels of collinearity between the assembled haplogenomes (Supplementary Figure 3 **and** Supplementary Figure 4). ALLMAPS was able to orientate 299 *E. urophylla* and 262 *E. grandis* contigs with two or more markers each., while 52 contigs for *E. urophylla* and 49 for *E. grandis* only had one marker and were placed without orientation (Table 2). A total of 1,067 contigs (corresponding to 63.37 Mb) of the *E. urophylla* and 1,268 contigs (67.78 Mb) of the *E. grandis* haplogenome assembly could not be anchored (Table 2) of which 863 (9.71 Mb) and 1,051 contigs (11.86 Mb) were smaller than 50 kb (Supplementary Table 4) and contained no markers.

**Table 2.** Summary statistics for parental linkage maps (gra_allmap and uro_allmap) and final consensus anchoring of the *E. urophylla* and *E. grandis* haplogenome contigs. A greater weight (indicated with w) was given to the linkage map of the species corresponding to the haplogenome being scaffolded. Scaffolds that contain no SNP markers or had ambiguous placements were counted as unplaced. Marker density (measured as number of markers per Mb) represents the sum of unique markers in the two linkage maps.

| *E. urophylla* | gra_allmap (w=1) | uro_allmap (w=2) | Anchored | Unplaced |
| --- | --- | --- | --- | --- |
| Linkage Groups | 11 | 11 | 11 | n.a. |
| Markers (unique) | 1,577 | 1,573 | 3,125 | 25 |
| Average markers per Mb | 3.5 | 3.5 | 6.5 | 0.4 |
| N50 Scaffolds | 76 | 79 | 81 | 2 |
| Scaffolds | 311 | 299 | 351 | 1,067 |
| Scaffolds with 1 marker | 83 | 80 | 52 | 13 |
| Scaffolds with 2 markers | 51 | 53 | 42 | 4 |
| Scaffolds with 3 markers | 41 | 37 | 44 | 0 |
| Scaffolds with ≥4 markers | 136 | 129 | 213 | 1 |
| **Total bases** | **448,984,013** | **447,297,011** | **481,132,251** | **63,374,165** |
| Percent of genome | 82.5% | 82.1% | 88.4% | 11.6% |

| *E. grandis* | gra_allmap (w=2) | uro_allmap (w=1) | Anchored | Unplaced |
| --- | --- | --- | --- | --- |
| Linkage groups | 11 | 11 | 11 | n.a. |
| Markers (unique) | 1,588 | 1,575 | 3,129 | 34 |
| Average markers per Mb | 3.3 | 3.4 | 6.3 | 0.5 |
| N50 Scaffolds | 72 | 72 | 73 | 1 |
| Scaffolds | 283 | 263 | 311 | 1,268 |
| Scaffolds with 1 marker | 62 | 60 | 49 | 21 |
| Scaffolds with 2 markers | 46 | 33 | 26 | 3 |
| Scaffolds with 3 markers | 32 | 32 | 30 | 1 |
| Scaffolds with ≥4 markers | 143 | 138 | 206 | 1 |
| **Total bases** | **477,075,775** | **464,179,728** | **498,948,047** | **67,775,781** |
| Percent of genome | 84.2% | 81.9% | 88.0% | 12.0% |

Most of the anchored assembly had a high level of congruence between the genetic and physical maps as indicated by the Pearson’s correlation coefficient (ρ) being close to −1 or 1, with the weakest correlation being ρ = 0.965 (Supplementary Figure 5) for *E. urophylla* and ρ = 0.938 for *E. grandis* (Supplementary Figure 6). Chromosome 3 and 5 differed from the *E. grandis* v2.0 reference genome by more than 20 Mb (Supplementary Figure 7), which could not be explained by a single missing genomic segment (Supplementary Figure 3). To investigate this, we aligned all unplaced scaffolds to the *E. grandis* v2.0 reference genome but did not observe any chromosomal preference for unplaced scaffolds (Supplementary Figure 8). This suggested that the chromosomal size differences were not due to scaffolds not being placed to those chromosomes (Supplementary Figure 8).

### Genome annotation

To further examine whether the smaller haplogenome assembly size is due to a difference in repeat content, we annotated repeat elements with RepeatMasker. A total of 48.34% of the *E. urophylla* haplogenome assembly comprised of repetitive elements, whereas it was 49.09% for the *E. grandis* haplogenome (Supplementary Table 5). In both cases, LTR retrotransposons were the most prevalent repetitive element, making up more than 21% of the assembled haplogenomes (Supplementary Table 5). DNA transposons made up ∼6% of the haplogenomes. These results are similar to previous repeat annotations for the v2.0 *E. grandis* reference assembly (Myburg *et al*., 2014, Table 1). We used LTR retriever to visualize the distribution of various LTR retrotransposon types (in bins of 300 kb Figure 2). LTR retriever, which is more sensitive for detection of LTR retrotransposons than RepeatModeler, identified 29.08% and 29.25% of the *E. grandis* and *E. urophylla* haplogenomes respectively, as LTR retrotransposons. Direct comparison of the LTR retrotransposon distribution pattern between *E. grandis* and *E. urophylla* was not possible as the assembled chromosomes differ in size, but there was good relative conservation in pattern with few notable exceptions e.g., on Chromosome 2 (Figure 2).

**Figure 2.**
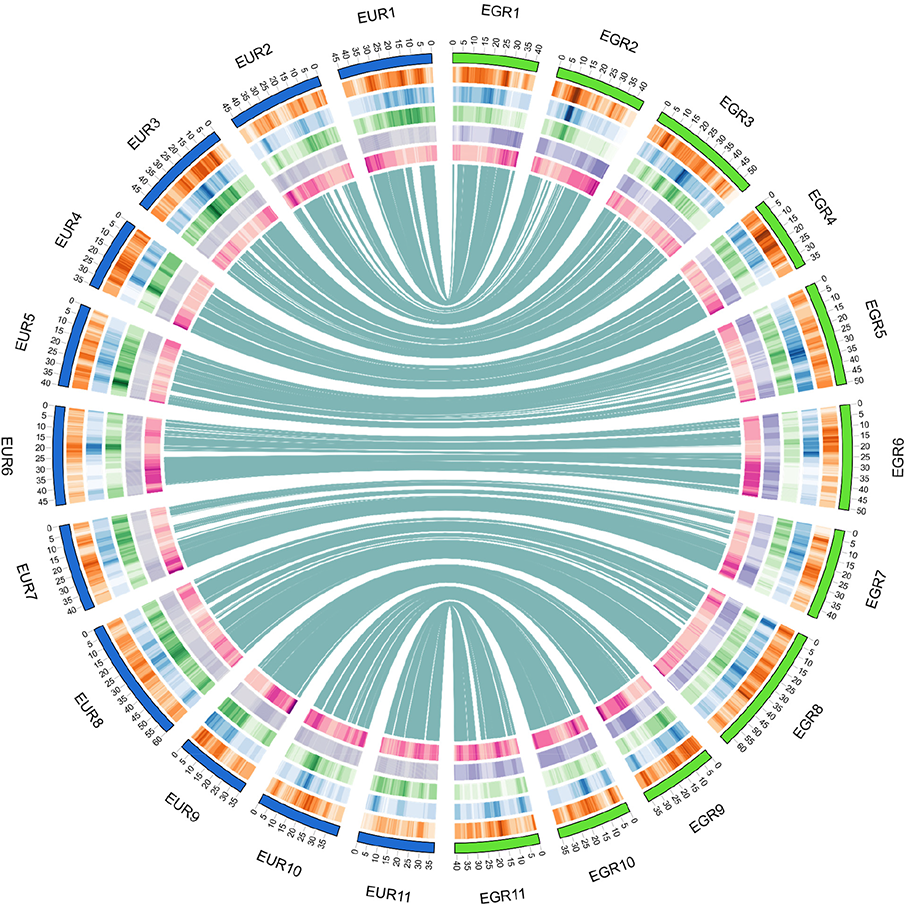
Synteny and distribution of LTR retrotransposons along the *E. grandis* and *E. urophylla* haplogenome assemblies for eleven scaffolded chromosomes. Syntenic regions are shown between the *E. urophylla* and *E. grandis* haplogenomes in the middle, based on SyRI (see Supplementary Figure 9). LTR retrotransposon distribution is shown for the *E. urophylla* (EUR) and the *E. grandis* (EGR) haplogenome assemblies. From outside to inside, the heatmaps show the distribution of Copia (orange, ranging from 6 to 21.5%), Gypsy (blue, ranging from 1.3 to 26.5%) and unknown (green, ranging from 2.8 to 16.6%) LTR retrotransposons, GC% (37.0 to 43.0%) and gene density (0 to 60.0%) with darker shades representing a higher percentage of retrotransposons within the bin. Chromosome number and size is indicated on the outer circle in megabases.

Structural (*de novo*) annotation resulted in 39,849 and 37,942 gene models for the *E. grandis* and *E. urophylla* haplogenomes, respectively (Table 1 and Supplementary Table 6). BUSCO completeness scores of 94.6% and 95.8% was obtained for the *E. grandis* and *E. urophylla* structural annotation models (Table 1). Functional annotation based on similarity searches or gene family assignment was possible for 35,572 and 33,915 structural gene models of *E. grandis* and *E. urophylla* (Supplementary Table 6).

### Structural variant analysis

*E. grandis* and *E. urophylla* are in the same section (*Latoangulatae*) and subgenus *Symphyomyrtus* but have non-overlapping natural ranges with unique adaptations such as greater resistance to fungal pathogens in *E. urophylla,* which has a more tropical distribution. Genetic linkage mapping has suggested high collinearity of their genomes (Hudson *et al*., 2011, Kullan *et al*., 2011, Bartholome *et al*., 2015), but a direct fine-scale comparison of genome synteny between these species has not been possible. Using the SyRI the whole-genome comparison tool, we revealed that a total of 257 Mb was syntenic between the two haplogenome assemblies, while 262.22 and 374.85 Mb were identified as rearranged in the *E. grandis* and *E. urophylla* haplogenomes, respectively (Figure 2, Figure 3, Supplementary Table 7, Supplementary Figure 9). In comparison, 318 Mb was syntenic between the *E. grandis* haplogenome and the *E. grandis* v2.0 reference genome (Supplementary Table 7, Supplementary Figure 9), but due to the difference in overall assembly size and methods used in the two studies, it is not possible to compare the genomic proportions. The regions rearranged between the haplogenomes included 189 inversions and 10,526 translocations (Figure 3, Supplementary Figure 9, Supplementary Table 7, Supplementary Table 9 **and** Supplementary Table 10). In addition, there were 16,865 duplications in the *E. grandis* and 21,149 duplications in the *E. urophylla* haplogenome (Figure 3, Supplementary Figure 9 **and** Supplementary Table 7). Together these results suggest that despite high collinearity previously reported for these species and observed here for the *E. grandis* and *E. urophylla* haplogenomes, extensive fine-scale rearrangements exist that have not been detected in previous studies.

**Figure 3.**
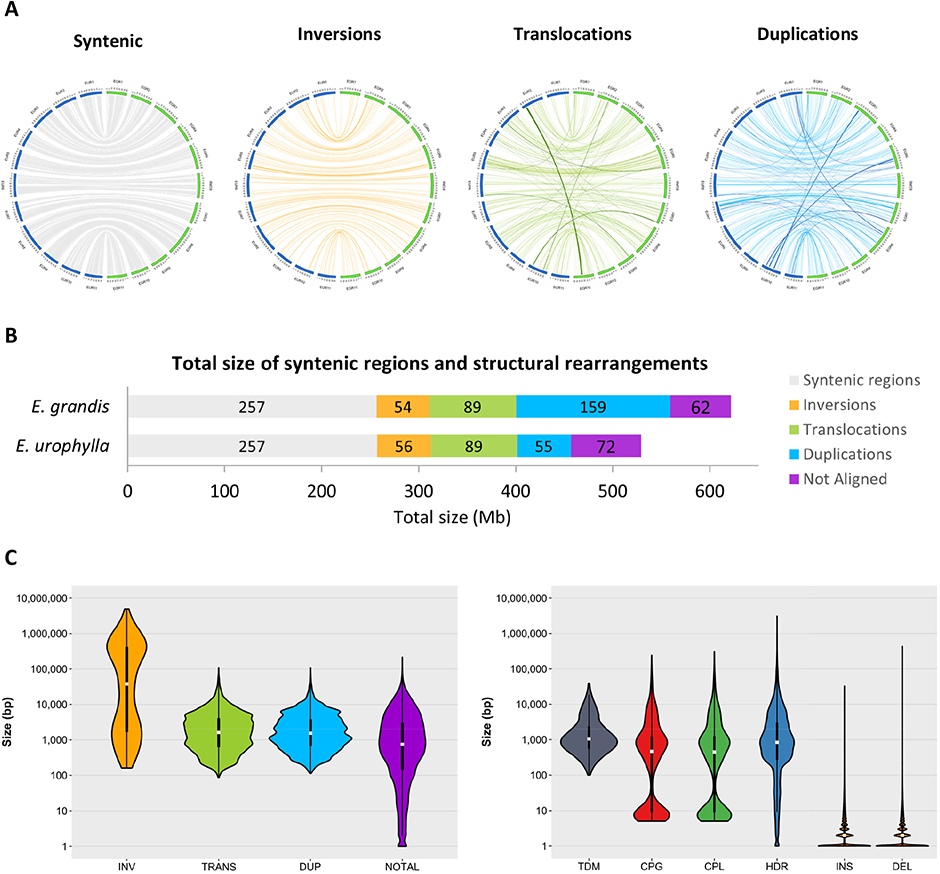
Size and distribution of structural rearrangements and local variants between the *E. grandis* and *E. urophylla* haplogenomes. (A) Distribution of syntenic regions and structural variants between the *E. grandis* and *E. urophylla* haplogenome assemblies. Links are shown between *E. urophylla* (EUR) chromosomes in blue and and *E. grandis* (EGR) chromosomes in green. Only variants of greater than 10 kilobases as identified by SyRI are shown. Darker links in show change in chromosome number between EUR and EGR. (B) Total size of syntenic and rearranged regions in megabases (Mb) for the *E. grandis* and *E. urophylla* haplogenome (see Supplementary Table 7 and Supplementary Table 8). The size of syntenic or rearranged regions are indicated within the bar in Mb, while the bar colour represents the rearrangement type. (C) Size distribution of rearranged regions (left) and local variants (right) between the *E. grandis* and *E. urophylla* haplogenomes. Size is indicated in base pairs on the y-axis (ranging from one to 4.91 Mb for rearrangements and one to 3.09 Mb for local variants), and the rearrangement type on the x-axis; INV are inversions, DUP are duplications, TRANS are translocations, NOTAL are regions that are not aligned, TDM are tandem repeats, CPG and CPL are copy gains/losses, HDR are highly diverged regions, INS are insertions and DEL are deletions.

Next, we investigated genome sequence divergence in syntenic regions, designated as “local variants” by SyRI, comprising 65.26 Mb and 66.45 Mb in the *E. grandis* and *E. urophylla* haplogenomes, respectively. These local variants (excluding SNPs) ranged from 1 bp (indels) to 3.09 Mb (highly diverged regions, HDR, Figure 3C). SNPs were the most prevalent class of local variants in terms of number, with 8.3 million SNPs between the *E. grandis* and *E. urophylla* haplogenomes, followed by small insertions and deletions (Supplementary Table 8). In terms of the total bases affected, highly diverged regions and copy gain/losses made up 9.6 Mb and 38 – 40 Mb of the haplogenome assemblies. Although there is a greater number of local variants compared to SVs, local variants made up 13.8% of the *E. urophylla* and 13.1% of the *E. grandis* chromosomal assembly compared to 54.5% and 75.1% in SV. This suggests that although local variants are more numerous, structural variants have a larger impact on genome architecture. This was also revealed in similar studies in tomato (Alonge *et al*., 2020) and grape (Zhou *et al*., 2019).

We performed gene-based synteny analysis between the *E. grandis* and *E. urophylla* haplogenomes, which confirmed high collinearity between the haplogenomes, with 23,390 gene pairs in 238 syntenic blocks (average 98.28 gene pairs per syntenic block with min = 4 gene pairs and max = 1296 gene pairs, Figure 4). A total of 227 blocks had 10 or more homologous gene pairs and 175 blocks had 30 or more gene pairs derived from the two haplogenomes. Of the 227 blocks, 86 blocks (8,114 genes and 37.88% of gene synteny blocks) are rearranged between the haplogenomes as inversions or translocations. The top GO enriched terms within these blocks belonged to regulation of transcription, DNA binding transcription factor activity and mRNA binding (a full list of enriched GO terms can be found in Supplementary Figure 10).

**Figure 4.**
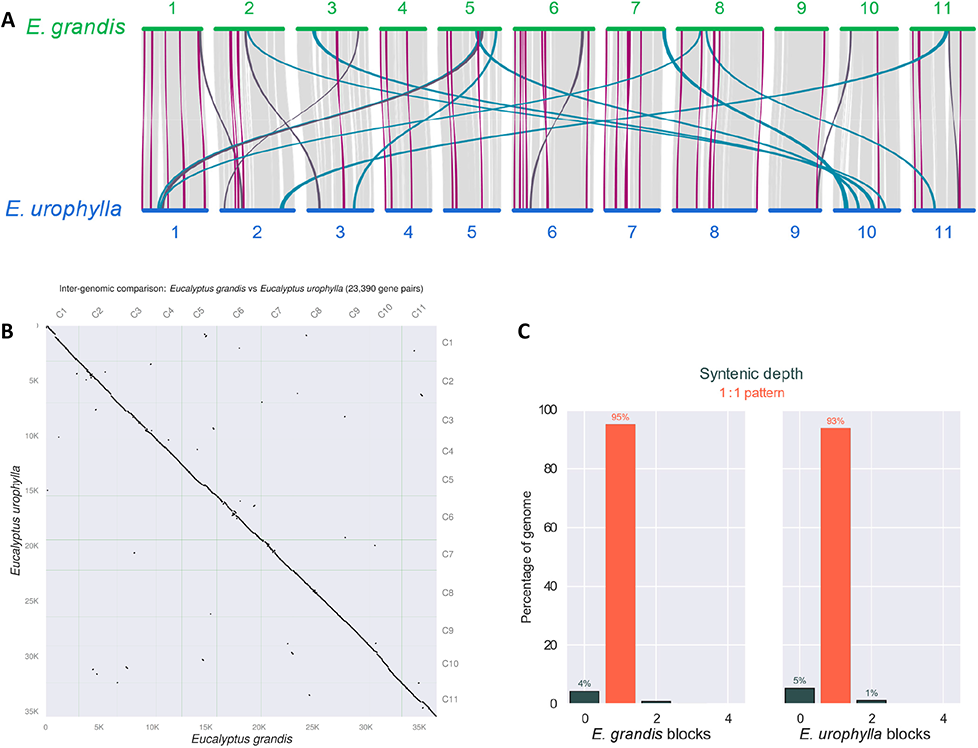
Gene synteny between *E. grandis* and *E. urophylla* haplogenome assemblies. (A) Chromosome-scale collinearity between *E. grandis* and *E. urophylla* haplogenome annotations. Lines in light grey indicate syntenic gene blocks, lines in purple indicate inverted gene blocks, blue indicated translocated gene block and dark grey inverted translocated gene blocks. Only block that span greater than 30 gene pairs are shown. (B) Dot-plot alignments of 23,390 gene pairs between the *E. grandis* and *E. urophylla* haplogenome annotations. (C) Bar graph showing syntenic depth of *E. grandis* and *E. urophylla* syntenic blocks. The majority of genes are in a 1 to 1 synteny pattern. All of the graphs were produced in MCScan JCVI v1.1.18 (Tang *et al*., 2008).

## Discussion

We have assessed the use of a trio-binning strategy to assemble high-quality haplogenomes in an F1 hybrid of two important eucalypt tree species as a starting point towards investigating pan-genome variation within and between these species. The high level of heterozygosity in the F_1_ hybrid enabled discrimination of almost all parental long reads and independent assembly of the two parental haplogenomes. These haploid assemblies, the first of their kind for a forest tree species, allowed us to circumvent the problem of co-assembly of alternative haplotypes which has presented a challenge for the assembly of highly heterozygous tree genomes, especially in intergenic DNA where complex structural variants from partially overlapping haplotypes may be co-assembled into a mosaic sequence (Myburg *et al*., 2014, Bartholome *et al*., 2015). Furthermore, the high coverage of long reads (50X per haplogenome) and the long-read length (N50 >27 Kb) allowed us to assemble across complex repeat structures leading overall to highly contiguous assemblies (contig N_50_ of 2.42 Mb for *E. grandis* and 1.93 Mb for *E. urophylla*). Intriguingly, we find that, despite having very high BUSCO completeness scores (>98%), the assembled haplogenomes (567 Mbp and 545 Mbp) were substantially smaller than the previous diploid reference genome assembly of 691.4 Mb (Myburg *et al*., 2014, Bartholome *et al*., 2015) and the ∼640 Mb flow cytometry estimate (Grattapaglia and Bradshaw Jr, 1994). High-density SNP genetic linkage maps enabled further improvement of haplogenome assembly contiguity (scaffold N_50_ >42 Mbp). Finally, we performed the first fine-scale structural and gene-based comparison for any two eucalypt genomes and show that SVs are more prevalent than detected in previous studies, but follow a similar class distribution pattern as in other plants with inversion events the least frequent, followed by translocation events and duplications being the most frequent (Goel *et al*., 2019, Jiao and Schneeberger, 2020).

### Trio-binning of a highly heterozygous F_1_ hybrid genome

The trio-binning strategy (Koren *et al*., 2018) allowed successful discrimination of the long reads derived from the *E. urophylla* and *E. grandis* haplogenomes. A total of 99.98% of the sequenced read bases could be assigned to one of the two haplo-bins, with only a small proportion (0.014%) of mostly shorter nanopore reads was not assigned to bins (N_50_ = 1,385 bp for un-binned vs N_50_ ∼ 27.5 kb for binned reads). The long-read data was split 51.80% vs 48.18% for *E. grandis* and *E. urophylla,* respectively, Supplementary Table 3), matching the assembly sizes, but it is not clear whether this can be generalized for individuals of the two species. Stringent cross-mapping of the parental short-read data to the two haplogenomes revealed, as expected, lower mapping rates to the opposite haplogenome (average 93.35% vs 84.94%, Supplementary Table 3) supporting that we have efficiently separated the haplogenome reads from the two species. The low level of BUSCO duplication in the assembled haplogenomes (less than 3.8%; Supplementary Figure 2) compared to 13.9% reported for a recent diploid *E. pauciflora* assembly (Wang *et al*., 2020a), supports that the haplotype binning was highly efficient. We further validated the size of phased blocks, as well as phase origin (**Supplementary Note 1**) and found that the haplogenome assemblies had very low haplotype switch error rates (lower than 0.033%) confirming the accuracy of haplotype separation. Together these results suggest that the trio-binning approach was highly efficient and accurate in the heterozygous F_1_ hybrid genome.

Haplotype separation is known to improve with higher levels of heterozygosity (Koren *et al*., 2018, Rhie *et al*., 2020). We observed high heterozygosity for both pure-species parents (2.14% for *E. grandis* and 2.63% for *E. urophylla*), and as expected, heterozygosity was substantially higher in the F_1_ hybrid offspring (estimated to be 3.46%; Supplementary Figure 1). Such high heterozygosity levels are expected for outcrossed organisms such as eucalypts (Moran *et al*., 1989, Gaiotto *et al*., 1997). Successful haplotype separation of an F_1_ hybrid of species within the same section of Myrtaceae (*Latoangulatae*) suggests that application of trio-binning for haplotype separation should be successful for most other viable *Eucalyptus* F_1_ hybrid combinations. In addition, the high heterozygosity observed in the pure species parents suggests that haplotype binning will also be successful in intraspecific crosses of *Eucalyptus* as the trio-binning strategy has been demonstrated to be efficient at much lower levels of heterozygosity (0.9% in the case of a F_1_ Brahman x Angus cattle hybrid and 1.36% for *A. thaliana*; Koren *et al*., 2018).

We note that the haplogenome assembly sizes, 546/481 Mb for *E. urophylla* and 568/498 Mb for *E. grandis* (total/scaffolded size) were much smaller than that of the current *E. grandis* v2.0 reference genome (691/612 Mb, Myburg *et al*., 2014; Bartholome *et al*., 2015) and previous estimates (∼ 640 Mb) based on flow cytometry (Grattapaglia and Bradshaw Jr., 1994). K-mer based genome size estimates of the parental reads predicted diploid genome sizes of 443 Mb for *E. urophylla,* 482 Mb for *E. grandis* and 478 Mb for the F_1_ hybrid (Supplementary Figure 1), which agreed with the scaffolded genome sizes of the two haplogenome assemblies. This apparent discrepancy was also observed in *E. pauciflora*, where k-mer based estimates were 408 Mb compared to the final 595 Mb assembly (Wang *et al*., 2020a). The total assembly sizes of the two haplogenomes were therefore approximately 70 - 100 Mb smaller than previous flow cytometry estimates for the two species and the total scaffolded sizes were 140 - 160 Mbp smaller than expected. This size discrepancy may be explained by several factors, which we explore below.

First, to exclude the possibility that the smaller assembly size was due to a portion of sequencing reads not being assembled, i.e., that we failed to assemble parts of the haplogenomes, we aligned the parental Illumina reads to the corresponding parental haplogenome assembly. We also aligned the raw short- and long-reads and the haplogenome assemblies to the *E. grandis* v2.0 reference genome to make sure all v2.0 genomic regions had sequencing coverage (**Supplementary Note 2**). This revealed that some regions had very high sequencing depth relative to the *E. grandis* v2.0 reference genome (**Supplementary Note 2**) presumably due to highly repetitive sequence content in those regions. More than 98.7% of parental Illumina reads aligned to their corresponding parental haplogenome, which suggests that almost all of the sequences in the parental genomes (that are amenable to Illumina sequencing) are represented in the haplogenomes (Supplementary Table 2), although it is possible that the regions with high sequencing depth represent repetitive regions that are collapsed in the haplogenome assemblies. To further investigate this possibility, we confirmed that the repeat content of the haplogenomes was not lower than that reported in the *E. grandis* v2.0 diploid reference assembly. In fact, the repeat content for the *E. urophylla* and *E. grandis* haplogenomes (48.16% and 48.91%, respectively, Table 1) was higher than that reported for the *E. grandis* v2.0 assembly (44.50%, Myburg *et al*., 2014) and for the more recent *E. pauciflora* assembly (44.77%, Wang *et al*., 2020a). This suggests that the observed size difference is most probably not due to the collapse of repetitive regions during haplogenome assembly. Rather, the slightly higher repeat content of our haplogenome assemblies probably reflect our ability to better assemble across such repeats using long-read technology in haplo-assemblies vs short-read/Sanger sequencing previously used for these highly heterozygous genomes. Previous size estimates were probably somewhat inflated in size due to the possible co-assembly of partially overlapping alternative haplotypes highly heterozygous regions distributed throughout the genome. Our analysis showed that Chromosomes 3 and 5 in the haplogenome assemblies were 20 Mb smaller than the corresponding chromosomes in the diploid *E. grandis* v2.0 assembly.

### Genetic linkage maps support high scaffolding rates

Overall, 88.4% and 88.0% of the haplogenome assemblies was anchored into 11 pseudo-chromosomes for *E. urophylla* and *E. grandis*. However, there are some limits to using ALLMAPS for genome scaffolding as the program cannot identify and separate duplicated regions that are misassembled or collapsed by the genome assembler due to high similarity (Tang *et al*., 2015). In addition, most genetic linkage maps contain regions such as centromeres with no or very low recombination and few DNA markers for anchoring and orientation of contigs. Many of the unanchored contigs may contain difficult to assemble, centromeric or other non-recombinogenic regions devoid of mapped DNA markers (average 0.4 and 0.5 markers per Mb for unanchored vs 6.5 and 6.3 markers per Mb for anchored *E. urophylla* and *E. grandis* contigs, respectively, Table 2). The N_50_ of the unanchored contigs was 324 kb, which was smaller than the average marker spacing in those regions (Supplementary Table 4). Thus, integration of additional proximity ligation or optical mapping data may lead to inclusion of some of the remaining unplaced contigs that had few markers to place or orient them. Despite this limitation we were able to produce eleven pseudo-chromosome scaffolds for each of the haplogenomes owing to the high density of SNP markers in the parental maps and the quality of the genetic maps as evidenced in collinearity of markers between the genetic map and the *de novo* assembled contigs, as well as high collinearity between the scaffolded assembly and the genetic linkage maps (Pearson’s correlation of ρ = 0.938 to ρ = 1.00; Supplementary Figure 5 **and** Supplementary Figure 6).

### Structural variants between *E. urophylla* and *E. grandis*

To our knowledge, this is the first genome-wide comparison of synteny and structural rearrangements between two eucalypt species. In addition, we had the advantage of being able to directly compare the two haplogenomes from the same F_1_ hybrid individual assembled using the same method. Using SyRI we found that 53.39% (256.88 Mb) of the 481.16 Mb chromosomal assembly of *E. urophylla* and 51.45% (256.75 Mb) of the 498.97 Mb chromosomal assembly of *E. grandis* was syntenic (Supplementary Table 7). We were able to identify 48,729 SVs between the two haplogenomes, with a 103.62 Mb difference between the two haplogenomes due to duplications (Supplementary Table 7). As seen in previous studies using SyRI for SV calling, we found that inversions were the smallest group of SVs in terms of number, followed by translocations, with duplications being the most abundant (189 inversions, 10,526 translocations and 38,014 duplications, Supplementary Table 7; Goel *et al*., 2019; Jiao and Schneeberger, 2020). Using unfolded site frequency spectrum of SVs, Zhou *et al*., (2019) found that there is purifying selection against SVs, and that there is stronger purifying selection against inversions and translocations compared to duplications as they have a more deleterious effect compared to duplications (Zhou *et al*., 2019). Stronger purifying selection against inversions and translocations in our haplogenome assemblies may therefore explain the lower frequency of these two classes of SV, however this will need to be tested in future sequencing projects including population-wide tracking of SVs.

With additional genome sequences for *E. grandis* and *E. urophylla*, a pan-genome reference assembly could be constructed as was done for *Arabidopsis* (Jiao and Schneeberger, 2020) and tomato (Alonge *et al*., 2020, Wang *et al*., 2020b). SyRI identifies SVs and local variants using three main steps: 1) identify syntenic alignments, 2) identify inverted, duplicated and translocated alignments and 3) identify “local variants” within alignment blocks. As such, there is a hierarchy of variation where local variants are found within alignment blocks, be they syntenic or rearranged regions. However, when looking for the functional effects of local and larger structural variants, it is important to note the hierarchy of genomic rearrangements, as local variants within rearranged regions show different inheritance patterns to those in syntenic regions. SVs can influence recombination studies as rearrangement hotspots typically have lower synteny and reduced recombination rates (Jiao and Schneeberger, 2020). In addition, SVs can influence gene expression directly or indirectly making their functional interpretation harder (Goel *et al*., 2019).

Surprisingly, despite the high completeness, we found that the total assembled size of each of the haplogenomes is substantially smaller than that of the *E. grandis* v2.0 reference genome and previous flow cytometry estimates. We propose that the size difference is not due to collapse of the repeat content of the haplogenome assemblies, but rather due to possible overestimation of the *E. grandis* v2.0 genome assembly as a result of inclusion of partially overlapping alternative haplotypes in highly heterozygous regions of the diploid genome assembly. However, resolving this discrepancy will require further *de novo* genome assemblies for *E. grandis*, possibly including resequencing using long read technology to update the genome assembly of the reference BRASUZ1 individual, as has been performed for some reference genomes that were originally assembled with Sanger sequencing data (Nurk *et al*., 2022).

Finally, we provide the first genome-wide SV between *E. urophylla* and *E. grandis*. We identify 48,729 SVs, ranging in size from 100 bp to 4.91 Mb. Some of these variants are large enough to be able to cover multiple genes and future studies will focus on understanding the genomic context and functional implications of these variants. An added advantage of this study is the reduced false discovery of SVs which may be introduced when comparing genomes assembled using different pipelines.

## Materials and Methods

### Sample background

Leaf tissues of an F_1_ *E. urophylla* x *E. grandis* hybrid offspring and its parents (*E. urophylla* seed parent and *E. grandis* pollen parent) were collected and used for DNA extractions. These individuals form part of a large nested association mapping trial and SNP data was used to generate high-density genetic linkage maps for both the *E. grandis* and *E. urophylla* parents (Candotti *et al*. unpublished). Sequencing both parents will enable a) inference of both haplotypes for the parental genomes and b) haplotype binning for genome phasing (Figure 1).

### DNA isolation

#### Illumina sequencing

Genomic DNA was extracted from 50 mg of leaf tissue for the *E. urophylla* and *E. grandis* parents using the NucleoSpin® Plant II Kit (Machery-Nagel, Germany). Gel electrophoresis was performed using a 0.8% w/v agarose gel to assess DNA quality. DNA quality was also assessed using a NanoDrop® ND-1000 spectrophotometer (Thermo Fisher Scientific) and quantified using a Qubit 2.0 Fluorometer (Thermo Fisher Scientific). Whole-genome sequencing of the F_1_ hybrid and its parents was performed on an Illumina NovaSeq6000 platform by Macrogen (Macrogen Inc., Seoul, Korea).

#### High molecular weight DNA extraction

Genomic DNA was extracted using 1.2 g of flash frozen ground leaf tissue. The ground material was suspended in 25 ml Guanidine buffer (20 mM EDTA, 100 mM NaCl, 1% Trition® X-100, 500 mM Guanidine-HCl and 10 mM Tris, pH 7.9), supplemented with 50 mg cellulase (Sigma-Aldrich) and 50 mg lysing enzyme (Sigma-Aldrich) incubated at 42 °C with gentle agitation. After 2.5 h, 10 µl RNase A (20 µg/ml) was added and the sample was incubated for 30 min at 37 °C, after which 50 mg proteinase K was added, and the mixture was incubated for another 2 h at 50 °C. The mixture was then centrifuged for 20 min at 12 000 x *g* and the clarified lysate transferred to an appropriate buffer QBT-equilibrated QIAGEN Genomic-tip 100/G column (Qiagen), after which the column was washed three times with 7 ml Buffer QC and HMW DNA was eluted with 5 ml Buffer QF. The DNA was precipitated by adding 0,7 V of isopropanol and centrifuged at 12 000 *g* for 20 min. The DNA pellet was washed twice with 70% Ethanol and resuspended in an appropriate volume of low salt TE (10 mM Tris-HCL pH 8.0; 0.1 mM of EDTA). Gel electrophoresis was performed using a 0.8% w/v agarose gel to assess DNA quality, and DNA quantity was assessed using a Qubit 2.0 Fluorometer (Thermo Fisher Scientific).

#### Nanopore sequencing

HMW DNA was prepared for MinION sequencing following the manufacturers protocol using the genomic sequencing kit SQK-LSK109 (Oxford Nanopore Technologies, Oxford, UK). Approximately 3.3 µg of HMW DNA from was used without exogenous shearing or size selection. HMW DNA was first repaired with NEBNext FFPE Repair Mix (New England Biolabs) and 3’-adenylated with NEBNext Ultra II End Repair/dA-Tailing Module (NEB). The DNA was then purified with AMPure XP beads (Beckmann Coulter) and ligated with sequencing adapters (ONT) using NEBNext Quick T4 DNA Ligase (NEB). After purification with AMPure XP beads (Beckman Coulter), the library was mixed with sequencing buffer (ONT) and library loading beads (ONT) and loaded on primed MinION R9.4 SpotOn flow cells (FLO-MIN106). MinION sequencing was performed with a MinION Mk1B sequencer running for 48 h.

The resulting FAST5 files were base-called and reads with a QV < 7 were removed with Oxford Nanopore Technologies’ Guppy base-calling software v3.4.5 (ONT) using parameters for FLO-MIN106 and SQK-LSK109 library type. RStudio was used summarise and visualise statistics for both DNA isolation methods based on the sequencing summary file generated by the Guppy base-caller. The Guppy base-caller may not remove all the sequence adapters so to ensure all sequence adapters are removed PoreChop v0.2.4 (https://github.com/rrwick/Porechop) was used. All scripts used in this study is available online (https://gitlab.com/Anneri/eucalyptus-haplogenome-synteny). The resulting adapter-less reads of both DNA isolation methods were combined into a single FASTQ file for further use.

PromethION sequencing was performed by the Centre for Genome Innovation (University of Connecticut, Connecticut, USA) on a FLO-PRO002 PromethION flow cell as per the PromethION sequencing protocol (ONT) using the SQK-LSK109 (ONT) sequencing kit with Circulomics Short Read Eliminator XS (Circulomics Inc.) size-selection. The flow cell was washed and reloaded after 38 h and run for an additional 6 h of sequencing. Base-calling was performed using the Guppy v3.4.5 basecaller and adapter removal was performed as stated above.

### Genome assembly

#### Trio-binning and haplogenome assembly

Illumina short-reads were used for k-mer based genome size estimation was performed using Jellyfish v2.2.6 (Marcais and Kingsford, 2011) for 21-mers and visualised with GenomeScope v2.0 (Ranallo-Benavidez *et al*., 2020). Long-reads of the F_1_ hybrid were binned into *E. urophylla* and *E. grandis* haplotype bins (corresponding to parental short-reads) using the Trio-Canu module in Canu v1.8 (Koren *et al*., 2017). Read contaminants were identified from the binned reads using Centrifuge v1.0.4-beta (Kim *et al*., 2016) and removed with a custom script. Similarly, contaminant reads were also identified and removed from short read data with Kraken v2.0.8-beta (Wood *et al*., 2019). The remaining raw reads were used for all assembly and alignment steps.

The binned reads corresponding to each of the parents were assembled separately, along with the corresponding parental short reads, using the MaSuRCA v3.3.4 (Zimin *et al*., 2017) genome assembler. MaSuRCa was chosen as initial testing of multiple genome assemblers (based on the BUSCO completion score, contig N_50_ and total assembly size) indicated that the MaSuRCA genome assembler performed the best. Quality of the resulting assemblies was assessed using QUAST v5.0.2 (Gurevich *et al*., 2013, Mikheenko *et al*., 2018) and BUSCO v4.0.2 (Simao *et al*., 2015, Seppey *et al*., 2019). To verify genome coverage of the assemblies, Illumina reads from each of the parental haplotypes were mapped to the corresponding and alternative assembled haplogenomes using BWA v0.7.5a-r405 (Li and Durbin, 2009) and mapping rate calculated using the flagstat module from Samtools v1.9 (Li *et al*., 2009).

#### Genome scaffolding

To improve assembly contiguity, scaffolding was performed for the MaSuRCa assembled *E. urophylla* and *E. grandis* genomes using high-density genetic linkage maps previously constructed for each of the parents (Candotti *et al*., unpublished). To resolve possible chimeric contigs that were assembled by MaSuRCa, Polar_Star (https://github.com/phasegenomics/polar_star) was used to infer breakpoints and split in contigs based on identification of read-depth outliers from the binned long-reads. After breakpoints were inferred and contigs split, all contigs smaller than 3 kb were removed before scaffolding with high-density genetic linkage maps. A BLAST database was created for both assembled haplogenomes to identify the position of 1,588 *E. grandis* and 1,575 *E. urophylla* SNP probes used to construct the genetic maps. A consensus map was constructed with ALLMAPS (Tang *et al*., 2015), consisting of SNPs that mapped to the assembled haplogenomes, and was used to perform genome scaffolding. For the consensus map construction, a weight of two was given to the parental genetic linkage map corresponding to species haplogenome to be scaffolded, while a weight of one was given for the alternative parental linkage map. Chromosome scaffold sizes from the two haplogenomes were compared to one another and to the *E. grandis* v2.0 genome to see whether there is a size difference between the *E. grandis* v2.0 reference potential bias in scaffolding of particular chromosomes. To validate if unplaced contigs/scaffolds were from a particular chromosome, unplaced contigs/scaffolds were aligned to the *E. grandis* v2.0 genome using MiniMap2 (Li, 2016) and alignments visualized with D-Genies (Cabanettes and Klopp, 2018).

### Genome annotation

Custom libraries of repetitive elements were constructed for the *E. urophylla* and *E. grandis* haplogenomes with RepeatModeler v1.0.8 (http://www.repeatmasker.org/RepeatModeler/) (Smit and Hubley, 2008). Repetitive elements were annotated with RepeatMasker v4.0.9 (http://www.repeatmasker.org/) (Smit *et al*., 2013) for the haplogenome assemblies. To eliminate the chance of missing repeat elements in either haplogenome due to them not being identified, the combined species library was used as input for RepeatMasker. Lastly, to identify the abundance of LTR retrotransposons, LTR retrotransposon candidates were identified with LTR retriever (Ou and Jiang, 2018) for both haplogenomes and their distribution visualised with Circos (Krzywinski *et al*., 2009).

RNA-Seq reads from previous studies were used for structural genome annotation. RNA-Seq reads used for the *E. grandis* haplogenome assembly were from the original genome assembly paper and included six different tissues from an *E. grandis* individual (Mizrachi *et al*., 2010, Vining *et al*., 2015) all data is available on https://eucgenie.org/. RNA-Seq data from three-year-old *E. grandis* x *E. urophylla* backcrossed with *E. urophylla* trees were used and only derived from two tissues (mature leaf and xylem) Bioproject: https://www.ncbi.nlm.nih.gov/bioproject/PRJNA354497 (Mizrachi *et al*., 2017). RNA-Seq reads were trimmed with Trimmomatic v0.39 (Bolger *et al*., 2014) and only paired reads were used for further work. Trimmed RNA-Seq reads were aligned to the relevant haplogenome assemblies with Hisat2 v2.1.0 (Kim *et al*., 2019). GenomeThreader v1.7.1 (Gremme *et al*., 2005) was used to align protein sequences of the *E. grandis* v2.0 genome annotation to the haplogenome assemblies. We used BRAKER2 v2.0.5 (Hoff *et al*., 2016) for structural gene prediction. To predict protein coding regions in the genome, Braker2 first converts RNA-Seq alignments to exon support with GeneMark-ET v4.38 (Lomsadze *et al*., 2014). This output is combined with protein alignments for two rounds of training with AUGUSTUS v3.2.3 (Stanke *et al*., 2006, Stanke *et al*., 2008, Camacho *et al*., 2009). The predicted gene space was then filtered with gFACs v1.1.3 (Caballero and Wegrzyn, 2019). Mono-exonic genes were filtered with InterProScan v5.35-74.0 to keep only those with known protein domains. Completeness of the structural annotations were assessed with BUSCO v4.0.2.

Functional genome annotation was performed with EnTAP v0.9.0 (Hart *et al*., 2020) using the following public databases: NCBI RefSeq complete and EMBL-EBI UniProt. This pipeline integrates similarity search and other annotation resources which include gene family (eggNOG), protein domains (Pfam), gene ontology and KEGG pathway assignment.

### Structural variant identification

To check for regions that are unassembled in the haplogenome assemblies compared to the *E. grandis* v2.0 reference genome, the *E. grandis* and *E. urophylla* haplogenomes were each aligned to the *E. grandis* v2.0 genome, with MiniMap2 (Li, 2016) and alignments visualised using D-Genies (Cabanettes and Klopp, 2018). Using the same method, the eleven assembled *E. grandis* and *E. urophylla* chromosomes were aligned to each other to visually identify genomic regions with possible large structural variants (SVs). To identify structural rearrangements (inversions, translocations and duplications) and local variations (SNPs, InDels, copy gains/losses, highly diverged regions and tandem repeats) between *E. grandis* and *E. urophylla*, haplogenome assemblies were aligned to each other using nucmer from the MUMmer3 toolbox (Kurtz *et al*., 2004) with alignment parameters “— maxmatch --c 100 -b 500 -l 50”. The resulting alignments were further filtered for alignment length (>100) and identity (>90). Identification of structural rearrangements and local variations was performed using the Synteny and Rearrangement Identifier (SyRI) pipeline (Goel *et al*., 2019). The same method was also used to identify regions that differed between the *E. grandis* haplogenome and the *E. grandis* v2.0 reference genome. As the linear visualisation of syntenic regions and variants from SyRI prohibits us from depicting inter-chromosomal events, synteny and variants of greater than 10 kb were visualised with Circos.

Syntenic gene pairs between the *E. grandis* and *E. urophylla* haplogenomes were identified using a python version of MCScan, JCVI v1.1.18 (Tang *et al*., 2008). Coding sequence and annotation gff3 files were used as input data to identify the syntenic blocks for each pair of species with the ‘jcvi.compara.catalog ortholog’ command and a c-score parameter of --cscore=0.95. Syntenic blocks were filtered with ‘jcvi.compara.synteny screen’ with parameters --minspan=30 --simple. The pattern of synteny was detected with jcvi.compara.synteny depth –histogram. Smaller syntenic blocks were also filtered with ‘jcvi.compara.synteny screen’ with parameters --minspan=10 --simple. Genes within inverted and translocated syntenic blocks that spanned ten or more gene pairs were checked for gene ontology enrichment terms using Blast2GO v1.20.14 (Conesa *et al*., 2005) and results were visualized using Tableau Professional Edition (Tableau Software Inc., Seattle, WA, USA).

## Funding

This work was funded by the South African Department of Science and Innovation and Technology Innovation Agency (DSI/TIA, Strategic Grant for Eucalyptus Genomics), the Forestry Sector Innovation Fund (FSIF) and Sappi South Africa through the Forest Molecular Genetics (FMG) Industry Consortium at the University of Pretoria (UP). AL acknowledges MSc bursary support from the National Research Foundation (NRF) of South Africa (Grant Number MND190406427887) and funding from the UP Postgraduate Studies Abroad Programme.

## Acknowledgements

The authors thank the UP Bioinformatics and Computational Biology Centre and University of Connecticut (UConn) Computational Biology Core for bioinformatics and computational biology support. Sappi Forest Research (Howick, South Africa) kindly provided the plant materials used in the study.

## Author contributions

AL performed all analyses in the manuscript and prepared the manuscript. JC constructed genetic linkage maps for genome scaffolding. TD and JLW provided bioinformatic and technical support and advised on data analysis and interpretation throughout the project. TD, EM and JLW co-supervised the project. AAM conceived and supervised the project. All authors read and contributed to the final manuscript.

## Conflict of interest

The authors declare no conflict of interest.

## Supplementary Tables

**Supplementary Table 1.** Illumina sequencing results. Raw read statistics are given followed by the mapping rate of reads to the main scaffolds of the *E. grandis* v2.0 reference genome (Myburg *et al*., 2014) as well as the *E. grandis* mitochondrial and plastid genomes (Pinard *et al*., 2019) after read contaminants have been removed. The total amount of sequencing data generated is given in gigabases (Gb).

| Sample ID | Total bases (Gb) | Total reads | Q20 (%) | Q30 (%) | Mapping rate<sup>a</sup> (%) | Properly paired<sup>b</sup> (%) |
| --- | --- | --- | --- | --- | --- | --- |
| *E. urophylla* parent | 127.5 | 884,340,104 | 97.097 | 92.515 | 94.34 | 82.61 |
| *E. grandis* parent | 141.6 | 937,939,296 | 96.357 | 90.786 | 95.07 | 86.55 |
| F<sub>1</sub> hybrid | 116.1 | 769,097,570 | 97.361 | 93.182 | 94.78 | 84.16 |

<sup>a</sup> Mapping rate of Illumina short reads with contaminants removed to the *E. grandis* v2.0 main scaffolds along with the mitochondrial and plastid reference genomes.
<sup>b</sup> Mapping rate of Illumina short reads with contaminants removed to *E. grandis* v2.0 main scaffolds along with the mitochondrial and plastid reference genomes that are properly paired reads.

**Supplementary Table 2.** Nanopore sequencing results for the F_1_ hybrid individual. kb – kilobase, Gb – gigabase, Q7 – quality score of seven.

| Sequencing run | Number of reads | Total bases (Gb) | Number reads > Q7<sup>a</sup> | Total bases > Q7 (Gb)<sup>b</sup> | Longest read > Q7 (kb) | Read N50 (kb) |
| --- | --- | --- | --- | --- | --- | --- |
| MinION run 1 | 2,169,269 | 11.18 | 1,645,369 (75.85%) | 9.30 (83.18%) | 182.95 | 18.96 |
| MinION run 2 | 429,264 | 2.55 | 364,541 (84.92%) | 2.28 (89.41%) | 304.89 | 23.86 |
| PromethION | 3,290,284 | 61.59 | 2,875,796 (87.40%) | 56.57 (91.85%) | 221.38 | 28.00 |
| **Total<sup>c</sup>** | **5,888,817** | **75.32** | **4,885,706 (82.97%)** | **68.15 (90.48%)** | **304.89** |  |

<sup>a</sup> Percentage of reads passing QC are shown in brackets.
<sup>b</sup> Percentage bases passing QC are shown in brackets.
<sup>c</sup> 99.45% of basecalled reads mapped to the *E. grandis* v2.0 main scaffolds along with the mitochondrial and plastid reference genomes, with 99.94% mapping of the *E. urophylla* read bin and 99.93% of the *E. grandis* read bin.

**Supplementary Table 3.** Summary statistics for long-read binning using the parental short reads.

| **Bin** | **Reads<sup>a</sup>** | **L50** | **N50 (bp)** | **Max read length** | **Sum** | **Percentage of total** | **Mapping rate<sup>b</sup>** | **Properly paired<sup>c</sup>** | **Mapping rate<sup>d</sup>** | **Properly paired<sup>e</sup>** |
| --- | --- | --- | --- | --- | --- | --- | --- | --- | --- | --- |
| *E. grandis* | 1,999,561 | 465,851 | 27,604 | 262,889 | 35.11 Gb | 51.80% | 98.73% | 93.79% | 98.11% | 85.03% |
| *E. urophylla* | 1,877,139 | 433,849 | 27,548 | 304,871 | 32.66 Gb | 48.18% | 99.10% | 92.91% | 97.67% | 84.85% |
| Unknown | 6,693 | 2,553 | 1,385 | 7,631 | 9.59 Mb | 0.014% | NA | NA | NA | NA |
| **Total** | **3,883,393** |  |  |  | **67.78 Gb** |  |  |  |  |  |

<sup>a</sup> Only reads greater than 500 base pairs were considered.
<sup>b</sup> Mapping rate of corresponding parental species Illumina short reads to haplome assembly.
<sup>c</sup> Mapping rate of parental species Illumina short reads to haplome assembly that are properly paired reads.
<sup>d</sup> Mapping rate of alternative parental species Illumina short reads to haplome assembly.
<sup>e</sup> Mapping rate of alternative parental species Illumina short reads to haplome assembly that are properly paired reads.

**Supplementary Table 4.** Summary statistics of placed and unplaced contigs after scaffolding with ALLMAPS for the *E. urophylla* and *E. grandis* haplogenomes respectively. # indicates number.

| Assembly | Unplaced *E. urophylla* | Placed *E. urophylla* | Unplaced *E. grandis* | Placed *E. grandis* |
| --- | --- | --- | --- | --- |
| # contigs ($\geq 0$ bp) | 1,067 | 11 | 1,268 | 11 |
| # contigs ($\geq 1000$ bp) | 1,067 | 11 | 1,268 | 11 |
| # contigs ($\geq 5000$ bp) | 796 | 11 | 968 | 11 |
| # contigs ($\geq 10000$ bp) | 519 | 11 | 599 | 11 |
| # contigs ($\geq 25000$ bp) | 308 | 11 | 325 | 11 |
| # contigs ($\geq 50000$ bp) | 204 | 11 | 217 | 11 |
| Total length ($\geq 0$ bp) | 63,374,165 | 481,166,251 | 67,775,781 | 498,977,947 |
| Total length ($\geq 1000$ bp) | 63,374,165 | 481,166,251 | 67,775,781 | 498,977,947 |
| Total length ($\geq 5000$ bp) | 62,333,808 | 481,166,251 | 66,626,180 | 498,977,947 |
| Total length ($\geq 10000$ bp) | 60,386,045 | 481,166,251 | 64,045,556 | 498,977,947 |
| Total length ($\geq 25000$ bp) | 57,251,799 | 481,166,251 | 59,766,617 | 498,977,947 |
| Total length ($\geq 50000$ bp) | 53,663,936 | 481,166,251 | 55,918,898 | 498,977,947 |
| # contigs | 1,067 | 11 | 1,268 | 11 |
| Largest contig | 2,720,265 | 60,186,531 | 2,440,300 | 63,773,828 |
| Total length | 63,374,165 | 481,166,251 | 67,775,781 | 498,977,947 |
| GC (%) | 39.5 | 39 | 39.52 | 39 |
| N50 (bp) | 324,500 | 45,562,418 | 324,100 | 44,251,077 |
| N75 (bp) | 118,300 | 40,242,915 | 96,704 | 40,936,616 |
| L50 | 46 | 5 | 53 | 5 |
| L75 | 127 | 8 | 141 | 8 |
| # N's per 100 kbp | 0.78 | 7.3 | 1.64 | 6.31 |

**Supplementary Table 5.**
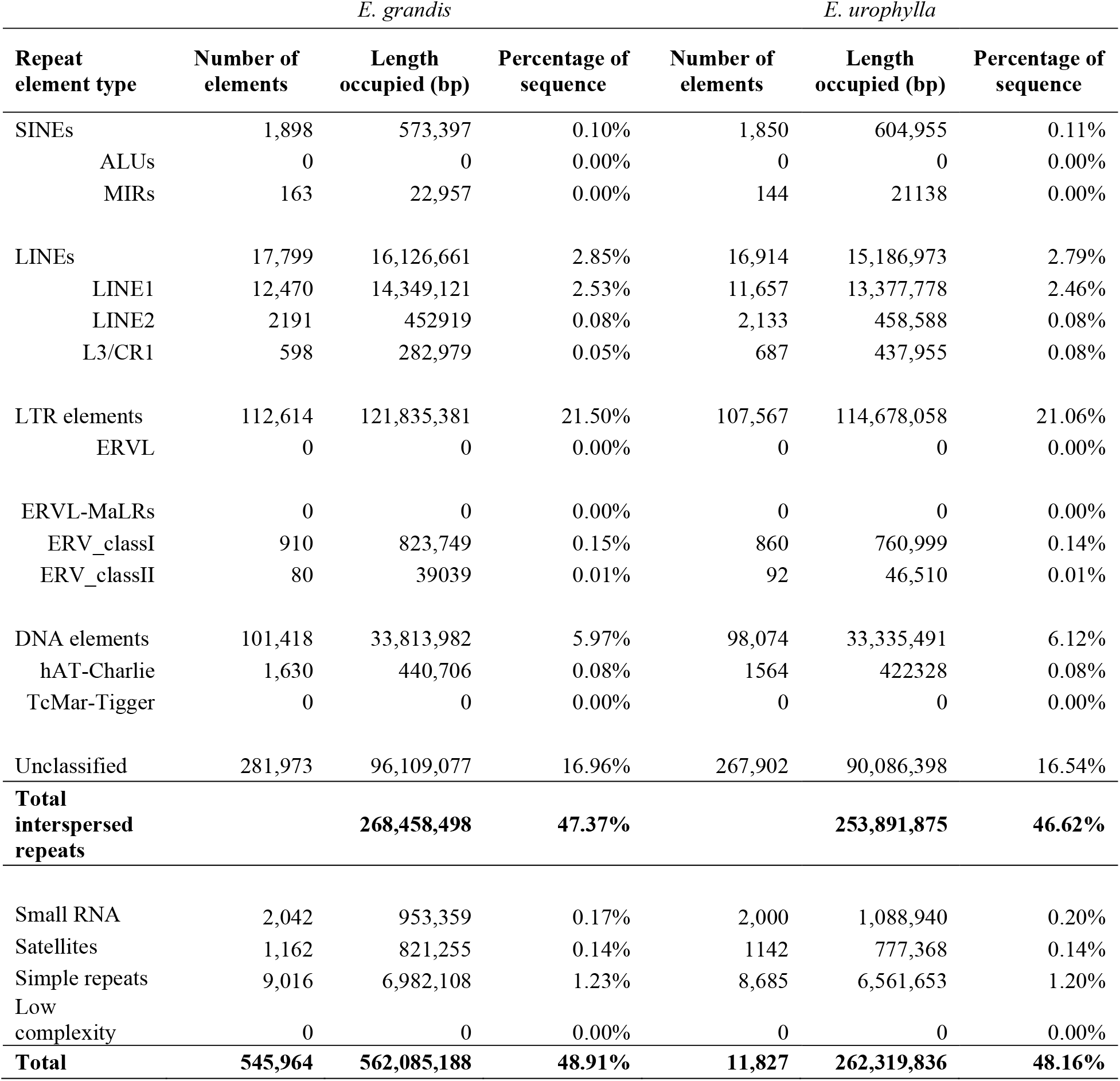
Repeat element content of assembled haplogenomes.

**Supplementary Table 6.**
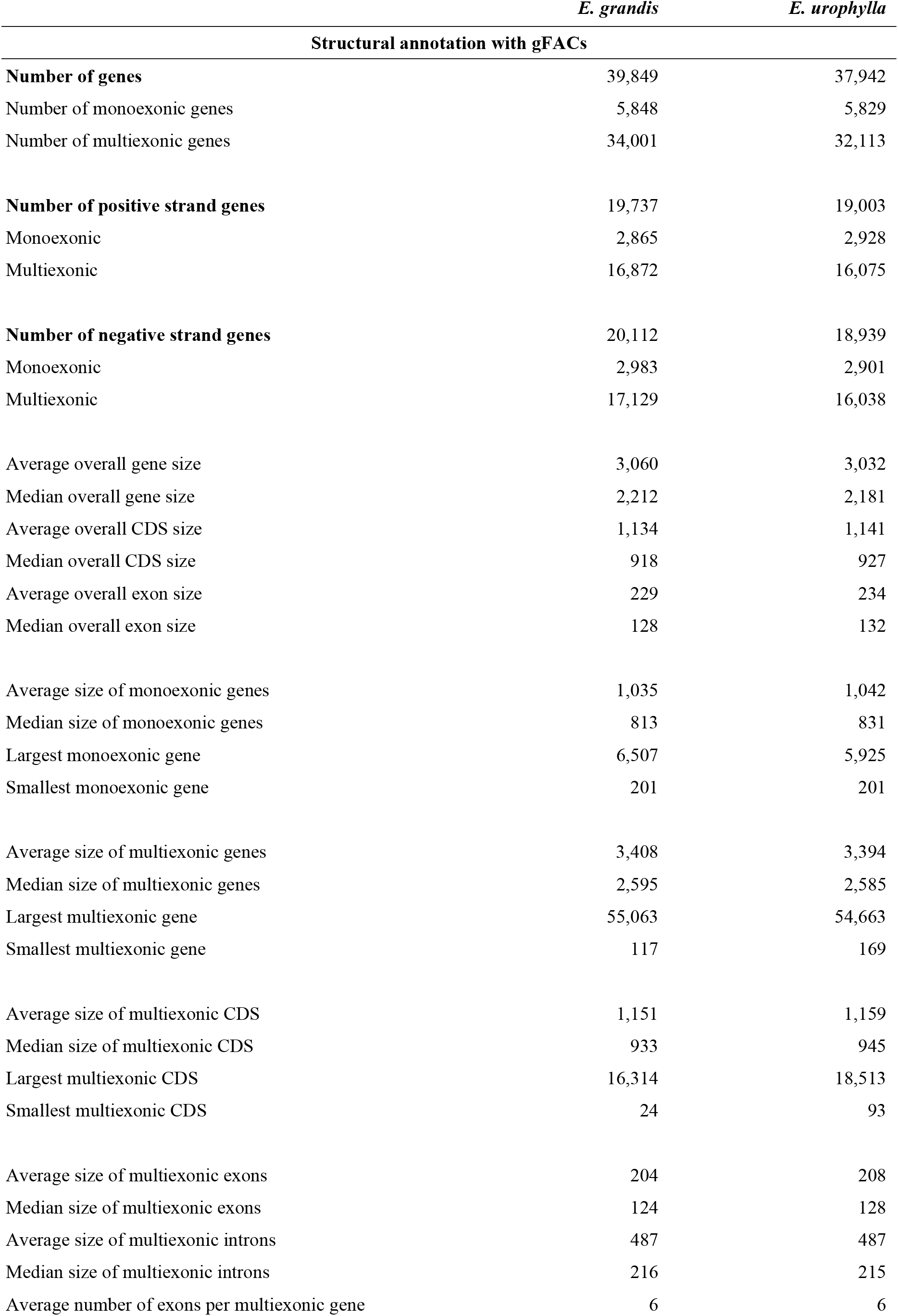
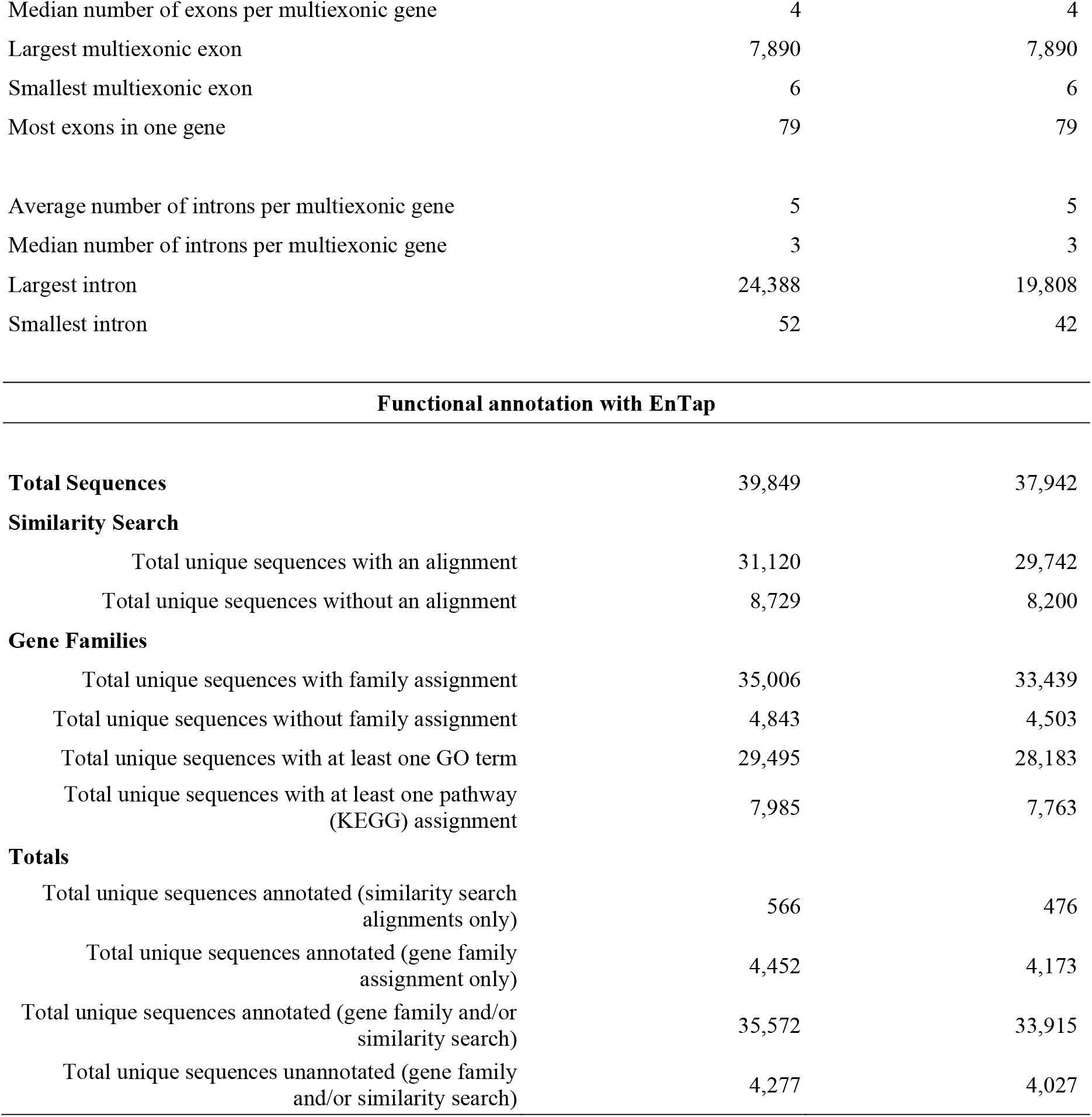
Haplogenome annotation statistics.

|  | *E. grandis* | *E. urophylla* |
| --- | --- | --- |
| **Structural annotation with gFACs** |  |  |
| **Number of genes** | 39,849 | 37,942 |
| Number of monoexonic genes | 5,848 | 5,829 |
| Number of multiexonic genes | 34,001 | 32,113 |
| **Number of positive strand genes** | 19,737 | 19,003 |
| Monoexonic | 2,865 | 2,928 |
| Multiexonic | 16,872 | 16,075 |
| **Number of negative strand genes** | 20,112 | 18,939 |
| Monoexonic | 2,983 | 2,901 |
| Multiexonic | 17,129 | 16,038 |
| Average overall gene size | 3,060 | 3,032 |
| Median overall gene size | 2,212 | 2,181 |
| Average overall CDS size | 1,134 | 1,141 |
| Median overall CDS size | 918 | 927 |
| Average overall exon size | 229 | 234 |
| Median overall exon size | 128 | 132 |
| Average size of monoexonic genes | 1,035 | 1,042 |
| Median size of monoexonic genes | 813 | 831 |
| Largest monoexonic gene | 6,507 | 5,925 |
| Smallest monoexonic gene | 201 | 201 |
| Average size of multiexonic genes | 3,408 | 3,394 |
| Median size of multiexonic genes | 2,595 | 2,585 |
| Largest multiexonic gene | 55,063 | 54,663 |
| Smallest multiexonic gene | 117 | 169 |
| Average size of multiexonic CDS | 1,151 | 1,159 |
| Median size of multiexonic CDS | 933 | 945 |
| Largest multiexonic CDS | 16,314 | 18,513 |
| Smallest multiexonic CDS | 24 | 93 |
| Average size of multiexonic exons | 204 | 208 |
| Median size of multiexonic exons | 124 | 128 |
| Average size of multiexonic introns | 487 | 487 |
| Median size of multiexonic introns | 216 | 215 |
| Average number of exons per multiexonic gene | 6 | 6 |
| Median number of exons per multiexonic gene | 4 | 4 |
| Largest multiexonic exon | 7,890 | 7,890 |
| Smallest multiexonic exon | 6 | 6 |
| Most exons in one gene | 79 | 79 |
| Average number of introns per multiexonic gene | 5 | 5 |
| Median number of introns per multiexonic gene | 3 | 3 |
| Largest intron | 24,388 | 19,808 |
| Smallest intron | 52 | 42 |
| **Functional annotation with EnTap** |  |  |
| **Total Sequences** | 39,849 | 37,942 |
| **Similarity Search** |  |  |
| Total unique sequences with an alignment | 31,120 | 29,742 |
| Total unique sequences without an alignment | 8,729 | 8,200 |
| **Gene Families** |  |  |
| Total unique sequences with family assignment | 35,006 | 33,439 |
| Total unique sequences without family assignment | 4,843 | 4,503 |
| Total unique sequences with at least one GO term | 29,495 | 28,183 |
| Total unique sequences with at least one pathway<br>(KEGG) assignment | 7,985 | 7,763 |
| **Totals** |  |  |
| Total unique sequences annotated (similarity search<br>alignments only) | 566 | 476 |
| Total unique sequences annotated (gene family<br>assignment only) | 4,452 | 4,173 |
| Total unique sequences annotated (gene family and/or<br>similarity search) | 35,572 | 33,915 |
| Total unique sequences unannotated (gene family<br>and/or similarity search) | 4,277 | 4,027 |

**Supplementary Table 7.** Number and total length of syntenic and rearranged regions in the *E. grandis* and *E. urophylla* haplogenomes. Regions are shown between the *E. grandis* v2.0 reference genome and the *E. grandis* haplogenome as well as the *E. grandis* haplogenome and the *E. urophylla* haplogenome. Rearrangements were called with SyRI (Synteny and rearrangement identifier) with a minimum 100 bp size, using the *E. grandis* v2.0, *E. grandis* or *E. urophylla* haplogenome as the reference in the analyses. Length is indicated in basepairs (bp). Only the eleven scaffolded chromosomes were compared for identification of rearranged regions.

| *E. grandis* v2.0 (reference) vs *E. grandis* haplogenome (query) |  |  |  |  |  |  |  |
| --- | --- | --- | --- | --- | --- | --- | --- |
| Variation type<sup>a</sup> | SYN | INV | TRANS | DUP (v2.0) | DUP (*E. grandis*) | Not aligned (v2.0) | Not aligned (*E. grandis*) |
| Count | 13,463 | 167 | 9,290 | 29,596 | 17,519 | 28,761 | 18,904 |
| Length (v2.0) | 317,981,657 | 57,482,207 | 75,974,491 | 141,439,740 | - | 111,692,400 | - |
| Length (*E. grandis*) | 317,513,455 | 45,151,373 | 75,544,141 | - | 50,831,969 | - | 41,963,576 |

| *E. grandis* haplogenome (reference) vs *E. urophylla* haplogenome (query) |  |  |  |  |  |  |  |
| --- | --- | --- | --- | --- | --- | --- | --- |
| Variation type | SYN | INV | TRANS | DUP (*E. grandis*) | DUP (*E. urophylla*) | Not aligned (*E. grandis*) | Not aligned (*E. urophylla*) |
| Count | 15,236 | 189 | 10,526 | 21,149 | 16,865 | 24,700 | 22,770 |
| Length (*E. grandis*) | 256,747,807 | 54,233,806 | 89,269,151 | 159,115,873 | - | 72,234,120 | - |
| Length (*E. urophylla*) | 256,876,296 | 55,605,620 | 88,801,518 | - | 55,495,066 | - | 62,322,133 |

| *E. urophylla* haplogenome (reference) vs *E. grandis* haplogenome (query) |  |  |  |  |  |  |  |
| --- | --- | --- | --- | --- | --- | --- | --- |
| Variation type | SYN | INV | TRANS | DUP (*E. urophylla*) | DUP (*E. grandis*) | Not aligned (*E. urophylla*) | Not aligned (*E. grandis*) |
| Count | 15,245 | 187 | 10,502 | 19,302 | 18,916 | 22,849 | 24,754 |
| Length (*E. urophylla*) | 256,751,376 | 55,557,398 | 86,591,743 | 145,450,405 | - | 62,437,935 | - |
| Length (*E. grandis*) | 256,583,791 | 54,570,060 | 86,799,621 | - | 63,010,166 | - | 72,125,631 |

SYN: syntenic region, INV: inversion, TRANS: translocation, DUP: duplication in the genome indicated in brackets, where v2.0 is the *E. grandis* v2.0 reference genome, *E. grandis* the *E. grandis* haplogenome and *E. urophylla* the *E. urophylla* haplogenome, Not aligned: unaligned regions in *E. grandis* or *E. urophylla* (query) haplogenome.

**Supplementary Table 8.** Number and total length of local sequence variation in syntenic and rearranged region in the *E. grandis* and *E. urophylla* haplogenomes. Local sequence variants were called with SyRI using the *E. grandis* v2.0, *E. grandis* and *E. urophylla* haplogenome as the reference genome respectively. Only the eleven scaffolded chromosomes were compared for local sequence variant identification.

| Variation type | SNPs | Insertions | Deletions | Copygains | Copylosses | Highly diverged | Tandem repeats | Total |
| --- | --- | --- | --- | --- | --- | --- | --- | --- |
| Count | 6,373,115 | 539,605 | 578,616 | 1,759 | 1,665 | 9,202 | 321 | 7,504,283 |
| Length *E. grandis* v2.0 | 6,373,115 | - | 5,615,561 | - | 10,885,284 | 45,970,827 | 685,226 | 69,530,013 |
| Length *E. grandis* | 6,373,115 | 6,116,677 | - | 11,424,778 | - | 31,670,584 | 870,720 | 56,455,874 |

***E. grandis* haplogenome (reference) vs *E. urophylla* haplogenome (query)**
| Variation type | SNPs | Insertions | Deletions | Copygains | Copylosses | Highly diverged | Tandem repeats | Total |
| --- | --- | --- | --- | --- | --- | --- | --- | --- |
| Count | 8,376,569 | 676,636 | 704,383 | 2,172 | 2,127 | 8,018 | 268 | 9,770,173 |
| Length *E. grandis* | 8,376,569 | - | 8,412,691 | - | 9,689,158 | 38,129,322 | 656,872 | 65,264,612 |
| Length *E. urophylla* | 8,376,569 | 7,629,721 | - | 9,578,692 | - | 40,181,747 | 680,693 | 66,447,422 |

***E. urophylla* haplogenome (reference) vs *E. grandis* haplogenome (query)**
| Variation type | SNPs | Insertions | Deletions | Copygains | Copylosses | Highly diverged | Tandem repeats | Total |
| --- | --- | --- | --- | --- | --- | --- | --- | --- |
| Count | 8,350,925 | 694,782 | 681,344 | 2,128 | 2,135 | 7,931 | 269 | 9,739,514 |
| Length *E. urophylla* | 8,350,925 | - | 7,603,918 | - | 9,326,217 | 39,641,637 | 680,104 | 65,602,801 |
| Length *E. grandis* | 8,350,925 | 8,043,419 | - | 9,884,667 | - | 37,649,655 | 653,045 | 64,581,711 |

**Supplementary Table 9.** Inversions larger than 50 kb between the *E. grandis* and *E. urophylla* haplogenomes.

| *E. grandis* haplogenome |  |  |  | *E. urophylla* haplogenome |  |  |  |
| --- | --- | --- | --- | --- | --- | --- | --- |
| Chr | Start | End | Reference length | Chr | Start | End | Query length |
| Chr01 | 23,388,498 | 23,774,716 | 386,218 | Chr01 | 27,866,247 | 28,333,853 | 467,606 |
| Chr01 | 29,088,860 | 29,535,584 | 446,724 | Chr01 | 33,200,026 | 33,802,257 | 602,231 |
| Chr01 | 38,325,932 | 38,988,219 | 662,287 | Chr01 | 43,142,878 | 43,537,862 | 394,984 |
| Chr01 | 38,989,424 | 39,795,647 | 806,223 | Chr01 | 43,575,027 | 44,294,133 | 719,106 |
| Chr02 | 10,256,034 | 11,495,357 | 1,239,323 | Chr02 | 12,449,219 | 13,806,880 | 1,357,661 |
| Chr02 | 16,268,629 | 18,735,729 | 2,467,100 | Chr02 | 16,317,615 | 19,174,190 | 2,856,575 |
| Chr02 | 21,273,193 | 21,613,433 | 340,240 | Chr02 | 21,133,426 | 21,629,094 | 495,668 |
| Chr02 | 21,614,540 | 21,866,290 | 251,750 | Chr02 | 22,058,000 | 22,319,351 | 261,351 |
| Chr02 | 24,530,674 | 24,753,625 | 222,951 | Chr02 | 23,723,944 | 23,839,960 | 116,016 |
| Chr02 | 34,946,307 | 35,416,437 | 470,130 | Chr02 | 33,832,261 | 34,243,757 | 411,496 |
| Chr02 | 7,710,307 | 8,128,604 | 418,297 | Chr02 | 8,216,904 | 8,593,150 | 376,246 |
| Chr02 | 9,120,701 | 9,141,333 | 20,632 | Chr02 | 9,309,778 | 9,532,539 | 222,761 |
| Chr02 | 9,543,813 | 10,251,800 | 707,987 | Chr02 | 10,024,991 | 11,058,528 | 1,033,537 |
| Chr03 | 17,924,896 | 18,361,694 | 436,798 | Chr03 | 12,844,626 | 13,544,499 | 699,873 |
| Chr03 | 22,750,923 | 23,075,176 | 324,253 | Chr03 | 17,872,261 | 18,383,259 | 510,998 |
| Chr03 | 24,894,740 | 25,020,737 | 125,997 | Chr03 | 19,732,494 | 19,842,712 | 110,218 |
| Chr03 | 26,582,840 | 26,695,689 | 112,849 | Chr03 | 22,211,437 | 22,302,731 | 91,294 |
| Chr03 | 29,450,360 | 29,499,020 | 48,660 | Chr03 | 25,057,971 | 25,153,753 | 95,782 |
| Chr03 | 31,007,465 | 33,222,991 | 2,215,526 | Chr03 | 27,113,759 | 29,387,690 | 2,273,931 |
| Chr03 | 42,077,277 | 42,865,022 | 787,745 | Chr03 | 39,421,192 | 40,598,627 | 1,177,435 |
| Chr03 | 52,056,591 | 52,768,850 | 712,259 | Chr03 | 46,996,755 | 47,400,472 | 403,717 |
| Chr04 | 10,204,799 | 10,608,432 | 403,633 | Chr04 | 9,959,942 | 10,372,400 | 412,458 |
| Chr04 | 14,353,739 | 15,314,090 | 960,351 | Chr04 | 13,693,800 | 13,989,776 | 295,976 |
| Chr04 | 18,680,518 | 19,330,061 | 649,543 | Chr04 | 18,130,979 | 18,386,712 | 255,733 |
| Chr04 | 19,856,681 | 19,916,877 | 60,196 | Chr04 | 18,916,777 | 18,948,259 | 31,482 |
| Chr04 | 24,074,346 | 25,225,372 | 1,151,026 | Chr04 | 23,089,700 | 24,877,693 | 1,787,993 |
| Chr04 | 285,506 | 1,781,371 | 1,495,865 | Chr04 | 8 | 1,997,000 | 1,996,992 |
| Chr04 | 5,839,102 | 7,451,977 | 1,612,875 | Chr04 | 6,099,540 | 7,272,200 | 1,172,660 |
| Chr05 | 1,139 | 191,613 | 190,474 | Chr05 | 1 | 199,651 | 199,650 |
| Chr05 | 13,767,410 | 14,264,679 | 497,269 | Chr05 | 12,544,177 | 13,230,121 | 685,944 |
| Chr05 | 18,731,556 | 18,772,986 | 41,430 | Chr05 | 17,841,218 | 18,037,838 | 196,620 |
| Chr05 | 23,214,319 | 23,331,219 | 116,900 | Chr05 | 20,525,113 | 20,682,115 | 157,002 |
| Chr05 | 26,312,453 | 26,639,176 | 326,723 | Chr05 | 23,326,012 | 23,619,675 | 293,663 |
| Chr05 | 39,764,872 | 40,079,538 | 314,666 | Chr05 | 31,002,806 | 31,460,527 | 457,721 |
| Chr05 | 3,977,112 | 4,219,061 | 241,949 | Chr05 | 3,704,167 | 3,953,788 | 249,621 |
| Chr05 | 40,147,672 | 40,620,311 | 472,639 | Chr05 | 31,467,622 | 32,031,945 | 564,323 |
| Chr05 | 40,664,102 | 41,683,031 | 1,018,929 | Chr05 | 32,044,329 | 32,499,398 | 455,069 |
| Chr05 | 41,852,810 | 42,212,844 | 360,034 | Chr05 | 32,508,317 | 32,842,567 | 334,250 |
| Chr05 | 42,231,666 | 42,497,444 | 265,778 | Chr05 | 33,225,046 | 33,730,213 | 505,167 |
| Chr05 | 45,116,621 | 45,471,128 | 354,507 | Chr05 | 36,775,424 | 37,162,018 | 386,594 |
| Chr05 | 4,790,253 | 5,284,213 | 493,960 | Chr05 | 4,587,258 | 5,054,184 | 466,926 |
| Chr05 | 6,762,823 | 7,811,079 | 1,048,256 | Chr05 | 6,186,655 | 7,075,904 | 889,249 |
| Chr06 | 1,422,135 | 2,104,051 | 681,916 | Chr06 | 1,501,336 | 2,307,495 | 806,159 |
| Chr06 | 17,562 | 116,068 | 98,506 | Chr06 | 1 | 105,034 | 105,033 |
| Chr06 | 18,585,323 | 19,047,661 | 462,338 | Chr06 | 17,287,053 | 17,586,309 | 299,256 |
| Chr06 | 21,901,981 | 22,015,039 | 113,058 | Chr06 | 17,587,701 | 17,694,441 | 106,740 |
| Chr06 | 37,843,975 | 37,953,970 | 109,995 | Chr06 | 36,451,444 | 36,543,431 | 91,987 |
| Chr06 | 4,348,505 | 5,014,910 | 666,405 | Chr06 | 4,809,501 | 5,634,648 | 825,147 |
| Chr06 | 46,380,498 | 46,880,027 | 499,529 | Chr06 | 42,887,607 | 43,469,053 | 581,446 |
| Chr06 | 49,293,183 | 49,600,931 | 307,748 | Chr06 | 46,607,955 | 46,889,702 | 281,747 |
| Chr06 | 5,790,165 | 6,245,897 | 455,732 | Chr06 | 6,531,092 | 6,952,534 | 421,442 |
| Chr06 | 6,276,170 | 6,681,490 | 405,320 | Chr06 | 6,952,991 | 7,550,358 | 597,367 |
| Chr07 | 10,195,846 | 11,089,184 | 893,338 | Chr07 | 8,134,852 | 8,493,558 | 358,706 |
| Chr07 | 11,303,891 | 12,661,932 | 1,358,041 | Chr07 | 8,550,716 | 13,463,227 | 4,912,511 |
| Chr07 | 13,073 | 685,800 | 672,727 | Chr07 | 53,539 | 498,307 | 444,768 |
| Chr07 | 15,398,973 | 15,469,228 | 70,255 | Chr07 | 14,007,680 | 14,077,253 | 69,573 |
| Chr07 | 15,548,182 | 15,826,775 | 278,593 | Chr07 | 14,078,345 | 14,337,355 | 259,010 |
| Chr07 | 17,183,241 | 17,610,391 | 427,150 | Chr07 | 15,818,970 | 16,167,843 | 348,873 |
| Chr07 | 18,528,803 | 18,920,067 | 391,264 | Chr07 | 16,770,493 | 17,235,509 | 465,016 |
| Chr07 | 20,211,368 | 22,642,279 | 2,430,911 | Chr07 | 19,458,012 | 21,757,921 | 2,299,909 |
| Chr07 | 23,468,568 | 23,626,990 | 158,422 | Chr07 | 21,768,463 | 21,937,670 | 169,207 |
| Chr07 | 24,191,002 | 24,827,475 | 636,473 | Chr07 | 23,584,113 | 24,382,712 | 798,599 |
| Chr07 | 31,325,104 | 31,831,058 | 505,954 | Chr07 | 31,095,389 | 31,656,188 | 560,799 |
| Chr07 | 3,157,850 | 4,559,613 | 1,401,763 | Chr07 | 2,510,914 | 2,920,972 | 410,058 |
| Chr08 | 12,725,330 | 12,863,580 | 138,250 | Chr08 | 11,688,255 | 11,819,476 | 131,221 |
| Chr08 | 16,700,735 | 17,318,973 | 618,238 | Chr08 | 14,631,661 | 15,206,380 | 574,719 |
| Chr08 | 19,875,397 | 21,709,986 | 1,834,589 | Chr08 | 18,154,032 | 21,017,051 | 2,863,019 |
| Chr08 | 2,510,912 | 3,087,516 | 576,604 | Chr08 | 2,492,921 | 2,934,491 | 441,570 |
| Chr08 | 27,297,239 | 28,670,928 | 1,373,689 | Chr08 | 26,916,592 | 28,009,638 | 1,093,046 |
| Chr08 | 29,225,629 | 29,316,542 | 90,913 | Chr08 | 28,143,627 | 28,201,928 | 58,301 |
| Chr08 | 3,501,184 | 5,038,276 | 1,537,092 | Chr08 | 3,372,176 | 4,298,000 | 925,824 |
| Chr08 | 419,214 | 1,055,841 | 636,627 | Chr08 | 407,448 | 877,416 | 469,968 |
| Chr08 | 45,309,936 | 45,408,786 | 98,850 | Chr08 | 42,198,433 | 42,388,897 | 190,464 |
| Chr08 | 45,522,689 | 46,091,114 | 568,425 | Chr08 | 42,436,168 | 42,559,996 | 123,828 |
| Chr08 | 63,156,082 | 63,769,512 | 613,430 | Chr08 | 59,525,804 | 60,126,627 | 600,823 |
| Chr09 | 13,376,823 | 13,702,567 | 325,744 | Chr09 | 12,998,893 | 13,338,127 | 339,234 |
| Chr09 | 2,707,324 | 3,062,759 | 355,435 | Chr09 | 3,777,976 | 4,028,769 | 250,793 |
| Chr09 | 27,619,973 | 27,697,783 | 77,810 | Chr09 | 26,834,362 | 26,876,310 | 41,948 |
| Chr09 | 3,101,186 | 3,530,067 | 428,881 | Chr09 | 4,029,319 | 4,440,544 | 411,225 |
| Chr09 | 6,826,654 | 8,005,141 | 1,178,487 | Chr09 | 7,719,031 | 8,059,633 | 340,602 |
| Chr10 | 233,352 | 527,104 | 293,752 | Chr10 | 20,102 | 323,104 | 303,002 |
| Chr10 | 24,242,636 | 24,889,751 | 647,115 | Chr10 | 27,630,162 | 28,354,048 | 723,886 |
| Chr10 | 26,475,377 | 26,532,239 | 56,862 | Chr10 | 29,859,221 | 29,888,792 | 29,571 |
| Chr10 | 26,902,198 | 26,970,937 | 68,739 | Chr10 | 30,235,379 | 30,306,797 | 71,418 |
| Chr10 | 34,082,797 | 34,441,174 | 358,377 | Chr10 | 38,683,430 | 39,016,229 | 332,799 |
| Chr10 | 35,068,172 | 35,243,214 | 175,042 | Chr10 | 39,806,630 | 39,909,104 | 102,474 |
| Chr11 | 11,262,172 | 11,396,228 | 134,056 | Chr11 | 11,417,392 | 11,457,497 | 40,105 |
| Chr11 | 13,155,091 | 13,298,496 | 143,405 | Chr11 | 12,829,340 | 13,017,471 | 188,131 |
| Chr11 | 1,408,650 | 1,977,286 | 568,636 | Chr11 | 1,848,260 | 2,259,527 | 411,267 |
| Chr11 | 20,764,115 | 20,973,486 | 209,371 | Chr11 | 19,775,853 | 20,065,496 | 289,643 |
| Chr11 | 25,701,549 | 26,104,207 | 402,658 | Chr11 | 21,898,345 | 22,199,580 | 301,235 |
| Chr11 | 31,583,111 | 32,300,126 | 717,015 | Chr11 | 27,699,796 | 28,409,715 | 709,919 |
| Chr11 | 35,437,224 | 35,496,143 | 58,919 | Chr11 | 31,711,764 | 31,775,563 | 63,799 |
| Chr11 | 5,589,851 | 6,679,990 | 1,090,139 | Chr11 | 5,522,667 | 6,499,452 | 976,785 |

**Supplementary Table 10.** Translocations between the *E. grandis* and *E. urophylla* haplogenomes that are larger than 50 kb.

| *E. grandis* haplogenome |  |  |  | *E. urophylla* haplogenome |  |  |  |
| --- | --- | --- | --- | --- | --- | --- | --- |
| Chr | Start | End | Reference length | Chr | Start | End | Query length |
| Chr01 | 2,749,813 | 2,819,595 | 69,782 | Chr01 | 1,917,560 | 1,987,365 | 69,805 |
| Chr02 | 8,220,591 | 8,275,256 | 54,665 | Chr02 | 23,641,937 | 23,696,439 | 54,502 |
| Chr04 | 12,396,028 | 12,452,029 | 56,001 | Chr04 | 12,703,390 | 12,759,400 | 56,010 |
| Chr06 | 20,361,544 | 20,482,641 | 121,097 | Chr06 | 21,352,697 | 21,473,569 | 120,872 |
| Chr06 | 20,682,722 | 20,765,954 | 83,232 | Chr06 | 21,659,542 | 21,742,402 | 82,860 |
| Chr07 | 43,614,628 | 43,679,697 | 65,069 | Chr10 | 7,513,443 | 7,578,449 | 65,006 |
| Chr07 | 43,753,217 | 43,808,068 | 54,851 | Chr10 | 7,644,800 | 7,699,782 | 54,982 |
| Chr07 | 43,934,853 | 44,028,924 | 94,071 | Chr10 | 7,818,126 | 7,912,179 | 94,053 |
| Chr07 | 44,029,147 | 44,090,096 | 60,949 | Chr10 | 7,912,179 | 7,972,966 | 60,787 |
| Chr08 | 28,793,518 | 28,854,941 | 61,423 | Chr08 | 26,691,561 | 26,752,939 | 61,378 |
| Chr08 | 28,960,832 | 29,054,796 | 93,964 | Chr08 | 26,841,892 | 26,935,863 | 93,971 |
| Chr09 | 8,771,435 | 8,829,203 | 57,768 | Chr06 | 45,043,500 | 45,101,519 | 58,019 |
| Chr09 | 8,837,050 | 8,896,455 | 59,405 | Chr06 | 45,101,515 | 45,160,936 | 59,421 |
| Chr10 | 7,148,524 | 7,214,985 | 66,461 | Chr10 | 6,545,748 | 6,612,181 | 66,433 |
| Chr11 | 23,115,517 | 23,174,363 | 58,846 | Chr02 | 37,900,157 | 37,958,643 | 58,486 |
| Chr11 | 24,151,203 | 24,203,956 | 52,753 | Chr02 | 39,003,539 | 39,056,219 | 52,680 |
| Chr11 | 24,238,142 | 24,290,314 | 52,172 | Chr02 | 39,079,522 | 39,131,773 | 52,251 |
| Chr11 | 24,387,150 | 24,509,566 | 122,416 | Chr02 | 39,238,929 | 39,361,120 | 122,191 |
| Chr11 | 24,591,384 | 24,648,523 | 57,139 | Chr02 | 39,436,979 | 39,493,981 | 57,002 |
| Chr11 | 24,855,007 | 24,924,836 | 69,829 | Chr02 | 39,719,434 | 39,789,354 | 69,920 |
| Chr11 | 24,936,151 | 25,001,436 | 65,285 | Chr02 | 39,800,932 | 39,866,164 | 65,232 |
| Chr11 | 25,001,695 | 25,072,755 | 71,060 | Chr02 | 39,866,476 | 39,937,490 | 71,014 |
| Chr11 | 25,123,131 | 25,186,141 | 63,010 | Chr02 | 39,980,961 | 40,044,235 | 63,274 |
| Chr11 | 25,240,856 | 25,293,593 | 52,737 | Chr02 | 40,102,319 | 40,155,043 | 52,724 |

## Supplementary Figures

**Supplementary Figure 1.**
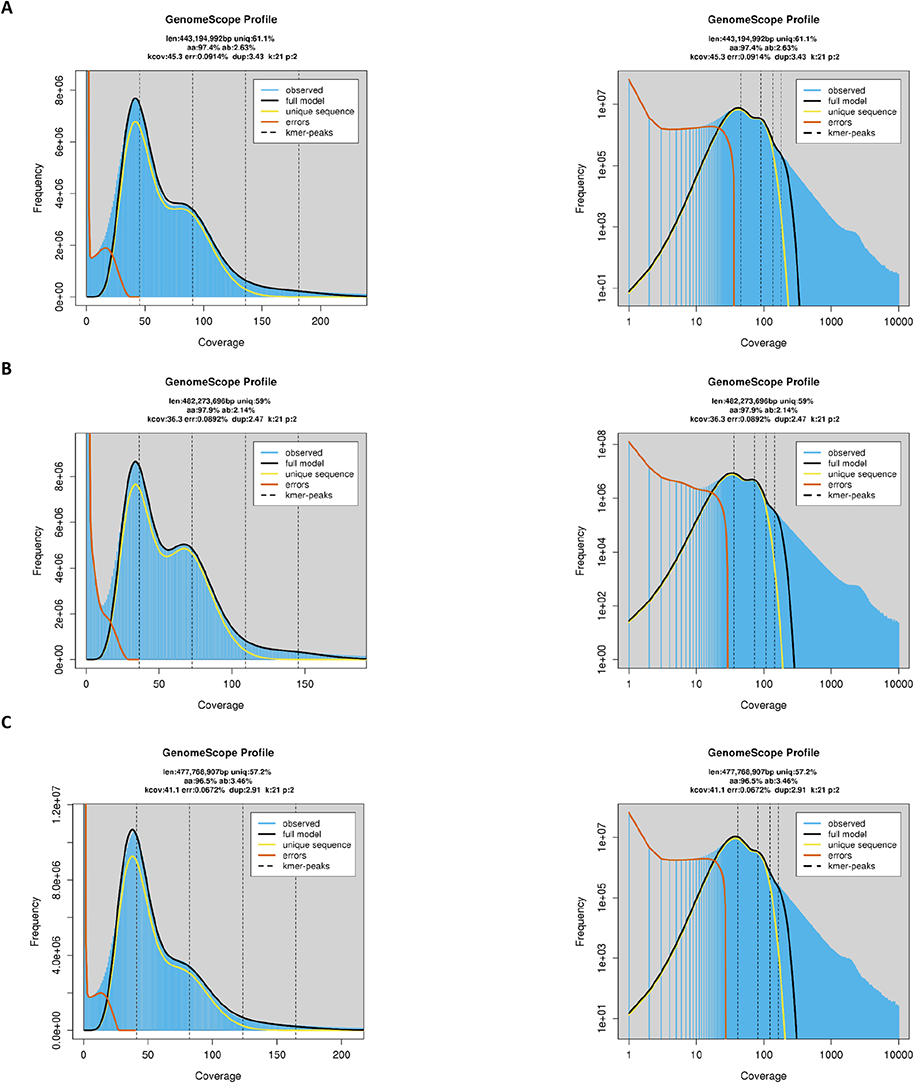
Genome size estimates for the (A) *E. urophylla*, (B) *E. grandis* and (C) the *E. urophylla* x *E. grandis* F_1_ hybrid genomes. Genome size was estimated at *k* = 21 with GenomeScope2.0.

**Supplementary Figure 2.**
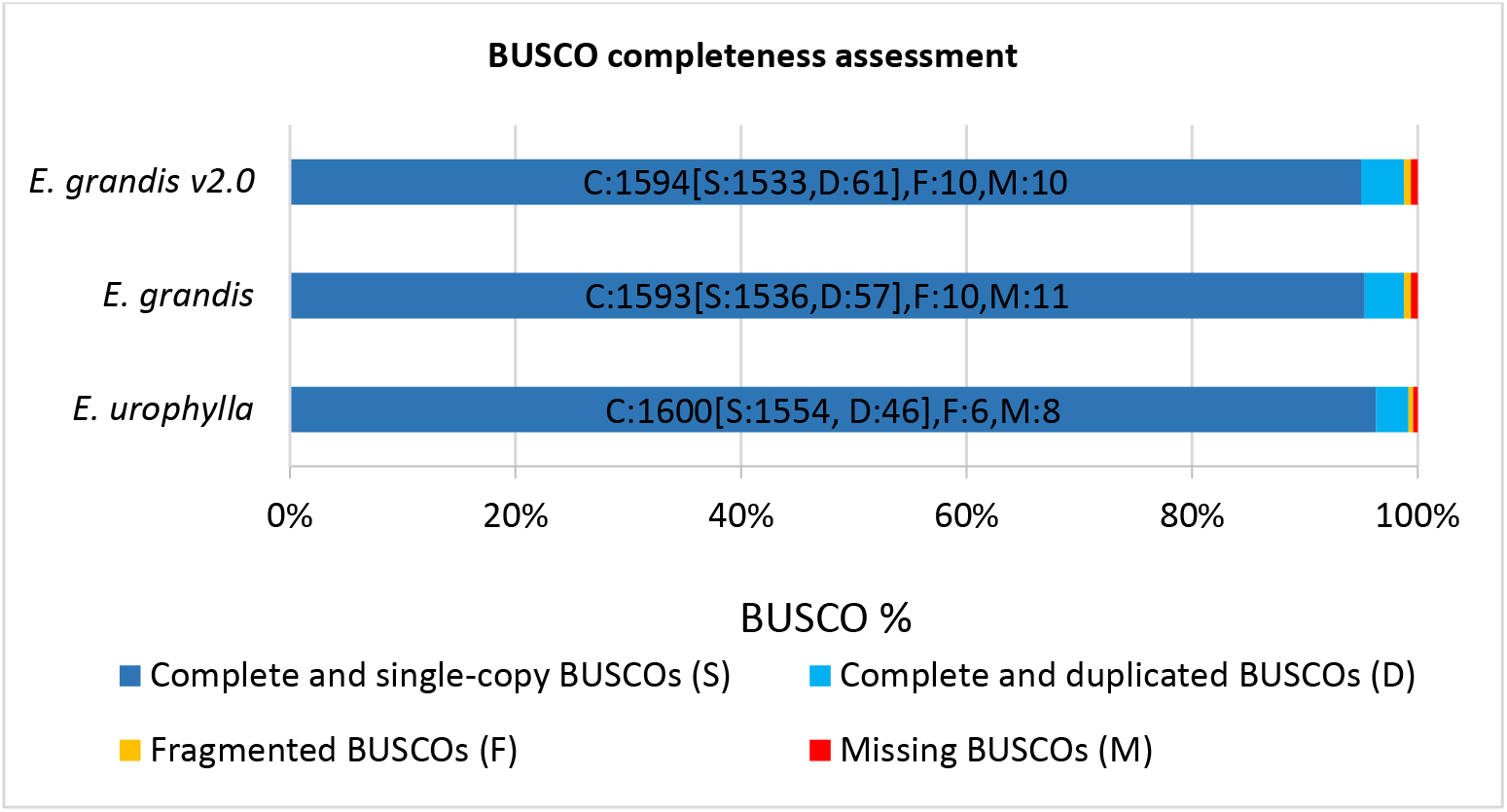
Benchmarking Universal Single-Copy Orthologs (BUSCO) completeness scores for both haplogenome assemblies as well as the currently available *E. grandis* v2.0 reference genome. A set of 1,614 embryophyte gene groups were used to calculate completeness of assembled genome. The bar indicates the percentage of genes belonging to categories as indicated by colour. The number of gene groups that are present (S - complete and single-copy, D - complete and duplicate-copy or F - fragmented) or absent (M - missing) are indicated by the numbers within the bar.

**Supplementary Figure 3.**
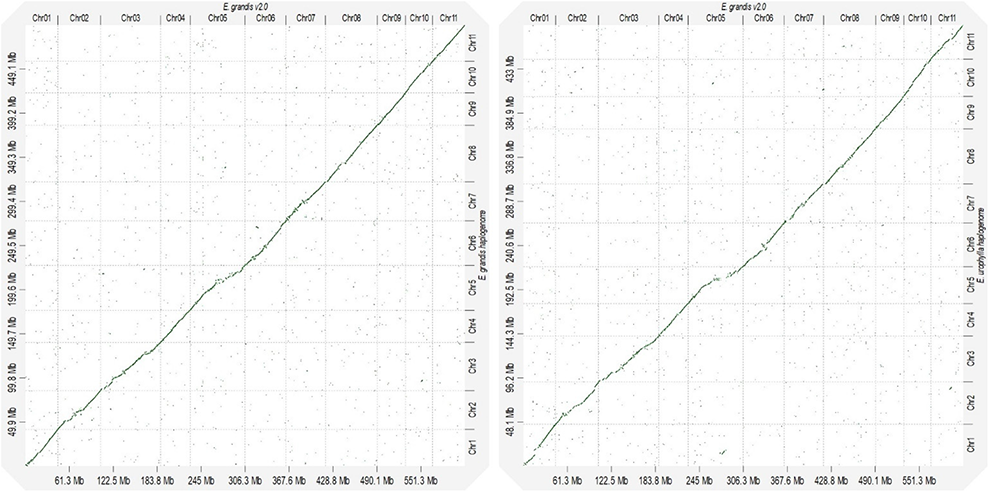
Alignment of placed haplogenome scaffolds to the *E. grandis* v2.0 reference genome. Alignments are shown for the *E. grandis* scaffolded haplogenome (y-axis) against the *E. grandis* v2.0 reference genome (x-axis) on the left and the *E. urophylla* scaffolded assembly (y-axis) against the *E. grandis* v2.0 reference genome (x-axis) on the right and is arranged by chromosome (from one to eleven). Alignment size is measured in megabases (Mb).

**Supplementary Figure 4.**
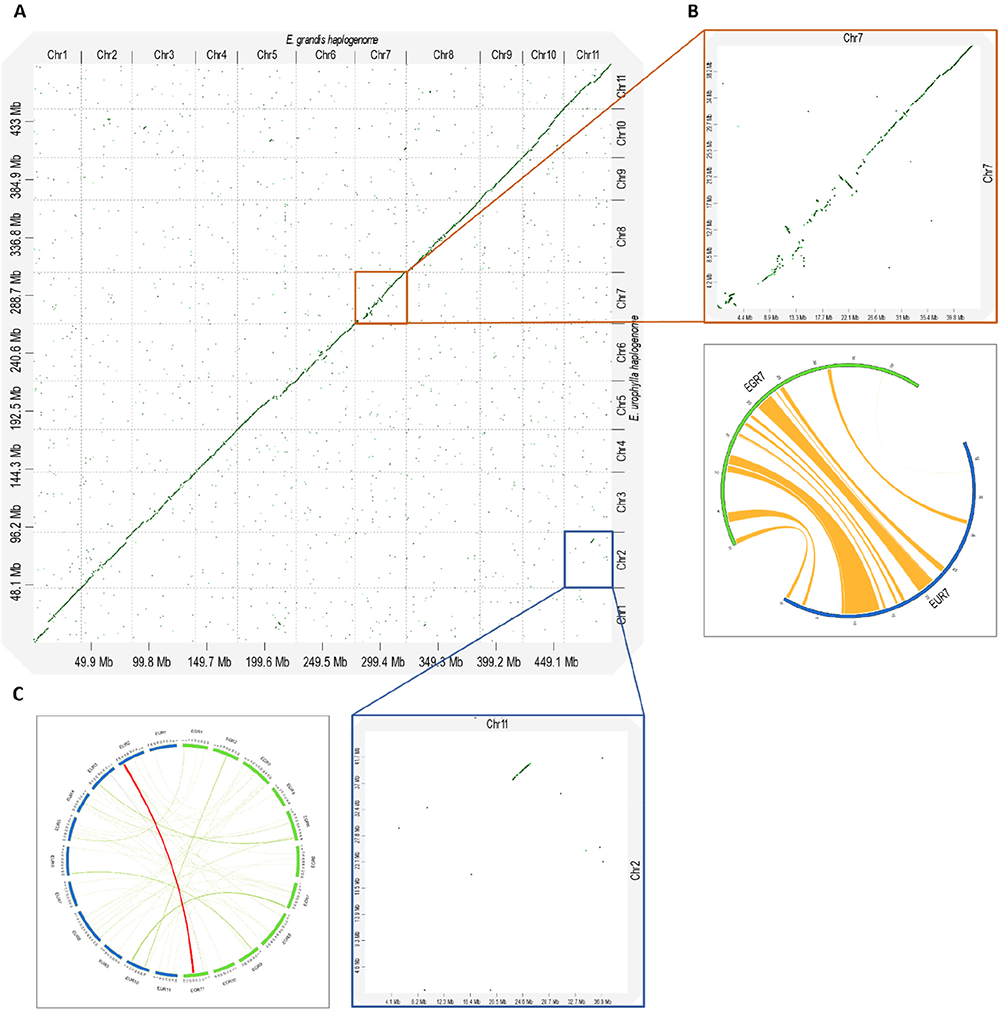
Alignment between the *E. grandis* and *E. urophylla* scaffolded haplogenome assemblies. (A) The *E. grandis* scaffolded haplogenome assembly (498.98 Mb) is shown on the x-axis and the *E. urophylla* scaffolded haplogenome assembly (481.16 Mb) on y-axis and is arranged by chromosome (from one to eleven). (B) The right-hand panel (orange block) is a zoom-in of an inversion on Chromosome 7 as seen with D-Genies (top), and a corresponding Circos visualization of the inversions called by SyRI (bottom). (C) The bottom panel (blue block) is a D-Genies zoom-in of a translocation from Chromosome 11 in *E. grandis* to Chromosome 2 in *E. urophylla* (on the right), and the corresponding event in a circos plot (highlighted in red). Alignment size is measured in megabases (Mb).

**Supplementary Figure 5.**
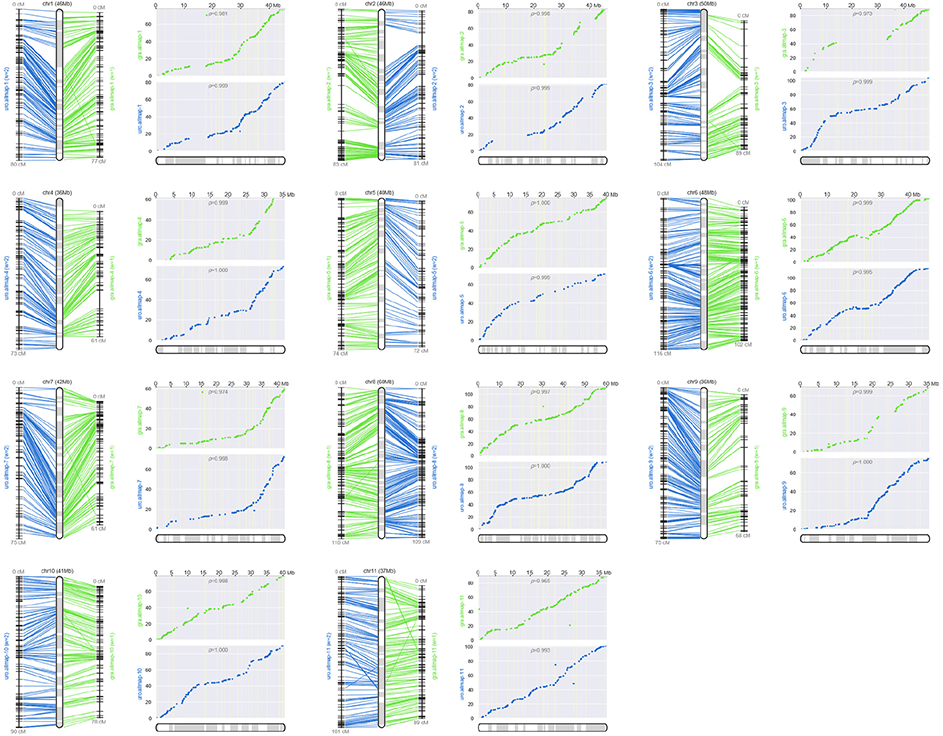
Pseudochromosomes of *E. urophylla* haplogenome, reconstructed from two genetic linkage input maps – uro.allmap and gra.allmap, with unequal weights (2 and 1 respectively). The left-hand panels for each chromosome represent CMAP-style presentation with lines connecting physical positions on the reconstructed chromosomes and genetic map positions of SNP markers used. Boxes alternating between grey and white in the CMAP-representations represent alternating scaffolds within the reconstructed chromosomes and mark scaffold boundaries. The right-hand panel has a set of two scatter plot, where dots on the x-axis represent the physical position on the chromosomes and the y-axis the map location for the *E. urophylla* (blue) and *E. grandis* (green). Pearson’s correlation coefficient is indicated as the *ρ*-value, and values range from −1 to 1 (where values closer to −1 and 1 indicates near-perfect collinearity).

**Supplementary Figure 6.**
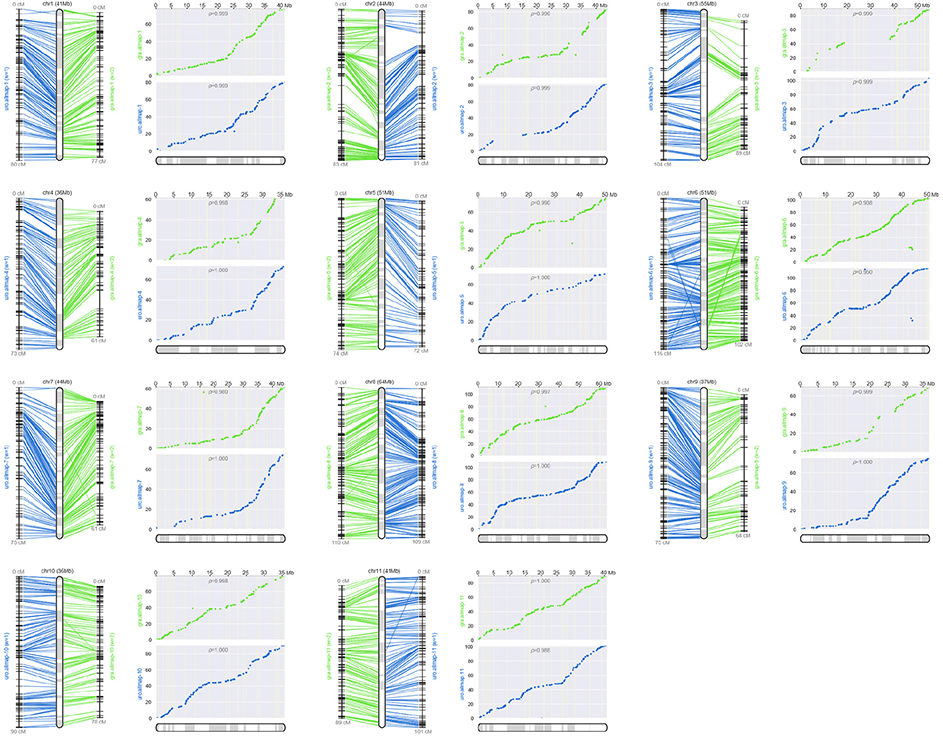
Pseudochromosomes of *E. grandis* haplogenome, reconstructed from two genetic linkage input maps – gra.allmap and uro.allmap, with unequal weights (2 and 1 respectively). The left-hand panels for each chromosome represent CMAP-style presentation with lines connecting physical positions on the reconstructed chromosomes and genetic map positions of SNP markers used. Boxes of alternating shades represent alternating scaffolds within the reconstructed chromosomes and mark scaffold boundaries. The right-hand panel has a set of two scatter plots, where dots on the x-axis represent the physical position on the chromosomes and the y-axis the map location for *E. urophylla* (blue) and *E. grandis* (green). Pearson’s correlation coefficient is also indicated (*ρ*-value), and values range from −1 to 1 (where values closer to −1 and 1 indicates near-perfect collinearity).

**Supplementary Figure 7.**
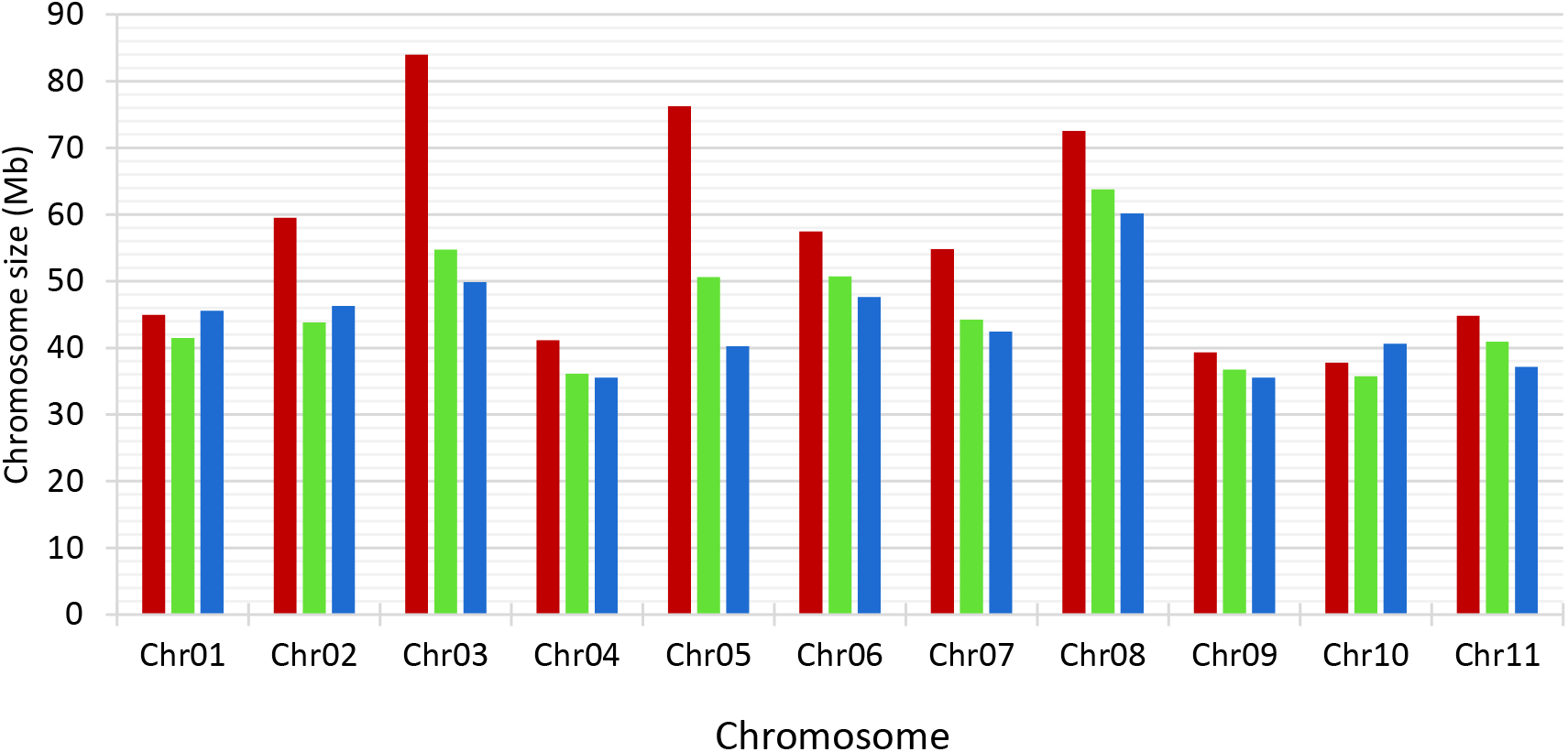
Scaffolded chromosome sizes of the *E. grandis* v2.0 and the scaffolded *E. grandis* and *E. urophylla* haplogenome assemblies. The total size of the 11 chromosomes is shown in megabases (Mb) for *E. grandis* v2.0 assembly (red), *E. grandis* (green) and *E. urophylla* (blue) haplogenome assemblies.

**Supplementary Figure 8.**
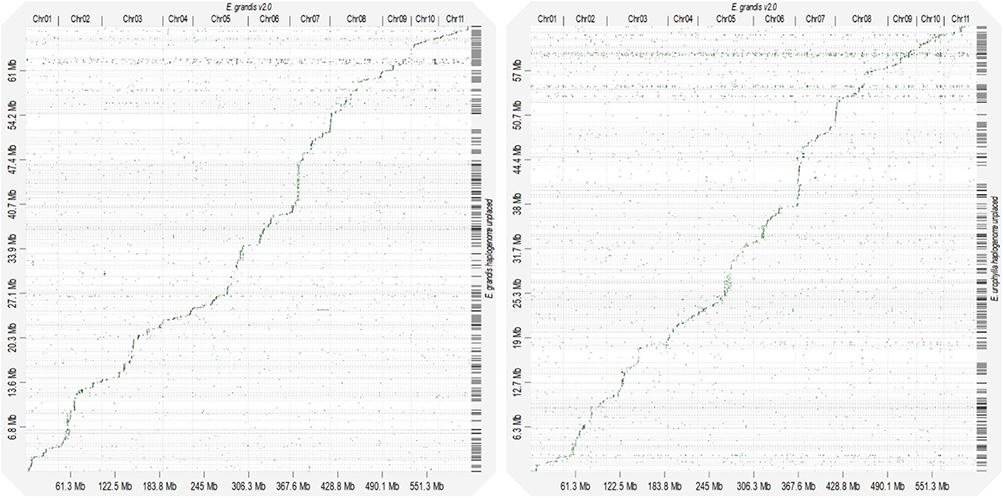
Alignment of unplaced *E. grandis* and *E. urophylla* haplogenome scaffolds to the *E. grandis* v2.0 reference genome. Alignments are shown for unplaced *E. grandis* scaffolds (y-axis) against the *E. grandis* v2.0 reference genome (x-axis) on the left and unplaced *E. urophylla* scaffolds (y-axis) against the *E. grandis* v2.0 reference genome (x-axis) on the right. Alignments are arranged by chromosome (from one to eleven) and alignment size is measured in megabases (Mb).

**Supplementary Figure 9.**
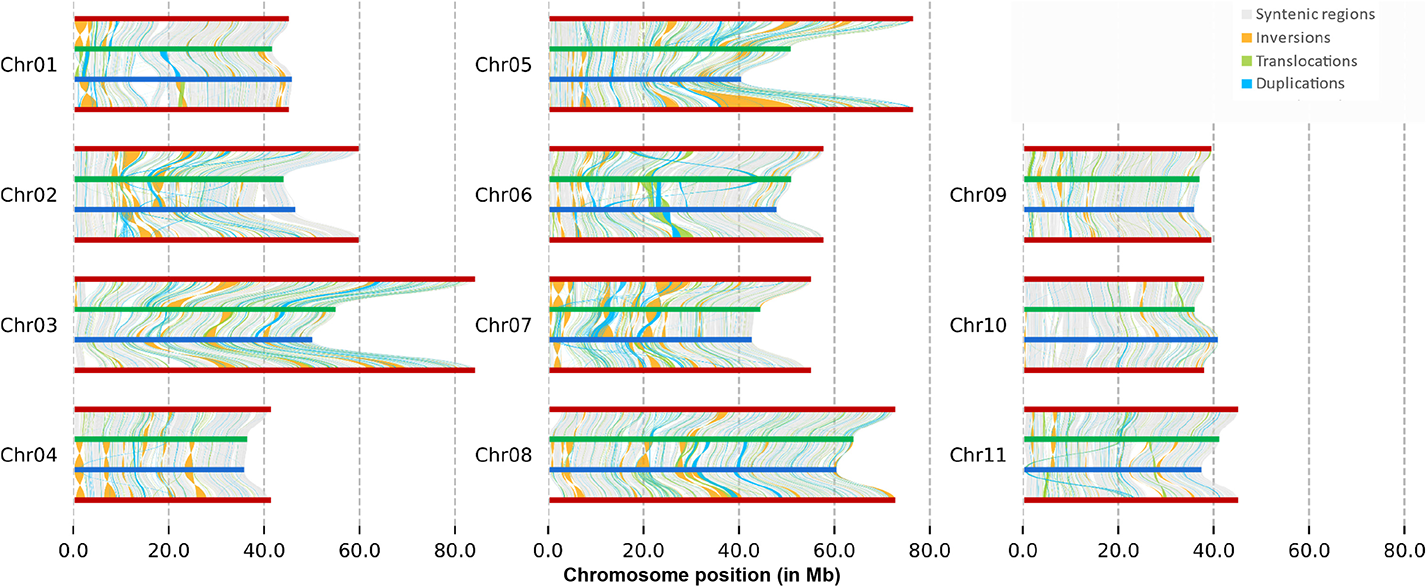
Syntenic and rearranged regions between the *E. grandis* v2.0, *E. grandis* and *E. urophylla* haplogenomes for all eleven chromosomes. Regions were identified using the Synteny and Rearrangement Identifier (SyRI), with a minimum rearrangement size of 100 base pairs. Chromosome number is indicated on the y-axis, while chromosome position is shown on the x-axis in megabase (Mb). Rearrangements are shown between the *E. grandis* v2.0 reference genome (dark red), *E. grandis* haplogenome (green) and the *E. urophylla* haplogenome (blue). Syntenic regions are indicated by grey lines, whereas non-aligned regions appear as white gaps in between syntenic regions. Green lines indicate translocations, orange indicate inversions and light blue translocations

**Supplementary Figure 10.**
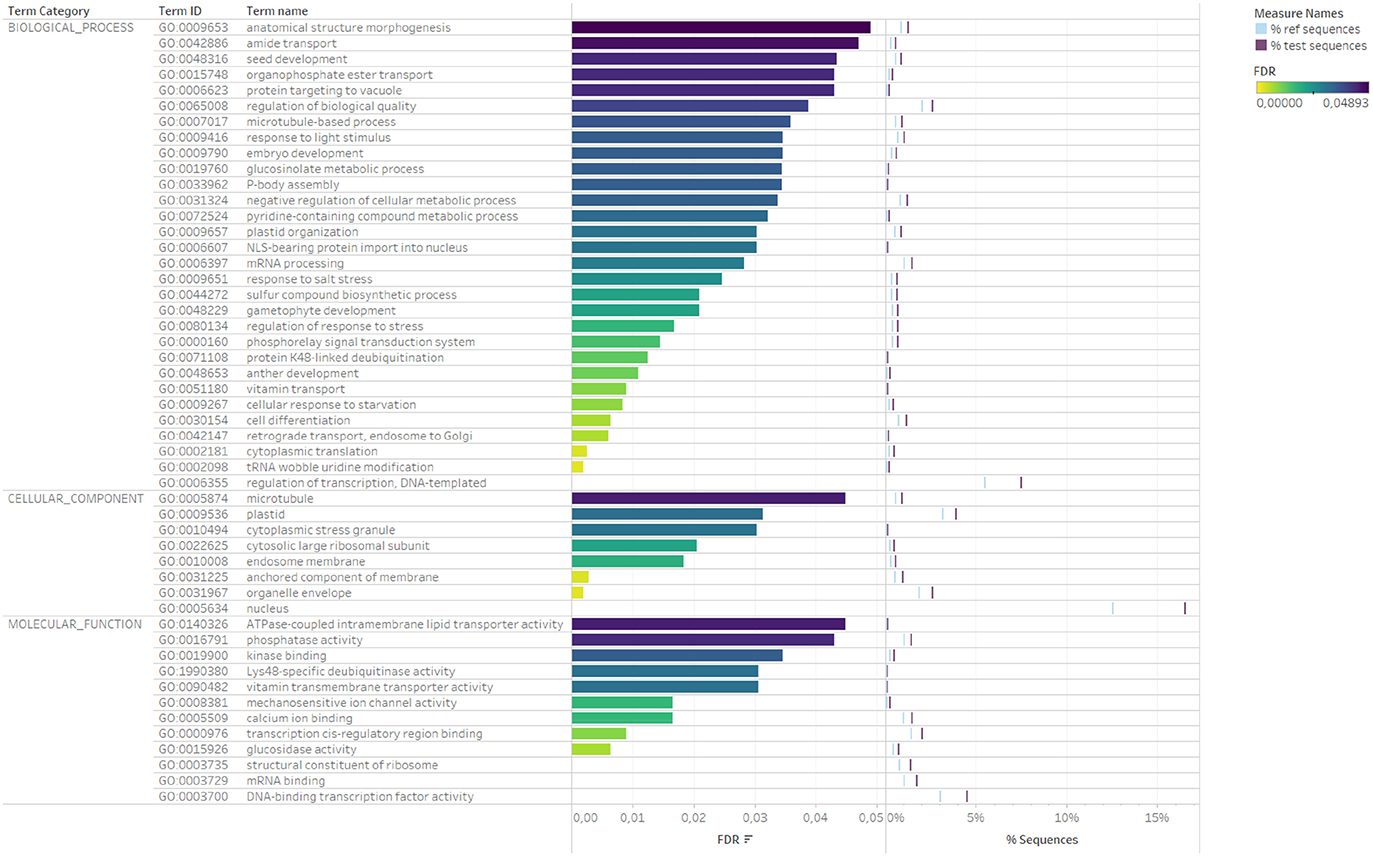
Enriched gene ontology (GO) terms for inverted and translocated gene alignment blocks. The enriched GO terms are divided by the three main GO categories. FDR shows the false discovery rate of the enriched terms. The percentage sequences found in the gene blocks are indicated for the reference (blue) and test gene (purple) sets.

## Supplementary Information

### Supplementary Note 1: Hap-mer based phasing completeness assessment

We found that separation of long-reads into haplotype bins before genome assembly resulted in splitting of the long-reads almost equally into *E. urophylla* and *E. grandis* haplotype bins (Supplementary Table 3). To further validate that long-reads were separated into the correct haplotype bins, and that the haplogenome assemblies contained mostly a single haplotype, we performed independent assessment of the haplotype specific k-mers contained within each haplogenome assembly and whether those correspond to the parent specific k-mers identified prior to trio-binning which was used for separation of long-reads into haplotype bins by Canu. We used Merqury v1.1 (Rhie *et al*., 2020) to further validate whether separation of long-reads into *E. urophylla* and *E. grandis* haplotype bins was successful. Using the parental haplo-genome assemblies we could estimate the inherited hap-mers for the child (i.e. haplotype specific k-mers present in the F_1_ haplogenome bins) to assess how well phased the assembled haplogenome assemblies are (Rhie *et al*., 2020). Using the parent specific hap-mers, we determined phase blocks (a consistent set of markers originating from a single haplotype) based on observed haplotype markers within the haplogenome assemblies with Merqury. We observed a block N_50_ and average block size of 42.45 Mb and 491.75 kb for the *E. urophylla* haplogenome assembly (Supplementary Table 11). In addition, using a maximum of 100 consecutive haplotype marker switches per phase-block window of 20 kb, we showed that the *E. urophylla* haplogenome assembly had a low switch error rate of 0.033% per block (Supplementary Table 11). In comparison, the *E. grandis* haplogenome assembly had a slightly larger block N_50_ size (43.82 Mb) and a smaller average block size (432.93 kb) with a lower switch error rate of 0.028% (Supplementary Table 11). As some short-range switches may be missed when allowing 100 consecutive switches per 20 kb phase block, we also tested phase block continuity by setting a more stringent parameter of only allowing ten switched per 20 kb block window. This resulted in an even lower phase switch error rate (0.025% and 0.020% for *E. urophylla* and *E. grandis,* respectively), even though the phase block sizes were smaller (average block N_50_ of 1.65 Mb and 2.37 Mb, respectively). These results further confirm that the long reads were separated into the correct haplotype bins and that there are few switch errors within our haplogenome assemblies.

*E. urophylla* hap-mer markers found in the *E. urophylla* haplogenome assembly and *E. grandis* hap-mer markers found in the *E. grandis* haplogenome assembly and few contaminating markers from the alternative haplogenome (Supplementary Table 11). This is reflected in the blob plot (Supplementary Figure 11), where the blob represents contigs/scaffolds, the blob size the size of the contig/scaffold, blob colour represents the parental hap-mer to which the blob belongs and how close the blob is to the x- or y-axis represents the assembly in which the hap-mer was found (Rhie *et al*., 2020). As expected, almost all blobs are close to one of the axes, with the colours matching that of the haplogenome it belongs to (red blobs of *E. urophylla* are close to the *E. urophylla* haplogenome axis, and blue *E. grandis* blobs are close to the *E. grandis* haplogenome axis; Supplementary Figure 11). This is expected due to the high level of heterozygosity within *Eucalyptus* (estimated with GenomeScope to be within a range of 1.62% to 3.6%, Supplementary Figure 12), which means that most k-mers in the offspring are actually parental hap-mers (Rhie *et al*., 2020). The high heterozygosity estimates enhance haplotype separation based on trio-binning with Canu, and, together with the results from Merqury, confirms that haplotype separation was highly successful and accurate.

Successful haplotype separation is further evidenced by Supplementary Figure 12A, where phase blocks that originate from the wrong haplogenome assembly cannot be seen when 100 and ten hap-mer marker switches are allowed per 20 kb block window. This supports that contigs likely contain markers from only one haplotype. In addition, when plotting the size of the phased blocks and contigs together, phase blocks were larger than contigs and, when plotting the size of phased blocks and scaffolds together, phase blocks were the same size as scaffolds showing good phasing performance (Supplementary Figure 12C and D) when 100 switches are allowed per 20 kb block. In comparison, when allowing only 10 switches per block, phase blocks have sizes similar to those of the contigs, indicating phase continuity within contigs. Together, these results suggest that Trio-binning with Canu was successful and have resulted in a highly phased haplogenome assembly for *E. grandis* and *E. urophylla*.

**Supplementary Table 11.** Phase block statistics of the *E. grandis* and *E. urophylla* haplo-genome assemblies. Switch error rates are also shown. The number of switch errors per 20 kb are indicated in the phase block column (10 or 100 errors per 20 kb).

| Phase blocks | Num. of blocks | Block sum (assembly size, bp) | Smallest block size (bp) | Avg. block size (bp) | Block N50 size (bp) | Longest block size (bp) | Num. of parent specific k-mers from the other haplotype | Total num. of parent specific k-mers in blocks | Switch (%) |
| --- | --- | --- | --- | --- | --- | --- | --- | --- | --- |
| *E. grandis* haplogenome 100/20 kb | 1,308 | 566,274,335 | 21 | 432,931 | 43,818,674 | 63,773,254 | 34,846 | 126,230,267 | 0.027% |
| *E. urophylla* haplogenome 100/20 kb | 1,106 | 543,874,933 | 21 | 491,750 | 42,454,739 | 60,186,461 | 39,881 | 119,331,039 | 0.033% |
| *E. grandis* haplogenome 10/20 kb | 2,300 | 566,082,275 | 21 | 246,123 | 2,374,034 | 10,482,862 | 25,382 | 126,230,267 | 0.020% |
| *E. urophylla* haplogenome 10/20 kb | 2,420 | 543,681,397 | 21 | 224,662 | 1,646,325 | 8,790,625 | 30,006 | 119,331,037 | 0.025% |

**Supplementary Figure 11.**
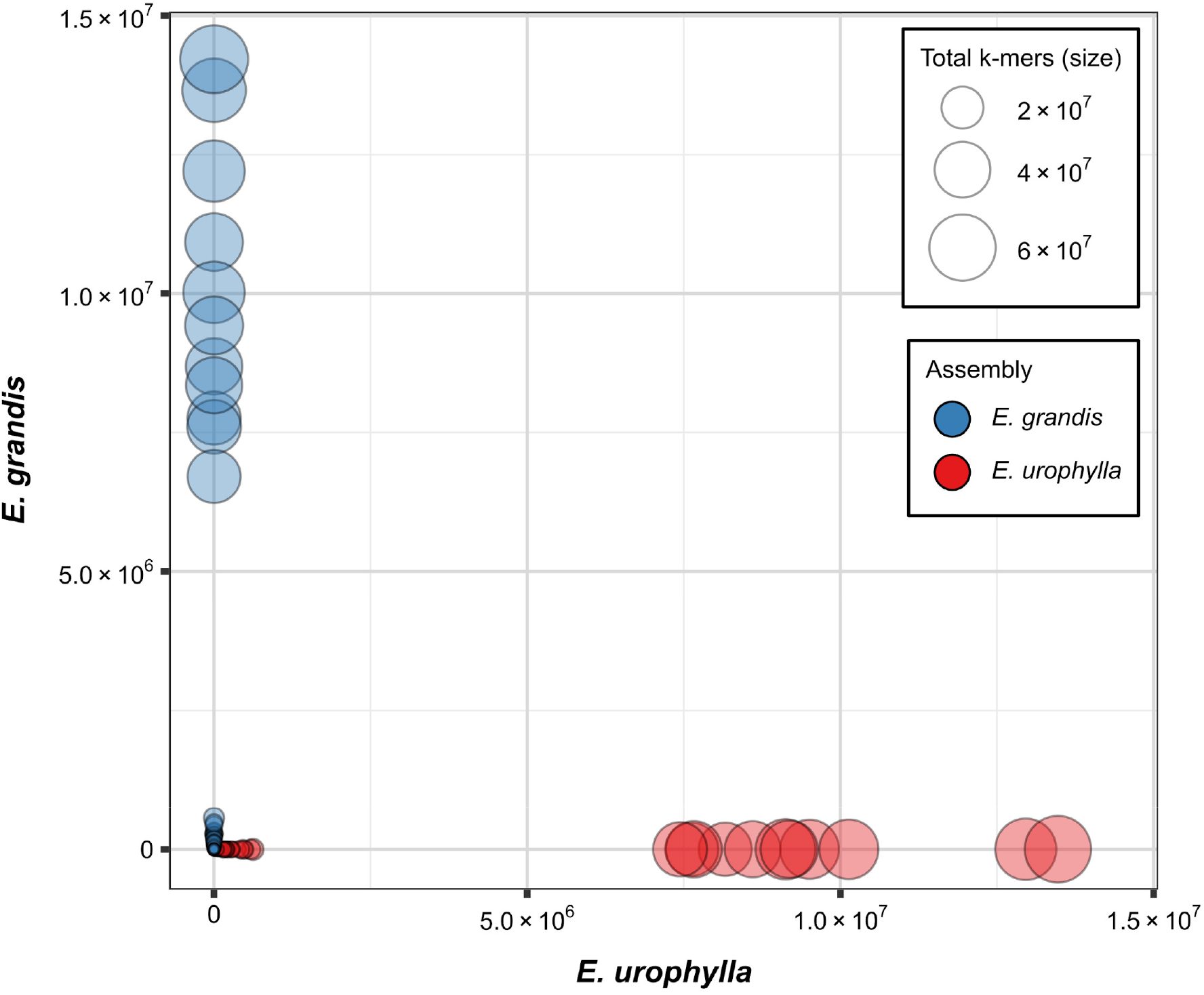
Hap-mer blob plot of the *E. grandis* and *E. urophylla* haplogenome assemblies. All hap-mer information was generated with Merqury v1.1 (Rhie *et al*., 2020). *E. urophylla* haplogenome contigs are represented by red blobs and *E. grandis* haplogenome contigs are represented by blue blobs. Blob size and contig size are proportional. The number of *E. grandis* (y-axis) and *E. urophylla* (x-axis) hap-mers are plotted per blob/contig. There are almost no *E. grandis* specific k-mers found in the *E. urophylla* assembly, while *E. urophylla* specific k-mers are found in the *E. urophylla* haplogenome assembly.

**Supplementary Figure 12.**
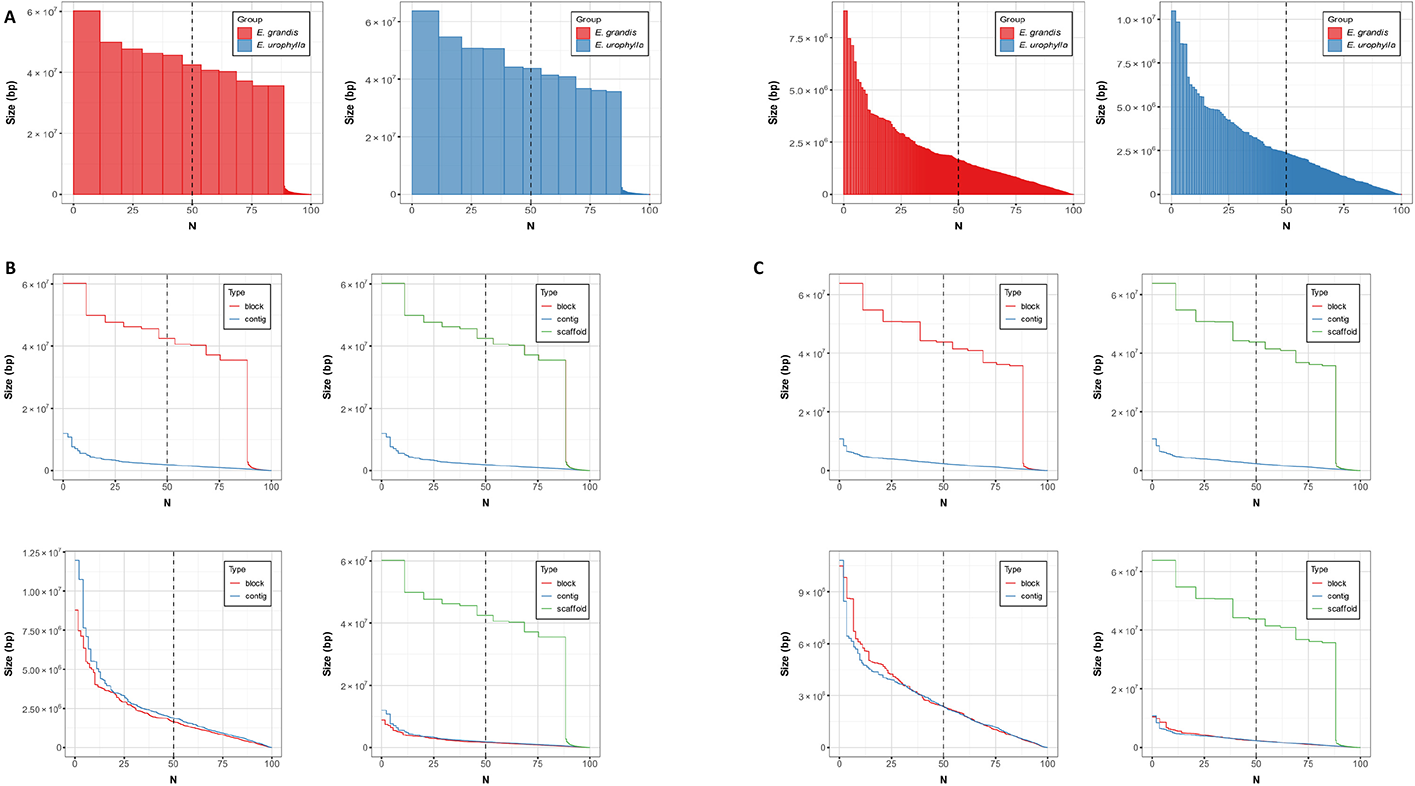
Evaluation of haplotype phase blocks. All hap-mer information was generated with Merqury v1.1 (Rhie *et al*., 2020). (A) Size sorted phase block N plots of the *E. urophylla* (red) and *E. grandis* (blue) haplogenome assemblies for 100 (left) and 10 (right) switch errors per 20 kb phase block. N shows the percentage of genome size covered by phase blocks of this size and larger are indicated on the x-axis, where the y-axis gives the block size. Blocks from the wrong haplotype are very small and are absent (too small to be seen). (B and C) Phase block N plots show the continuity of the *E. urophylla* (B) and *E. grandis* (C) haplogenome assemblies (100 and 10 switch errors allowed per 20 kb on the top and bottom respectively).

### Supplementary Note 2: Read and assembly alignment and validation of high peak content

To validate that the lower assembly sizes observed in our study were not due to genomic regions that were missing due to low or no long-read sequence coverage, we aligned 150 bp PE short-read sequencing data of the *E. grandis* and *E. urophylla* parents, binned long-read sequencing data and our haplogenome assemblies (contigs) to the *E. grandis* v2.0 reference genome (coverage was calculated as normalized reads per kilobase per million mapped reads and visualised in bins of 100 kb, Supplementary Figure 13A). Short-read sequencing data had a mapping rate of 94.34%, 95.07% and 94.78% for the *E. urophylla* and *E. grandis* parent as well as the F_1_ hybrid (Supplementary Table 1). Long-read sequencing data had a mapping rate of 99.45% (Supplementary Table 2), showing that almost all short- and long-reads mapped to the *E. grandis* v2.0 reference genome. There were no bins with zero sequence coverage, suggesting that the entire genome was sequenced, and that the smaller assembly size was not due to low or no long-read sequence coverage. We noted that there were some bins that had very high sequence coverage, and that some of these regions included mitochondrial and chloroplast sequences, which is expected as there is organellar sequence introgression into the nuclear genome of *Eucalyptus* (Pinard *et al*., 2019). However, we cannot distinguish between reads mapping to introgressed regions and those that are derived from organellar genomes. Another potential cause for bins with high sequence coverage could be repeat elements within that region. To further evaluate the nature of the sequences within high coverage bins, we extracted the bin sequences from the *E. grandis* v2.0 reference genome where genome coverage was above the total bin average within high coverage bins. We used the extracted sequences to identify their origin as either organellar or repetitive, by searching them against a blast database created from the mitochondrial and chloroplast genomes (Pinard *et al*., 2019), or by performing repeat element identification with RepeatModeler and RepeatMasker as previously described in the methods and materials.

We found that organellar introgression is indeed responsible for some of the high coverage bins (Supplementary Table 12). This was expected as Pinard *et al*., (2019) showed that there is significant introgression of organellar DNA into many regions of the nuclear genome of *E. grandis.* In particular, the high coverage bin on chromosome 9 is due to organellar DNA introgression in *E. grandis* and *E. urophylla* (Supplementary Table 12 and Supplementary Figure 14), however whether their origin is ancestral or shared still needs to be explored. This is as expected as Pinard *et al*., (2019) also found multiple introgression events in chromosome 9. Although introgression of organellar DNA explains some of the high coverage bins we have observed, the majority of identified high coverage bins contained repetitive elements, many of which are rRNA elements from the rnd-2 repeat class (Supplementary Table 12 and Supplementary Figure 14). There were also only two repeat family classes on a single chromosome (Supplementary Table 12 and Supplementary Figure 14). In conclusion, bins with high coverage are mostly the result of the high number of repeat elements found within then, with the exception of chromosome 9, which is due to organellar introgression.

**Supplementary Table 12.** *E. grandis* and *E. urophylla* high coverage bin content. A summary of the blast and RepeatMasker results is given for the genomic sequences in high *E. grandis* v2.0 genome coverage bins. The genomic regions are given (chromosome followed by the sequence position) as Query seqid, and the repeat element or mitochondrial or chloroplast sequence as the Subject seqid. Results are sorted by chromosome followed by the sequence positions. NC_040010.1 is mitochondrial genome sequences and MG925369.1 are chloroplast genome sequences. Mitochondrial, chloroplast, repeat family/class are indicated in the Type column.

| Query seqid | Query start | Query end | Subject seqid | Type | Subject start | Subject end |
| --- | --- | --- | --- | --- | --- | --- |
| *E. grandis* |  |  |  |  |  |  |
| Chr01:27900113-27901812 | 818 | 945 | NC_040010.1 | mitochondrial | 1528 | 1400 |
| Chr01:27904946-27909398 | 2 | 4450 | rnd-2_family-40 | Unknown | 1 | 4456 |
| Chr01:27905124-27906277 | 1 | 1153 | rnd-2_family-40 | Unknown | 179 | 1341 |
| Chr01:27906500-27907809 | 1 | 1309 | rnd-2_family-40 | Unknown | 1561 | 2869 |
| Chr01:27907885-27909052 | 1 | 1167 | rnd-2_family-40 | Unknown | 2947 | 4112 |
| Chr01:27910186-27912824 | 306 | 1078 | rnd-2_family-40 | Unknown | 1851 | 3272 |
| Chr01:27910186-27912824 | 1063 | 2572 | rnd-2_family-40 | Unknown | 2358 | 3409 |
| Chr01:27910756-27910861 | 1 | 105 | rnd-2_family-40 | Unknown | 2100 | 2202 |
| Chr01:27910945-27911345 | 1 | 298 | rnd-2_family-40 | Unknown | 2932 | 3243 |
| Chr01:27910945-27911345 | 304 | 400 | rnd-2_family-40 | Unknown | 2262 | 2357 |
| Chr01:27911384-27912795 | 5 | 1374 | rnd-2_family-40 | Unknown | 2358 | 3409 |
| Chr01:27912844-27915201 | 67 | 161 | rnd-2_family-40 | Unknown | 572 | 662 |
| Chr01:27912844-27916498 | 67 | 161 | rnd-2_family-40 | Unknown | 572 | 662 |
| Chr01:27929727-27932852 | 1590 | 3123 | rnd-2_family-40 | Unknown | 1 | 1544 |
| Chr01:27930207-27932411 | 1110 | 2202 | rnd-2_family-40 | Unknown | 1 | 1101 |
| Chr01:27932461-27933315 | 1 | 583 | rnd-2_family-40 | Unknown | 1154 | 1745 |
| Chr01:27932461-27933315 | 582 | 854 | NC_040010.1 | mitochondrial | 152314 | 152582 |
| Chr01:27932894-27933012 | 1 | 118 | rnd-2_family-40 | Unknown | 1590 | 1707 |
| Chr01:27933390-27936405 | 356 | 1235 | rnd-2_family-40 | Unknown | 1713 | 3237 |
| Chr01:27933390-27936405 | 1242 | 1354 | rnd-2_family-40 | Unknown | 2260 | 2373 |
| Chr01:27933390-27936405 | 1354 | 2854 | rnd-2_family-40 | Unknown | 2358 | 3546 |
| Chr01:27933390-27936405 | 2911 | 3014 | rnd-2_family-40 | Unknown | 4352 | 4456 |
| Chr01:27933787-27935996 | 1 | 838 | rnd-2_family-40 | Unknown | 1755 | 3237 |
| Chr01:27933787-27935996 | 845 | 957 | rnd-2_family-40 | Unknown | 2260 | 2373 |
| Chr01:27933787-27935996 | 957 | 2209 | rnd-2_family-40 | Unknown | 2358 | 3298 |
| Chr01:27936076-27936202 | 1 | 126 | rnd-2_family-40 | Unknown | 3379 | 3504 |
| Chr01:27936428-27939805 | 814 | 958 | rnd-2_family-40 | Unknown | 560 | 702 |
| Chr01:27936428-27939805 | 3244 | 3361 | rnd-2_family-40 | Unknown | 2396 | 2513 |
| Chr01:27936999-27937661 | 243 | 387 | rnd-2_family-40 | Unknown | 560 | 702 |
| Chr01:27939234-27939742 | 438 | 508 | rnd-2_family-40 | Unknown | 2396 | 2466 |
| Chr01:27939817-27944583 | 1 | 1842 | rnd-2_family-40 | Unknown | 2605 | 4456 |
| Chr01:27939817-27944583 | 2899 | 3668 | rnd-2_family-40 | Unknown | 1851 | 3272 |
| Chr01:27939817-27944583 | 3653 | 4365 | rnd-2_family-40 | Unknown | 6 | 685 |
| Chr01:27939817-27944583 | 4413 | 4632 | NC_040010.1 | mitochondrial | 152280 | 152500 |
| Chr01:27940076-27942426 | 1 | 1583 | rnd-2_family-40 | Unknown | 2872 | 4456 |
| Chr01:27940076-27942426 | 935 | 1013 | NC_040010.1 | mitochondrial | 436100 | 436179 |
| Chr01:27942542-27944575 | 174 | 923 | rnd-2_family-40 | Unknown | 1851 | 3244 |
| Chr01:27942542-27944575 | 928 | 1640 | rnd-2_family-40 | Unknown | 6 | 685 |
| Chr01:27942542-27944575 | 1688 | 1907 | NC_040010.1 | mitochondrial | 152280 | 152500 |
| Chr01:27944607-27956058 | 711 | 1391 | rnd-2_family-40 | Unknown | 2654 | 3356 |
| Chr01:27944607-27956058 | 1603 | 1709 | rnd-2_family-40 | Unknown | 560 | 662 |
| Chr01:27945169-27947611 | 149 | 829 | rnd-2_family-40 | Unknown | 2654 | 3356 |
| Chr01:27945169-27947611 | 1041 | 1147 | rnd-2_family-40 | Unknown | 560 | 662 |
| Chr01:27956077-27960535 | 672 | 3240 | rnd-2_family-40 | Unknown | 14 | 3240 |
| Chr01:27956077-27960535 | 3239 | 3351 | rnd-2_family-40 | Unknown | 2260 | 2373 |
| Chr01:27956077-27960535 | 3351 | 3600 | rnd-2_family-40 | Unknown | 86 | 330 |
| Chr01:27956118-27957607 | 631 | 1483 | rnd-2_family-40 | Unknown | 14 | 867 |
| Chr01:27957619-27960426 | 1 | 1698 | rnd-2_family-40 | Unknown | 873 | 3240 |
| Chr01:27957619-27960426 | 1697 | 1809 | rnd-2_family-40 | Unknown | 2260 | 2373 |
| Chr01:27957619-27960426 | 1809 | 2058 | rnd-2_family-40 | Unknown | 86 | 330 |
| Chr01:27960576-27965778 | 2645 | 2965 | rnd-2_family-40 | Unknown | 2358 | 2607 |
| Chr01:27962629-27963653 | 592 | 912 | rnd-2_family-40 | Unknown | 2358 | 2607 |
| Chr01:27965815-27967195 | 278 | 913 | rnd-2_family-40 | Unknown | 2684 | 3352 |
| Chr01:27965815-27967195 | 1134 | 1279 | rnd-2_family-40 | Unknown | 560 | 702 |
| Chr01:27966095-27966990 | 1 | 633 | rnd-2_family-40 | Unknown | 2687 | 3352 |
| Chr01:27967713-27975626 | 6977 | 7913 | rnd-2_family-40 | Unknown | 1 | 942 |
| Chr01:27973686-27974796 | 1004 | 1110 | rnd-2_family-40 | Unknown | 1 | 108 |
| Chr01:27975718-27976943 | 1 | 1225 | rnd-2_family-40 | Unknown | 1036 | 2259 |
| Chr01:27975965-27976224 | 1 | 259 | rnd-2_family-40 | Unknown | 1281 | 1540 |
| Chr01:27976675-27977201 | 1 | 526 | rnd-2_family-40 | Unknown | 1990 | 2264 |
| Chr01:27977261-27977344 | 1 | 83 | rnd-2_family-40 | Unknown | 2275 | 2357 |
| Chr01:27977755-27977887 | 4 | 132 | rnd-2_family-40 | Unknown | 2444 | 2572 |
| Chr01:27978212-27978291 | 1 | 79 | rnd-2_family-40 | Unknown | 2907 | 2985 |
| Chr01:27978614-27978675 | 1 | 61 | rnd-2_family-40 | Unknown | 3310 | 3370 |
| Chr01:27979737-27980327 | 189 | 334 | rnd-2_family-40 | Unknown | 560 | 702 |
| Chr01:27992448-27993267 | 158 | 819 | rnd-2_family-40 | Unknown | 1 | 663 |
| Chr01:27993781-27993980 | 1 | 199 | rnd-2_family-40 | Unknown | 1280 | 1479 |
| Chr01:27994158-27994259 | 1 | 101 | rnd-2_family-40 | Unknown | 1648 | 1748 |
| Chr01:27998719-27998772 | 1 | 53 | rnd-2_family-40 | Unknown | 264 | 316 |
| Chr02:22324794-22324881 | 1 | 81 | rnd-2_family-5 | rRNA | -235 | 899 |
| Chr02:22324894-22325061 | 2 | 167 | rnd-2_family-5 | rRNA | -346 | 788 |
| Chr02:22325073-22325232 | 3 | 159 | rnd-2_family-5 | rRNA | -477 | 657 |
| Chr02:22331577-22331666 | 1 | 89 | rnd-2_family-5 | rRNA | -413 | 721 |
| Chr02:22351050-22351099 | 1 | 49 | rnd-2_family-5 | rRNA | -236 | 898 |
| Chr02:22354417-22354496 | 1 | 79 | rnd-2_family-5 | rRNA | -500 | 634 |
| Chr02:22361244-22361395 | 1 | 151 | rnd-2_family-7 | rRNA | -1 | 2497 |
| Chr02:22361450-22361633 | 1 | 183 | rnd-2_family-7 | rRNA | -207 | 2291 |
| Chr02:22361657-22361886 | 1 | 229 | rnd-2_family-7 | rRNA | -415 | 2083 |
| Chr02:22363675-22364328 | 265 | 529 | rnd-2_family-5 | rRNA | -680 | 454 |
| Chr02:22383597-22383625 | 1 | 28 | rnd-2_family-5 | rRNA | -66 | 1068 |
| Chr02:22383636-22387348 | 843 | 956 | rnd-2_family-5 | rRNA | -253 | 881 |
| Chr02:22383636-22387348 | 1402 | 1666 | rnd-2_family-5 | rRNA | 99 | 454 |
| Chr02:22383636-22387348 | 2140 | 2201 | rnd-2_family-5 | rRNA | -992 | 142 |
| Chr02:22383636-22387348 | 2193 | 2420 | rnd-2_family-5 | rRNA | 1 | 227 |
| Chr02:22383636-22387348 | 2202 | 3014 | rnd-2_family-5 | rRNA | -360 | 774 |
| Chr02:22383636-22387348 | 2486 | 3700 | rnd-2_family-5 | rRNA | 2 | 1134 |
| Chr02:22384542-22385844 | 1 | 50 | rnd-2_family-5 | rRNA | -326 | 808 |
| Chr02:22384542-22385844 | 496 | 760 | rnd-2_family-5 | rRNA | 99 | 454 |
| Chr02:22384542-22385844 | 1234 | 1302 | rnd-2_family-5 | rRNA | -992 | 142 |
| Chr02:22385034-22385154 | 4 | 120 | rnd-2_family-5 | rRNA | 217 | 331 |
| Chr02:22385238-22385372 | 1 | 39 | rnd-2_family-5 | rRNA | 416 | 454 |
| Chr02:22385549-22385883 | 227 | 331 | rnd-2_family-5 | rRNA | -992 | 142 |
| Chr02:22386100-22387076 | 22 | 976 | rnd-2_family-5 | rRNA | 2 | 899 |
| Chr02:22387015-22387072 | 1 | 57 | rnd-2_family-5 | rRNA | 839 | 895 |
| Chr02:22387099-22387240 | 1 | 141 | rnd-2_family-5 | rRNA | 924 | 1064 |
| Chr02:22387208-22387258 | 1 | 50 | rnd-2_family-5 | rRNA | 1033 | 1082 |
| Chr02:22387272-22387338 | 1 | 64 | rnd-2_family-5 | rRNA | 1066 | 1134 |
| Chr02:22387371-22391423 | 142 | 2637 | rnd-2_family-7 | rRNA | 0 | 2498 |
| Chr02:22387510-22388473 | 3 | 963 | rnd-2_family-7 | rRNA | 0 | 2498 |
| Chr02:22387513-22388150 | 1 | 637 | rnd-2_family-7 | rRNA | -1 | 2497 |
| Chr02:22388484-22388627 | 1 | 143 | rnd-2_family-7 | rRNA | -973 | 1525 |
| Chr02:22388639-22388943 | 1 | 304 | rnd-2_family-7 | rRNA | -1128 | 1370 |
| Chr02:22388984-22389313 | 1 | 329 | rnd-2_family-7 | rRNA | -1473 | 1025 |
| Chr02:22389335-22389521 | 1 | 186 | rnd-2_family-7 | rRNA | -1824 | 674 |
| Chr02:22389538-22389836 | 1 | 298 | rnd-2_family-7 | rRNA | -2027 | 471 |
| Chr02:22398421-22401537 | 9 | 48 | rnd-2_family-5 | rRNA | -63 | 1071 |
| Chr02:22398421-22401537 | 212 | 300 | rnd-2_family-5 | rRNA | 144 | 228 |
| Chr02:22398421-22401537 | 893 | 1006 | rnd-2_family-5 | rRNA | -253 | 881 |
| Chr02:22398421-22401537 | 1452 | 1716 | rnd-2_family-5 | rRNA | 99 | 454 |
| Chr02:22398421-22401537 | 2194 | 2258 | rnd-2_family-5 | rRNA | -992 | 142 |
| Chr02:22398421-22401537 | 2247 | 3116 | rnd-2_family-5 | rRNA | 1 | 1009 |
| Chr02:22399335-22400258 | 1 | 92 | rnd-2_family-5 | rRNA | -275 | 859 |
| Chr02:22399335-22400258 | 538 | 802 | rnd-2_family-5 | rRNA | 99 | 454 |
| Chr02:22400399-22401537 | 216 | 280 | rnd-2_family-5 | rRNA | -992 | 142 |
| Chr02:22400399-22401537 | 269 | 1138 | rnd-2_family-5 | rRNA | 1 | 1009 |
| Chr02:22404442-22404583 | 101 | 141 | rnd-2_family-5 | rRNA | 756 | 799 |
| Chr02:22412412-22413943 | 1 | 1531 | rnd-2_family-7 | rRNA | 97 | 1627 |
| Chr02:22442538-22451781 | 1 | 1335 | rnd-2_family-7 | rRNA | 1038 | 2367 |
| Chr02:22442538-22451781 | 1333 | 1463 | rnd-2_family-7 | rRNA | 2327 | 2460 |
| Chr02:22442538-22451781 | 1449 | 1716 | rnd-2_family-7 | rRNA | 2254 | 2498 |
| Chr02:22442538-22451781 | 1822 | 2884 | rnd-2_family-5 | rRNA | 0 | 1134 |
| Chr02:22442538-22451781 | 3602 | 3715 | rnd-2_family-5 | rRNA | 756 | 881 |
| Chr02:22442538-22451781 | 4308 | 4396 | rnd-2_family-5 | rRNA | -906 | 228 |
| Chr02:22442538-22451781 | 4560 | 4599 | rnd-2_family-5 | rRNA | 1032 | 1071 |
| Chr02:22442538-22451781 | 7252 | 9243 | rnd-2_family-7 | rRNA | 2 | 1983 |
| Chr02:22443302-22443506 | 1 | 204 | rnd-2_family-7 | rRNA | 1802 | 2005 |
| Chr02:22443529-22444113 | 1 | 344 | rnd-2_family-7 | rRNA | 2029 | 2367 |
| Chr02:22443529-22444113 | 342 | 472 | rnd-2_family-7 | rRNA | 2327 | 2460 |
| Chr02:22443529-22444113 | 458 | 583 | rnd-2_family-7 | rRNA | 2254 | 2369 |
| Chr02:22444127-22444507 | 1 | 127 | rnd-2_family-7 | rRNA | 2372 | 2498 |
| Chr02:22444127-22444507 | 233 | 380 | rnd-2_family-5 | rRNA | 0 | 1134 |
| Chr02:22444583-22445042 | 1 | 442 | rnd-2_family-5 | rRNA | -237 | 897 |
| Chr02:22445098-22445297 | 1 | 188 | rnd-2_family-5 | rRNA | -362 | 772 |
| Chr02:22445329-22445373 | 2 | 44 | rnd-2_family-5 | rRNA | -529 | 605 |
| Chr02:22446018-22449703 | 122 | 235 | rnd-2_family-5 | rRNA | 756 | 881 |
| Chr02:22446018-22449703 | 828 | 916 | rnd-2_family-5 | rRNA | -906 | 228 |
| Chr02:22446018-22449703 | 1080 | 1119 | rnd-2_family-5 | rRNA | 1032 | 1071 |
| Chr02:22449788-22450114 | 2 | 326 | rnd-2_family-7 | rRNA | 2 | 326 |
| Chr02:22450125-22450236 | 1 | 106 | rnd-2_family-5 | rRNA | -624 | 510 |
| Chr02:22450165-22450229 | 1 | 64 | rnd-2_family-5 | rRNA | -622 | 512 |
| Chr02:22450244-22450360 | 10 | 116 | rnd-2_family-7 | rRNA | 455 | 561 |
| Chr02:22450371-22450534 | 1 | 163 | rnd-2_family-7 | rRNA | 573 | 735 |
| Chr02:22450545-22450593 | 1 | 48 | rnd-2_family-7 | rRNA | 747 | 794 |
| Chr02:22450610-22450962 | 1 | 352 | rnd-2_family-7 | rRNA | 812 | 1163 |
| Chr02:22450974-22451577 | 1 | 603 | rnd-2_family-7 | rRNA | 1176 | 1778 |
| Chr02:22451596-22451781 | 1 | 185 | rnd-2_family-7 | rRNA | 1798 | 1983 |
| Chr02:22454416-22455567 | 6 | 217 | rnd-2_family-5 | rRNA | -903 | 231 |
| Chr02:22454416-22455567 | 125 | 270 | rnd-2_family-5 | rRNA | 12 | 142 |
| Chr02:22454416-22455567 | 743 | 1007 | rnd-2_family-5 | rRNA | -680 | 454 |
| Chr02:22461592-22462381 | 1 | 376 | rnd-2_family-7 | rRNA | 2096 | 2460 |
| Chr02:22461592-22462381 | 362 | 612 | rnd-2_family-7 | rRNA | 2254 | 2498 |
| Chr02:22461592-22462381 | 627 | 696 | rnd-2_family-5 | rRNA | -449 | 685 |
| Chr02:22461592-22462381 | 692 | 780 | rnd-2_family-5 | rRNA | 1038 | 1117 |
| Chr02:22461592-22462381 | 738 | 789 | rnd-2_family-5 | rRNA | 0 | 1134 |
| Chr04:1305-5155 | 1 | 3850 | rnd-2_family-2 | Unknown | 1 | 703 |
| Chr07:28766528-28775042 | 2103 | 3671 | NC_040010.1 | mitochondrial | 1577 | 1 |
| Chr07:28767636-28767757 | 1 | 121 | NC_040010.1 | mitochondrial | 1296 | 1175 |
| Chr07:28767777-28770565 | 854 | 2422 | NC_040010.1 | mitochondrial | 1577 | 1 |
| Chr07:28770856-28775023 | 2104 | 3670 | NC_040010.1 | mitochondrial | 1577 | 1 |
| Chr07:28775105-28779796 | 2513 | 4080 | NC_040010.1 | mitochondrial | 1577 | 1 |
| Chr07:28779940-28780380 | 1 | 440 | NC_040010.1 | mitochondrial | 530 | 91 |
| Chr08:3021-5525 | 2 | 2504 | rnd-2_family-2 | Unknown | 6 | 696 |
| Chr08:436-3002 | 3 | 2566 | rnd-2_family-2 | Unknown | 6 | 703 |
| Chr08:5539-8320 | 52 | 2778 | rnd-2_family-2 | Unknown | 6 | 703 |
| Chr09:28439306-28439677 | 1 | 371 | NC_040010.1 | mitochondrial | 269327 | 269697 |
| Chr09:28444416-28445106 | 1 | 690 | NC_040010.1 | mitochondrial | 274439 | 275128 |
| Chr09:28446035-28446150 | 1 | 115 | NC_040010.1 | mitochondrial | 276058 | 276172 |
| Chr09:28446331-28447025 | 1 | 694 | NC_040010.1 | mitochondrial | 276354 | 277045 |
| Chr09:28447432-28447515 | 1 | 83 | NC_040010.1 | mitochondrial | 277453 | 277535 |
| Chr09:28447697-28447937 | 1 | 240 | NC_040010.1 | mitochondrial | 277718 | 277957 |
| Chr09:28455592-28455658 | 1 | 66 | NC_040010.1 | mitochondrial | 285623 | 285688 |
| Chr09:28455675-28455819 | 1 | 144 | NC_040010.1 | mitochondrial | 285706 | 285849 |
| Chr09:28456407-28456820 | 1 | 413 | NC_040010.1 | mitochondrial | 286438 | 286850 |
| Chr09:28458096-28458505 | 1 | 409 | NC_040010.1 | mitochondrial | 288128 | 288536 |
| Chr09:28470448-28471709 | 1 | 1261 | MG925369.1 | chloroplast | 100652 | 101912 |
| Chr09:28472440-28472983 | 1 | 543 | MG925369.1 | chloroplast | 102643 | 103185 |
| Chr09:28473031-28473130 | 1 | 99 | MG925369.1 | chloroplast | 103234 | 103332 |
| Chr09:28473262-28473993 | 1 | 731 | MG925369.1 | chloroplast | 103465 | 104195 |
| Chr09:28473262-28473993 | 569 | 731 | NC_040010.1 | mitochondrial | 476331 | 476493 |
| Chr09:28474042-28474263 | 1 | 221 | MG925369.1 | chloroplast | 104245 | 104465 |
| Chr09:28474042-28474263 | 1 | 221 | NC_040010.1 | mitochondrial | 476543 | 476763 |
| Chr09:28474377-28475057 | 1 | 680 | MG925369.1 | chloroplast | 104580 | 105259 |
| Chr09:28474377-28475057 | 1 | 680 | NC_040010.1 | mitochondrial | 476878 | 477557 |
| Chr09:28475081-28475313 | 1 | 232 | MG925369.1 | chloroplast | 105284 | 105515 |
| Chr09:28475081-28475313 | 8 | 232 | NC_040010.1 | mitochondrial | 477690 | 477914 |
| Chr09:28477909-28478023 | 1 | 114 | MG925369.1 | chloroplast | 107058 | 107171 |
| Chr09:28477909-28478023 | 1 | 114 | NC_040010.1 | mitochondrial | 123515 | 123628 |
| Chr09:28478200-28479134 | 321 | 934 | NC_040010.1 | mitochondrial | 291070 | 291683 |
| Chr09:28484219-28484629 | 1 | 410 | NC_040010.1 | mitochondrial | 296770 | 297179 |
| Chr09:28485755-28485968 | 1 | 213 | NC_040010.1 | mitochondrial | 298369 | 298581 |
| Chr09:28486279-28486454 | 1 | 175 | NC_040010.1 | mitochondrial | 298893 | 299067 |
| Chr09:28488520-28488772 | 1 | 252 | NC_040010.1 | mitochondrial | 301135 | 301386 |
| Chr09:28489954-28490880 | 1 | 926 | NC_040010.1 | mitochondrial | 302571 | 303496 |
| Chr09:28491392-28491687 | 1 | 295 | NC_040010.1 | mitochondrial | 304057 | 304351 |
| Chr09:28494819-28495465 | 1 | 646 | NC_040010.1 | mitochondrial | 307394 | 308039 |
| Chr09:28495963-28496584 | 1 | 621 | NC_040010.1 | mitochondrial | 308538 | 309158 |
| Chr09:28497280-28497483 | 1 | 203 | NC_040010.1 | mitochondrial | 309855 | 310057 |
| Chr09:28497548-28497837 | 67 | 289 | NC_040010.1 | mitochondrial | 310248 | 310470 |
| Chr09:28498452-28498808 | 1 | 356 | NC_040010.1 | mitochondrial | 311086 | 311441 |
| Chr09:28498846-28499532 | 1 | 686 | NC_040010.1 | mitochondrial | 311480 | 312165 |
| Chr10:32730112-32730311 | 1 | 199 | MG925369.1 | chloroplast | 100704 | 100506 |
| Chr10:32730391-32730696 | 1 | 305 | MG925369.1 | chloroplast | 100424 | 100120 |
| Chr10:32730776-32730963 | 1 | 187 | MG925369.1 | chloroplast | 99702 | 99888 |
| Chr10:32732160-32732296 | 4 | 136 | MG925369.1 | chloroplast | 32644 | 32787 |
| Chr10:32732160-32732296 | 4 | 136 | NC_040010.1 | mitochondrial | 159194 | 159051 |
| Chr10:32745461-32745819 | 1 | 358 | MG925369.1 | chloroplast | 89122 | 89479 |
| Chr10:32745461-32745819 | 1 | 358 | NC_040010.1 | mitochondrial | 23775 | 23418 |
| Chr10:32749232-32749415 | 1 | 183 | MG925369.1 | chloroplast | 57977 | 58159 |
| Chr10:32753395-32753617 | 1 | 222 | MG925369.1 | chloroplast | 27696 | 27917 |
| Chr10:32765140-32765194 | 1 | 54 | MG925369.1 | chloroplast | 23078 | 23025 |
| Chr10:32769304-32769430 | 1 | 36 | MG925369.1 | chloroplast | 90646 | 90681 |
| Chr10:32769304-32769430 | 1 | 36 | NC_040010.1 | mitochondrial | 22251 | 22216 |
| Chr10:51819-51891 | 1 | 72 | rnd-2_family-2 | Unknown | 73 | 144 |
| Chr10:52214-52263 | 1 | 49 | rnd-2_family-2 | Unknown | 102 | 150 |
| Chr10:52488-52632 | 1 | 144 | rnd-2_family-2 | Unknown | 216 | 360 |
| Chr10:54026-54861 | 3 | 835 | rnd-2_family-2 | Unknown | 1 | 703 |
| Chr10:55576-56233 | 2 | 651 | rnd-2_family-2 | Unknown | 52 | 702 |
| Chr10:61671-78127 | 1 | 4864 | rnd-2_family-2 | Unknown | 1 | 703 |
| Chr10:61671-78127 | 4969 | 16452 | rnd-2_family-2 | Unknown | 1 | 703 |
| Chr10:61671-78129 | 1 | 4864 | rnd-2_family-2 | Unknown | 1 | 703 |
| Chr10:61671-78129 | 4969 | 16452 | rnd-2_family-2 | Unknown | 1 | 703 |
| Chr10:61761-61924 | 1 | 163 | rnd-2_family-2 | Unknown | 539 | 701 |
| Chr10:61961-62286 | 1 | 325 | rnd-2_family-2 | Unknown | 189 | 514 |
| Chr10:62325-62681 | 2 | 356 | rnd-2_family-2 | Unknown | 189 | 544 |
| Chr10:62739-63028 | 1 | 287 | rnd-2_family-2 | Unknown | 236 | 524 |
| Chr10:63056-63212 | 1 | 156 | rnd-2_family-2 | Unknown | 387 | 542 |
| Chr10:63281-63590 | 1 | 309 | rnd-2_family-2 | Unknown | 260 | 569 |
| Chr10:63638-63731 | 1 | 93 | rnd-2_family-2 | Unknown | 68 | 160 |
| Chr10:63795-64151 | 1 | 356 | rnd-2_family-2 | Unknown | 42 | 395 |
| Chr10:64193-64272 | 1 | 79 | rnd-2_family-2 | Unknown | 71 | 149 |
| Chr10:64300-64402 | 1 | 102 | rnd-2_family-2 | Unknown | 362 | 463 |
| Chr10:64433-64922 | 1 | 489 | rnd-2_family-2 | Unknown | 129 | 618 |
| Chr10:64946-65011 | 1 | 65 | rnd-2_family-2 | Unknown | 460 | 524 |
| Chr10:65050-65379 | 1 | 323 | rnd-2_family-2 | Unknown | 381 | 703 |
| Chr10:65401-65564 | 1 | 163 | rnd-2_family-2 | Unknown | 365 | 527 |
| Chr10:65663-66438 | 1 | 775 | rnd-2_family-2 | Unknown | 1 | 703 |
| Chr10:66672-66755 | 1 | 83 | rnd-2_family-2 | Unknown | 75 | 157 |
| Chr10:66786-67110 | 1 | 324 | rnd-2_family-2 | Unknown | 204 | 528 |
| Chr10:67174-67323 | 1 | 149 | rnd-2_family-2 | Unknown | 43 | 191 |
| Chr10:67387-67645 | 1 | 258 | rnd-2_family-2 | Unknown | 440 | 697 |
| Chr10:67679-67795 | 1 | 116 | rnd-2_family-2 | Unknown | 1 | 116 |
| Chr10:67813-68074 | 1 | 260 | rnd-2_family-2 | Unknown | 443 | 703 |
| Chr10:68184-68273 | 1 | 89 | rnd-2_family-2 | Unknown | 95 | 182 |
| Chr10:68419-68890 | 1 | 471 | rnd-2_family-2 | Unknown | 146 | 618 |
| Chr10:69132-69213 | 1 | 80 | rnd-2_family-2 | Unknown | 253 | 332 |
| Chr10:69232-69296 | 1 | 64 | rnd-2_family-2 | Unknown | 168 | 231 |
| Chr10:69322-69459 | 1 | 137 | rnd-2_family-2 | Unknown | 75 | 211 |
| Chr10:69504-69720 | 2 | 216 | rnd-2_family-2 | Unknown | 75 | 289 |
| Chr10:69895-70035 | 1 | 140 | rnd-2_family-2 | Unknown | 99 | 238 |
| Chr10:70366-70526 | 1 | 160 | rnd-2_family-2 | Unknown | 75 | 234 |
| Chr10:70689-71002 | 1 | 313 | rnd-2_family-2 | Unknown | 215 | 528 |
| Chr10:71096-71396 | 1 | 300 | rnd-2_family-2 | Unknown | 75 | 375 |
| Chr10:71429-71626 | 1 | 197 | rnd-2_family-2 | Unknown | 409 | 605 |
| Chr10:71835-71993 | 1 | 158 | rnd-2_family-2 | Unknown | 455 | 612 |
| Chr10:72056-72107 | 1 | 51 | rnd-2_family-2 | Unknown | 126 | 176 |
| Chr10:72214-72365 | 1 | 151 | rnd-2_family-2 | Unknown | 102 | 252 |
| Chr10:72387-72522 | 1 | 135 | rnd-2_family-2 | Unknown | 92 | 227 |
| Chr10:72621-72806 | 1 | 185 | rnd-2_family-2 | Unknown | 328 | 512 |
| Chr10:73043-73341 | 1 | 297 | rnd-2_family-2 | Unknown | 384 | 680 |
| Chr10:73419-73542 | 1 | 122 | rnd-2_family-2 | Unknown | 577 | 698 |
| Chr10:73575-74282 | 2 | 703 | rnd-2_family-2 | Unknown | 1 | 703 |
| Chr10:74443-75237 | 1 | 794 | rnd-2_family-2 | Unknown | 1 | 703 |
| Chr10:75306-76368 | 1 | 1062 | rnd-2_family-2 | Unknown | 1 | 703 |
| Chr10:76394-76479 | 1 | 85 | rnd-2_family-2 | Unknown | 75 | 158 |
| Chr10:76579-77019 | 1 | 440 | rnd-2_family-2 | Unknown | 75 | 515 |
| Chr10:77079-77366 | 1 | 287 | rnd-2_family-2 | Unknown | 393 | 679 |
| Chr10:77408-77805 | 1 | 397 | rnd-2_family-2 | Unknown | 172 | 568 |
| Chr10:77892-77945 | 1 | 53 | rnd-2_family-2 | Unknown | 106 | 158 |
| Chr10:77996-78035 | 1 | 39 | rnd-2_family-2 | Unknown | 210 | 248 |
| Chr11:24024757-24026068 | 1146 | 1251 | rnd-2_family-45 | Unknown | 379 | 484 |
| Chr11:24024757-24026068 | 1269 | 1310 | rnd-2_family-45 | Unknown | 474 | 515 |
| Chr11:24028609-24032432 | 2 | 41 | rnd-2_family-45 | Unknown | 445 | 484 |
| Chr11:24028609-24032432 | 48 | 577 | rnd-2_family-45 | Unknown | 761 | 1379 |
| Chr11:24042370-24042566 | 146 | 196 | rnd-2_family-45 | Unknown | 1 | 51 |
| Chr11:24042680-24043121 | 1 | 441 | rnd-2_family-45 | Unknown | 166 | 609 |
| Chr11:24043199-24047383 | 1 | 694 | rnd-2_family-45 | Unknown | 688 | 1382 |
| Chr11:24047445-24049188 | 1619 | 1743 | rnd-2_family-45 | Unknown | 1 | 125 |
| Chr11:24049219-24049802 | 1 | 583 | rnd-2_family-45 | Unknown | 199 | 781 |
| Chr11:24049856-24051712 | 1 | 538 | rnd-2_family-45 | Unknown | 840 | 1382 |
| Chr11:24070224-24071609 | 2 | 1382 | rnd-2_family-45 | Unknown | 1 | 1382 |
| Chr11:24080542-24083870 | 1 | 522 | rnd-2_family-45 | Unknown | 412 | 967 |
| Chr11:24080542-24083870 | 540 | 870 | rnd-2_family-45 | Unknown | 1054 | 1376 |
| Chr11:24084573-24085756 | 358 | 444 | rnd-2_family-45 | Unknown | 395 | 484 |
| Chr11:24084573-24085756 | 451 | 980 | rnd-2_family-45 | Unknown | 761 | 1379 |
| Chr11:24096117-24098112 | 863 | 1995 | rnd-2_family-45 | Unknown | 1 | 1128 |
| *E. urophylla* |  |  |  |  |  |  |
| Chr02:22324906-22325221 | 3 | 314 | rnd-2_family-5 | rRNA | -87 | 4605 |
| Chr02:22350942-22351100 | 1 | 131 | rnd-2_family-5 | rRNA | 4236 | 4364 |
| Chr02:22354435-22354497 | 1 | 61 | rnd-2_family-5 | rRNA | 4236 | 4296 |
| Chr02:22355414-22355852 | 4 | 73 | rnd-2_family-5 | rRNA | 1205 | 1280 |
| Chr02:22361242-22361405 | 1 | 163 | rnd-2_family-5 | rRNA | -780 | 3912 |
| Chr02:22361452-22361929 | 1 | 477 | rnd-2_family-5 | rRNA | -996 | 3696 |
| Chr02:22363675-22364401 | 305 | 358 | rnd-2_family-5 | rRNA | 1220 | 1279 |
| Chr02:22363675-22364401 | 362 | 431 | rnd-2_family-5 | rRNA | 1205 | 1280 |
| Chr02:22383595-22387348 | 1 | 31 | rnd-2_family-5 | rRNA | 4178 | 4208 |
| Chr02:22383595-22387348 | 1541 | 1610 | rnd-2_family-5 | rRNA | -3412 | 1280 |
| Chr02:22383595-22387348 | 1614 | 1667 | rnd-2_family-5 | rRNA | -3413 | 1279 |
| Chr02:22383595-22387348 | 2224 | 2352 | rnd-2_family-5 | rRNA | 4571 | 4691 |
| Chr02:22383595-22387348 | 2269 | 2723 | rnd-2_family-5 | rRNA | 0 | 4692 |
| Chr02:22383595-22387348 | 2684 | 3413 | rnd-2_family-5 | rRNA | 4236 | 4686 |
| Chr02:22383595-22387348 | 3103 | 3753 | rnd-2_family-5 | rRNA | 0 | 4692 |
| Chr02:22383826-22385890 | 1310 | 1379 | rnd-2_family-5 | rRNA | -3412 | 1280 |
| Chr02:22383826-22385890 | 1383 | 1436 | rnd-2_family-5 | rRNA | -3413 | 1279 |
| Chr02:22383826-22385890 | 1995 | 2064 | rnd-2_family-5 | rRNA | -250 | 4442 |
| Chr02:22384628-22385451 | 508 | 577 | rnd-2_family-5 | rRNA | -3412 | 1280 |
| Chr02:22384628-22385451 | 581 | 634 | rnd-2_family-5 | rRNA | -3413 | 1279 |
| Chr02:22385911-22391416 | 1 | 407 | rnd-2_family-5 | rRNA | -28 | 4664 |
| Chr02:22385911-22391416 | 52 | 483 | rnd-2_family-5 | rRNA | 4267 | 4685 |
| Chr02:22385911-22391416 | 408 | 740 | rnd-2_family-5 | rRNA | -109 | 4583 |
| Chr02:22385911-22391416 | 549 | 1097 | rnd-2_family-5 | rRNA | 4236 | 4686 |
| Chr02:22385911-22391416 | 787 | 5505 | rnd-2_family-5 | rRNA | 0 | 4692 |
| Chr02:22385920-22386235 | 1 | 315 | rnd-2_family-5 | rRNA | -37 | 4655 |
| Chr02:22386247-22386308 | 2 | 61 | rnd-2_family-5 | rRNA | -388 | 4304 |
| Chr02:22386344-22387232 | 9 | 664 | rnd-2_family-5 | rRNA | 4236 | 4686 |
| Chr02:22386344-22387232 | 354 | 888 | rnd-2_family-5 | rRNA | 0 | 4692 |
| Chr02:22387266-22387953 | 1 | 687 | rnd-2_family-5 | rRNA | -535 | 4157 |
| Chr02:22387371-22391423 | 1 | 4045 | rnd-2_family-5 | rRNA | -640 | 4052 |
| Chr02:22387970-22388203 | 1 | 233 | rnd-2_family-5 | rRNA | -1245 | 3447 |
| Chr02:22398432-22401443 | 1 | 37 | rnd-2_family-5 | rRNA | 4180 | 4216 |
| Chr02:22398432-22401443 | 1539 | 1608 | rnd-2_family-5 | rRNA | -3412 | 1280 |
| Chr02:22398432-22401443 | 1612 | 1665 | rnd-2_family-5 | rRNA | -3413 | 1279 |
| Chr02:22398432-22401443 | 2228 | 2405 | rnd-2_family-5 | rRNA | -84 | 4608 |
| Chr02:22398432-22401443 | 2234 | 2450 | rnd-2_family-5 | rRNA | 4465 | 4692 |
| Chr02:22398432-22401443 | 2406 | 2835 | rnd-2_family-5 | rRNA | 0 | 4692 |
| Chr02:22398432-22401443 | 2453 | 2886 | rnd-2_family-5 | rRNA | 4233 | 4685 |
| Chr02:22398432-22401443 | 2837 | 3011 | rnd-2_family-5 | rRNA | -185 | 4507 |
| Chr02:22399334-22400324 | 637 | 706 | rnd-2_family-5 | rRNA | -3412 | 1280 |
| Chr02:22399334-22400324 | 710 | 763 | rnd-2_family-5 | rRNA | -3413 | 1279 |
| Chr02:22399342-22400314 | 629 | 698 | rnd-2_family-5 | rRNA | -3412 | 1280 |
| Chr02:22399342-22400314 | 702 | 755 | rnd-2_family-5 | rRNA | -3413 | 1279 |
| Chr02:22400403-22401476 | 255 | 636 | rnd-2_family-5 | rRNA | 4316 | 4692 |
| Chr02:22400403-22401476 | 435 | 864 | rnd-2_family-5 | rRNA | 0 | 4692 |
| Chr02:22400403-22401476 | 661 | 1073 | rnd-2_family-5 | rRNA | 4233 | 4663 |
| Chr02:22400573-22400895 | 87 | 322 | rnd-2_family-5 | rRNA | -84 | 4608 |
| Chr02:22400940-22401001 | 2 | 61 | rnd-2_family-5 | rRNA | -388 | 4304 |
| Chr02:22409506-22411840 | 1404 | 2334 | rnd-2_family-5 | rRNA | 1 | 931 |
| Chr02:22409514-22413388 | 1396 | 3874 | rnd-2_family-5 | rRNA | 1 | 2479 |
| Chr02:22410204-22411039 | 706 | 835 | rnd-2_family-5 | rRNA | 1 | 130 |
| Chr02:22411851-22412147 | 1 | 296 | rnd-2_family-5 | rRNA | 943 | 1236 |
| Chr02:22412162-22412788 | 3 | 626 | rnd-2_family-5 | rRNA | 1256 | 1879 |
| Chr02:22412804-22412991 | 1 | 187 | rnd-2_family-5 | rRNA | 1896 | 2082 |
| Chr02:22413004-22413066 | 1 | 62 | rnd-2_family-5 | rRNA | 2096 | 2157 |
| Chr02:22413408-22413477 | 1 | 69 | rnd-2_family-5 | rRNA | 2500 | 2568 |
| Chr02:22413489-22413592 | 1 | 103 | rnd-2_family-5 | rRNA | 2581 | 2683 |
| Chr02:22442537-22443877 | 1 | 1336 | rnd-2_family-5 | rRNA | 2444 | 3780 |
| Chr02:22443304-22443516 | 1 | 212 | rnd-2_family-5 | rRNA | 3211 | 3422 |
| Chr02:22443319-22443861 | 1 | 542 | rnd-2_family-5 | rRNA | 3226 | 3768 |
| Chr02:22443533-22443873 | 1 | 340 | rnd-2_family-5 | rRNA | 3440 | 3780 |
| Chr02:22443987-22444113 | 1 | 125 | rnd-2_family-5 | rRNA | 3662 | 3782 |
| Chr02:22444038-22451781 | 1 | 921 | rnd-2_family-5 | rRNA | 3713 | 4692 |
| Chr02:22444038-22451781 | 738 | 1220 | rnd-2_family-5 | rRNA | -6 | 4686 |
| Chr02:22444038-22451781 | 1062 | 1392 | rnd-2_family-5 | rRNA | 4236 | 4608 |
| Chr02:22444038-22451781 | 3060 | 3099 | rnd-2_family-5 | rRNA | -476 | 4216 |
| Chr02:22444038-22451781 | 4489 | 5751 | rnd-2_family-5 | rRNA | 1 | 1265 |
| Chr02:22444038-22451781 | 5750 | 7743 | rnd-2_family-5 | rRNA | 1407 | 3390 |
| Chr02:22444043-22444113 | 1 | 69 | rnd-2_family-5 | rRNA | 3718 | 3782 |
| Chr02:22444127-22444170 | 1 | 42 | rnd-2_family-5 | rRNA | 3785 | 3826 |
| Chr02:22444127-22449758 | 1 | 832 | rnd-2_family-5 | rRNA | 3785 | 4692 |
| Chr02:22444127-22449758 | 649 | 1131 | rnd-2_family-5 | rRNA | -6 | 4686 |
| Chr02:22444127-22449758 | 973 | 1303 | rnd-2_family-5 | rRNA | 4236 | 4608 |
| Chr02:22444127-22449758 | 2971 | 3010 | rnd-2_family-5 | rRNA | -476 | 4216 |
| Chr02:22444127-22449758 | 4400 | 5631 | rnd-2_family-5 | rRNA | 1 | 1231 |
| Chr02:22444363-22445065 | 1 | 596 | rnd-2_family-5 | rRNA | 4092 | 4692 |
| Chr02:22444363-22445065 | 287 | 702 | rnd-2_family-5 | rRNA | -6 | 4686 |
| Chr02:22445099-22445378 | 1 | 279 | rnd-2_family-5 | rRNA | 4236 | 4538 |
| Chr02:22448026-22448678 | 501 | 652 | rnd-2_family-5 | rRNA | 1 | 152 |
| Chr02:22449122-22449268 | 1 | 146 | rnd-2_family-5 | rRNA | 597 | 742 |
| Chr02:22449812-22451781 | 1 | 1969 | rnd-2_family-5 | rRNA | 1432 | 3390 |
| Chr02:22450121-22450235 | 1 | 114 | rnd-2_family-5 | rRNA | 4412 | 4538 |
| Chr02:22454414-22455498 | 2 | 227 | rnd-2_family-5 | rRNA | 4370 | 4692 |
| Chr02:22454414-22455498 | 785 | 838 | rnd-2_family-5 | rRNA | 1220 | 1279 |
| Chr02:22454414-22455498 | 842 | 911 | rnd-2_family-5 | rRNA | 1205 | 1280 |
| Chr02:22454416-22461960 | 9 | 225 | rnd-2_family-5 | rRNA | 4370 | 4608 |
| Chr02:22454416-22461960 | 783 | 836 | rnd-2_family-5 | rRNA | 1220 | 1279 |
| Chr02:22454416-22461960 | 840 | 909 | rnd-2_family-5 | rRNA | 1205 | 1280 |
| Chr02:22454416-22461960 | 2373 | 2411 | rnd-2_family-5 | rRNA | -476 | 4216 |
| Chr02:22454416-22461960 | 3766 | 7544 | rnd-2_family-5 | rRNA | 1 | 3865 |
| Chr02:22461623-22461823 | 1 | 200 | rnd-2_family-5 | rRNA | 3534 | 3735 |
| Chr02:22462028-22462360 | 1 | 297 | rnd-2_family-5 | rRNA | 3720 | 4045 |
| Chr08:17700-29944 | 9 | 10033 | rnd-2_family-8 | Unknown | 45 | 715 |
| Chr08:17700-29944 | 10097 | 12152 | rnd-2_family-8 | Unknown | 59 | 715 |
| Chr08:17700-29991 | 9 | 10033 | rnd-2_family-8 | Unknown | 45 | 715 |
| Chr08:17700-29991 | 10097 | 12291 | rnd-2_family-8 | Unknown | 57 | 715 |
| Chr09:28437603-28437722 | 1 | 119 | NC_040010.1 | Mitochondrial | 267646 | 267764 |
| Chr09:28439381-28439435 | 1 | 54 | NC_040010.1 | Mitochondrial | 269402 | 269455 |
| Chr09:28439504-28439703 | 1 | 199 | NC_040010.1 | Mitochondrial | 269525 | 269723 |
| Chr09:28439892-28440021 | 1 | 129 | NC_040010.1 | Mitochondrial | 269913 | 270041 |
| Chr09:28444447-28445247 | 1 | 800 | NC_040010.1 | Mitochondrial | 274470 | 275269 |
| Chr09:28445380-28445475 | 1 | 95 | NC_040010.1 | Mitochondrial | 275403 | 275497 |
| Chr09:28446057-28446129 | 1 | 72 | NC_040010.1 | Mitochondrial | 276080 | 276151 |
| Chr09:28446336-28446999 | 1 | 663 | NC_040010.1 | Mitochondrial | 276359 | 277019 |
| Chr09:28447412-28448014 | 1 | 602 | NC_040010.1 | Mitochondrial | 277433 | 278034 |
| Chr09:28448187-28448338 | 1 | 151 | NC_040010.1 | Mitochondrial | 278208 | 278364 |
| Chr09:28451258-28451561 | 1 | 303 | NC_040010.1 | Mitochondrial | 281286 | 281588 |
| Chr09:28456459-28456580 | 1 | 121 | NC_040010.1 | Mitochondrial | 286490 | 286610 |
| Chr09:28456671-28456785 | 1 | 114 | NC_040010.1 | Mitochondrial | 286702 | 286815 |
| Chr09:28457605-28457827 | 1 | 222 | NC_040010.1 | Mitochondrial | 287636 | 287857 |
| Chr09:28457906-28458525 | 1 | 619 | NC_040010.1 | Mitochondrial | 287937 | 288556 |
| Chr09:28458599-28458877 | 1 | 278 | NC_040010.1 | Mitochondrial | 288627 | 288909 |
| Chr09:28458927-28459128 | 1 | 201 | NC_040010.1 | Mitochondrial | 288960 | 289160 |
| Chr09:28459527-28459559 | 1 | 32 | NC_040010.1 | Mitochondrial | 19954 | 19923 |
| Chr09:28459575-28459874 | 1 | 299 | NC_040010.1 | Mitochondrial | 289603 | 289901 |
| Chr09:28468142-28471705 | 2281 | 3563 | MG925369.1 | chloroplast | 100626 | 101908 |
| Chr09:28468142-28471705 | 1 | 38 | NC_040010.1 | Mitochondrial | 7407 | 7370 |
| Chr09:28468142-28475684 | 2281 | 7542 | MG925369.1 | chloroplast | 100626 | 105890 |
| Chr09:28468142-28475684 | 5689 | 7048 | NC_040010.1 | Mitochondrial | 476331 | 477690 |
| Chr09:28472443-28473000 | 1 | 557 | MG925369.1 | chloroplast | 102646 | 103202 |
| Chr09:28473263-28473981 | 1 | 718 | MG925369.1 | chloroplast | 103466 | 104183 |
| Chr09:28473263-28473981 | 568 | 718 | NC_040010.1 | Mitochondrial | 476331 | 476481 |
| Chr09:28474046-28474262 | 1 | 216 | MG925369.1 | chloroplast | 104249 | 104464 |
| Chr09:28474046-28474262 | 1 | 216 | NC_040010.1 | Mitochondrial | 476547 | 476762 |
| Chr09:28474377-28475065 | 1 | 688 | MG925369.1 | chloroplast | 104580 | 105267 |
| Chr09:28474377-28475065 | 1 | 688 | NC_040010.1 | Mitochondrial | 476878 | 477565 |
| Chr09:28475083-28475314 | 1 | 231 | MG925369.1 | chloroplast | 105286 | 105516 |
| Chr09:28475083-28475314 | 6 | 231 | NC_040010.1 | Mitochondrial | 477690 | 477915 |
| Chr09:28477624-28479611 | 1 | 495 | MG925369.1 | chloroplast | 106773 | 107267 |
| Chr09:28477624-28479611 | 897 | 1987 | NC_040010.1 | Mitochondrial | 291070 | 292160 |
| Chr09:28477862-28479115 | 1 | 257 | MG925369.1 | chloroplast | 107011 | 107267 |
| Chr09:28477862-28479115 | 659 | 1253 | NC_040010.1 | Mitochondrial | 291070 | 291664 |
| Chr09:28479734-28479938 | 1 | 204 | NC_040010.1 | Mitochondrial | 292284 | 292487 |
| Chr09:28479971-28480041 | 1 | 70 | NC_040010.1 | Mitochondrial | 292521 | 292590 |
| Chr09:28480066-28480228 | 1 | 162 | NC_040010.1 | Mitochondrial | 292616 | 292777 |
| Chr09:28484235-28484573 | 1 | 338 | NC_040010.1 | Mitochondrial | 296786 | 297123 |
| Chr09:28488566-28488768 | 1 | 202 | NC_040010.1 | Mitochondrial | 301181 | 301382 |
| Chr09:28488888-28488998 | 1 | 110 | NC_040010.1 | Mitochondrial | 301503 | 301612 |
| Chr09:28489249-28489422 | 1 | 173 | NC_040010.1 | Mitochondrial | 301864 | 302036 |
| Chr09:28489454-28489672 | 1 | 218 | NC_040010.1 | Mitochondrial | 302069 | 302286 |
| Chr09:28489961-28490878 | 1 | 917 | NC_040010.1 | Mitochondrial | 302578 | 303494 |
| Chr09:28491456-28491654 | 1 | 198 | NC_040010.1 | Mitochondrial | 304121 | 304318 |
| Chr09:28492672-28492783 | 1 | 111 | NC_040010.1 | Mitochondrial | 305244 | 305354 |
| Chr09:28494838-28495447 | 1 | 609 | NC_040010.1 | Mitochondrial | 307413 | 308021 |
| Chr09:28495983-28496570 | 1 | 587 | NC_040010.1 | Mitochondrial | 308558 | 309144 |
| Chr09:28498517-28498785 | 1 | 268 | NC_040010.1 | Mitochondrial | 311151 | 311418 |
| Chr09:28498872-28499502 | 1 | 630 | NC_040010.1 | Mitochondrial | 311506 | 312135 |
| Chr10:54026-54853 | 3 | 827 | rnd-2_family-8 | Unknown | 1 | 715 |
| Chr10:55590-56226 | 1 | 636 | rnd-2_family-8 | Unknown | 1 | 715 |
| Chr10:61671-78118 | 1 | 4864 | rnd-2_family-8 | Unknown | 1 | 715 |
| Chr10:61671-78118 | 4969 | 7441 | rnd-2_family-8 | Unknown | 1 | 715 |
| Chr10:61671-78118 | 7451 | 8400 | rnd-2_family-8 | Unknown | 1 | 715 |
| Chr10:61671-78118 | 8544 | 16447 | rnd-2_family-8 | Unknown | 1 | 715 |
| Chr10:61677-78127 | 1 | 4858 | rnd-2_family-8 | Unknown | 1 | 715 |
| Chr10:61677-78127 | 4963 | 7435 | rnd-2_family-8 | Unknown | 1 | 715 |
| Chr10:61677-78127 | 7445 | 8394 | rnd-2_family-8 | Unknown | 1 | 715 |
| Chr10:61677-78127 | 8538 | 16450 | rnd-2_family-8 | Unknown | 1 | 715 |
| Chr10:61806-62087 | 2 | 281 | rnd-2_family-8 | Unknown | 93 | 372 |
| Chr10:62327-62714 | 1 | 387 | rnd-2_family-8 | Unknown | 65 | 451 |
| Chr10:62787-63009 | 1 | 220 | rnd-2_family-8 | Unknown | 159 | 379 |
| Chr10:63088-63199 | 1 | 111 | rnd-2_family-8 | Unknown | 110 | 220 |
| Chr10:63300-63579 | 1 | 279 | rnd-2_family-8 | Unknown | 154 | 432 |
| Chr10:63662-63721 | 1 | 59 | rnd-2_family-8 | Unknown | 150 | 208 |
| Chr10:63847-64145 | 1 | 298 | rnd-2_family-8 | Unknown | 149 | 446 |
| Chr10:64221-64253 | 1 | 32 | rnd-2_family-8 | Unknown | 157 | 188 |
| Chr10:64302-64401 | 1 | 99 | rnd-2_family-8 | Unknown | 238 | 336 |
| Chr10:64491-64923 | 1 | 431 | rnd-2_family-8 | Unknown | 62 | 492 |
| Chr10:65069-65252 | 1 | 182 | rnd-2_family-8 | Unknown | 91 | 272 |
| Chr10:65311-65350 | 1 | 39 | rnd-2_family-8 | Unknown | 150 | 188 |
| Chr10:65413-65573 | 1 | 160 | rnd-2_family-8 | Unknown | 251 | 410 |
| Chr10:65666-65848 | 1 | 182 | rnd-2_family-8 | Unknown | 153 | 334 |
| Chr10:66040-66344 | 1 | 304 | rnd-2_family-8 | Unknown | 162 | 465 |
| Chr10:66693-66733 | 1 | 40 | rnd-2_family-8 | Unknown | 154 | 193 |
| Chr10:66796-67079 | 1 | 283 | rnd-2_family-8 | Unknown | 89 | 371 |
| Chr10:67214-67267 | 1 | 53 | rnd-2_family-8 | Unknown | 141 | 193 |
| Chr10:67370-67645 | 2 | 275 | rnd-2_family-8 | Unknown | 115 | 388 |
| Chr10:67687-67788 | 1 | 101 | rnd-2_family-8 | Unknown | 67 | 167 |
| Chr10:67864-68055 | 1 | 191 | rnd-2_family-8 | Unknown | 185 | 375 |
| Chr10:68418-68855 | 1 | 437 | rnd-2_family-8 | Unknown | 20 | 457 |
| Chr10:68972-69041 | 1 | 69 | rnd-2_family-8 | Unknown | 26 | 94 |
| Chr10:69522-69720 | 1 | 198 | rnd-2_family-8 | Unknown | 150 | 347 |
| Chr10:69893-70035 | 1 | 142 | rnd-2_family-8 | Unknown | 155 | 296 |
| Chr10:70710-70917 | 1 | 207 | rnd-2_family-8 | Unknown | 477 | 683 |
| Chr10:71187-71258 | 1 | 71 | rnd-2_family-8 | Unknown | 590 | 660 |
| Chr10:71331-71404 | 1 | 73 | rnd-2_family-8 | Unknown | 2 | 74 |
| Chr10:71425-71631 | 1 | 206 | rnd-2_family-8 | Unknown | 462 | 667 |
| Chr10:71848-71961 | 1 | 113 | rnd-2_family-8 | Unknown | 159 | 271 |
| Chr10:72056-72114 | 1 | 58 | rnd-2_family-8 | Unknown | 1 | 58 |
| Chr10:72553-72589 | 1 | 36 | rnd-2_family-8 | Unknown | 500 | 535 |
| Chr10:72621-72787 | 1 | 166 | rnd-2_family-8 | Unknown | 202 | 367 |
| Chr10:73091-73319 | 1 | 228 | rnd-2_family-8 | Unknown | 123 | 350 |
| Chr10:73563-74054 | 1 | 491 | rnd-2_family-8 | Unknown | 46 | 535 |
| Chr10:74094-74274 | 1 | 179 | rnd-2_family-8 | Unknown | 28 | 206 |
| Chr10:74463-74500 | 1 | 37 | rnd-2_family-8 | Unknown | 33 | 69 |
| Chr10:74527-74758 | 1 | 231 | rnd-2_family-8 | Unknown | 463 | 693 |
| Chr10:74780-75700 | 18 | 915 | rnd-2_family-8 | Unknown | 1 | 715 |
| Chr10:75712-75794 | 2 | 82 | rnd-2_family-8 | Unknown | 1 | 81 |
| Chr10:76052-76369 | 1 | 316 | rnd-2_family-8 | Unknown | 340 | 655 |
| Chr10:76628-77345 | 3 | 717 | rnd-2_family-8 | Unknown | 1 | 715 |
| Chr10:77430-77804 | 1 | 374 | rnd-2_family-8 | Unknown | 69 | 441 |
| Chr10:77822-77855 | 1 | 33 | rnd-2_family-8 | Unknown | 277 | 309 |
| Chr11:24028678-24032420 | 1 | 523 | rnd-2_family-9 | Unknown | 581 | 1192 |
| Chr11:24042713-24042789 | 5 | 76 | rnd-2_family-9 | Unknown | 1 | 72 |
| Chr11:24042902-24043101 | 1 | 199 | rnd-2_family-9 | Unknown | 189 | 387 |
| Chr11:24043204-24046225 | 1 | 698 | rnd-2_family-9 | Unknown | 491 | 1190 |
| Chr11:24049224-24049797 | 1 | 573 | rnd-2_family-9 | Unknown | 2 | 574 |
| Chr11:24049861-24051475 | 1 | 545 | rnd-2_family-9 | Unknown | 644 | 1193 |
| Chr11:24070256-24071621 | 172 | 1362 | rnd-2_family-9 | Unknown | 1 | 1193 |
| Chr11:24080480-24083129 | 30 | 584 | rnd-2_family-9 | Unknown | 177 | 766 |
| Chr11:24080480-24083129 | 602 | 950 | rnd-2_family-9 | Unknown | 853 | 1192 |
| Chr11:24084573-24085764 | 358 | 444 | rnd-2_family-9 | Unknown | 193 | 282 |
| Chr11:24084573-24085764 | 451 | 995 | rnd-2_family-9 | Unknown | 559 | 1192 |
| Chr11:24096497-24098093 | 685 | 1596 | rnd-2_family-9 | Unknown | 1 | 908 |
| Chr11:24098339-24098495 | 1 | 34 | rnd-2_family-9 | Unknown | 1160 | 1193 |

**Supplementary Figure 13.**
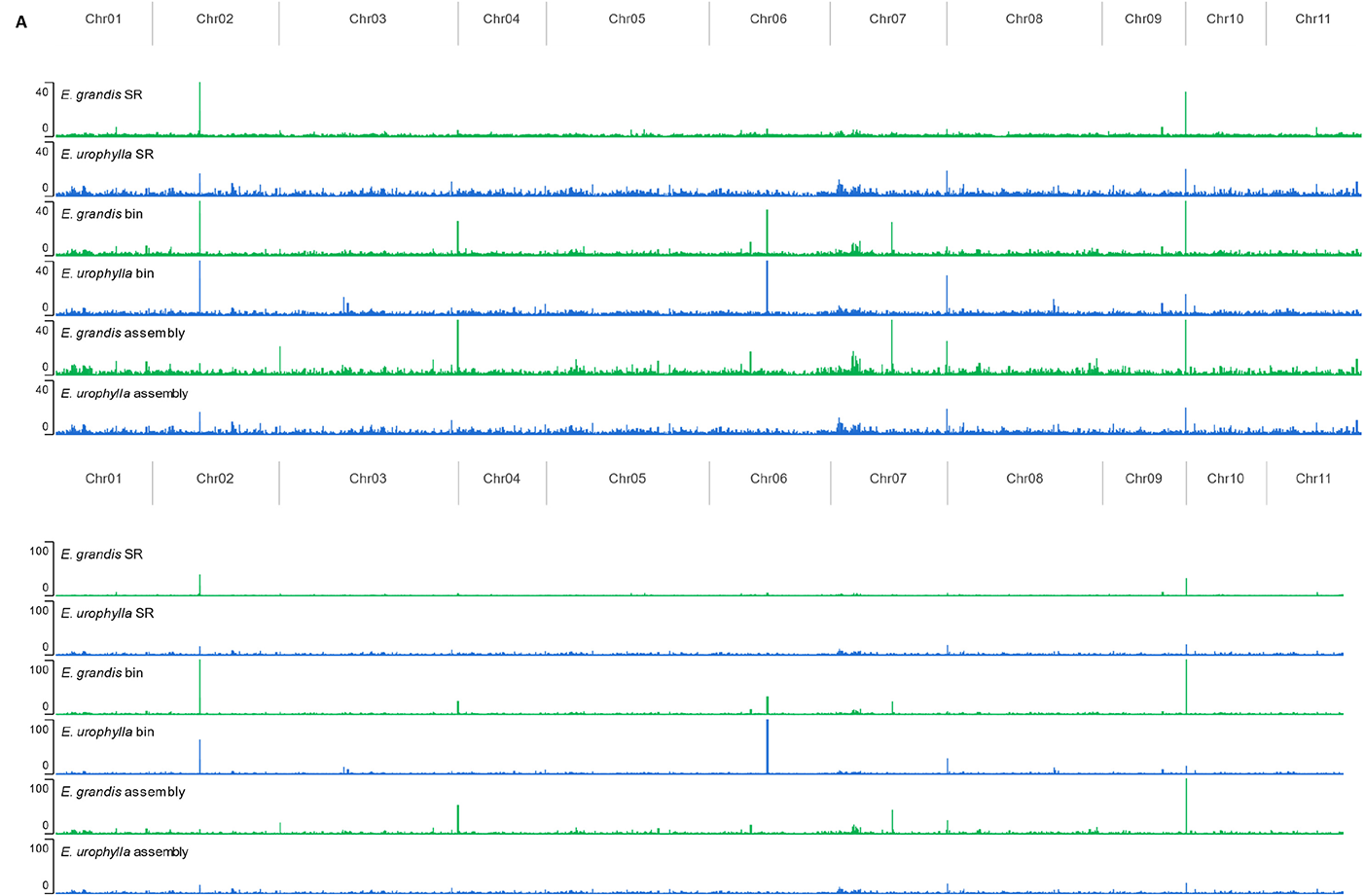
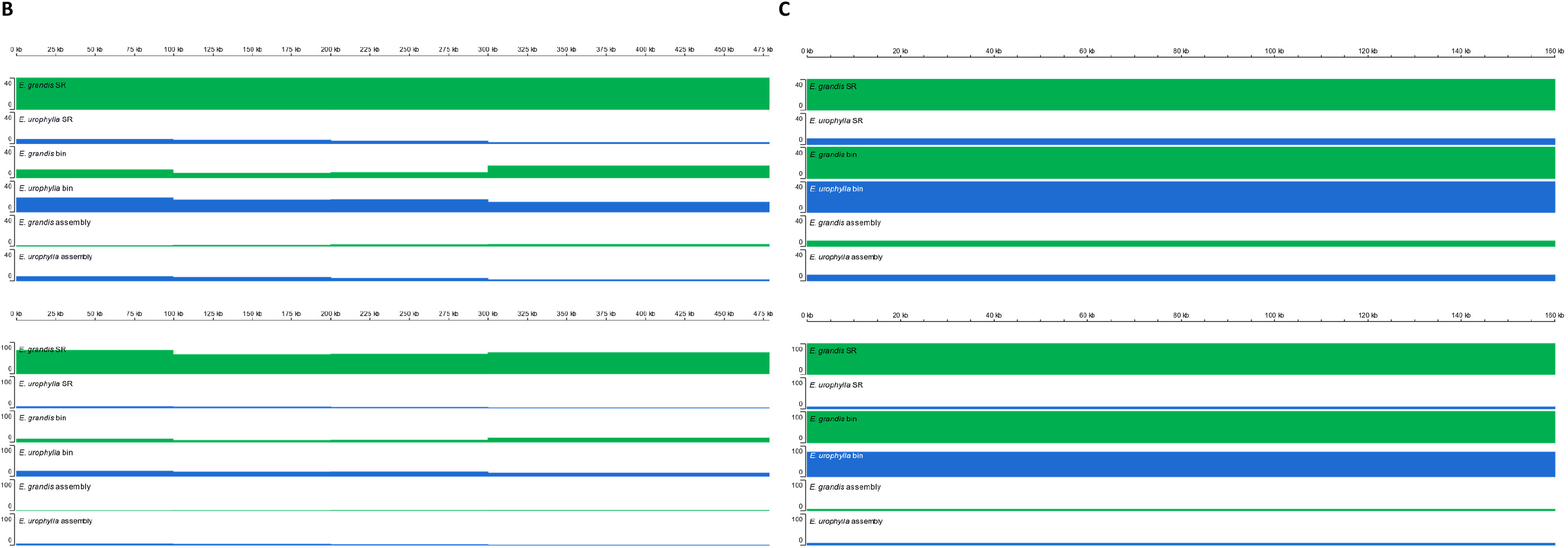
Genome coverage of the *E. grandis* v2.0 nuclear reference and plastid genomes. (A) Alignment of *E. grandis* (FK1758, green) and *E. urophylla* (FK1756, blue) parental short-read (SR), binned long-read sequencing data and haplogenome assemblies (contigs) to the *E. grandis* v2.0 reference genome (Myburg *et al*., 2014; Bartholome *et al*., 2015). Coverage is shown on the y-axis, with max coverage parameters set to 40X (top panel) and 100X (bottom panel), along the eleven *Eucalyptus* chromosomes in bins of 100 kb shown on the x-axis. Alignment of the same sequencing data and assemblies to the *E. grandis* (B) mitochondrial (478.8 kb) and (C) chloroplast (160.1 kb) genomes (Pinard *et al*., 2019), at 40X (top panel) and 100X (bottom panel) maximum coverage. All alignments were viewed in the IGV browser and bins were 100 kb in size.

**Supplementary Figure 14.**
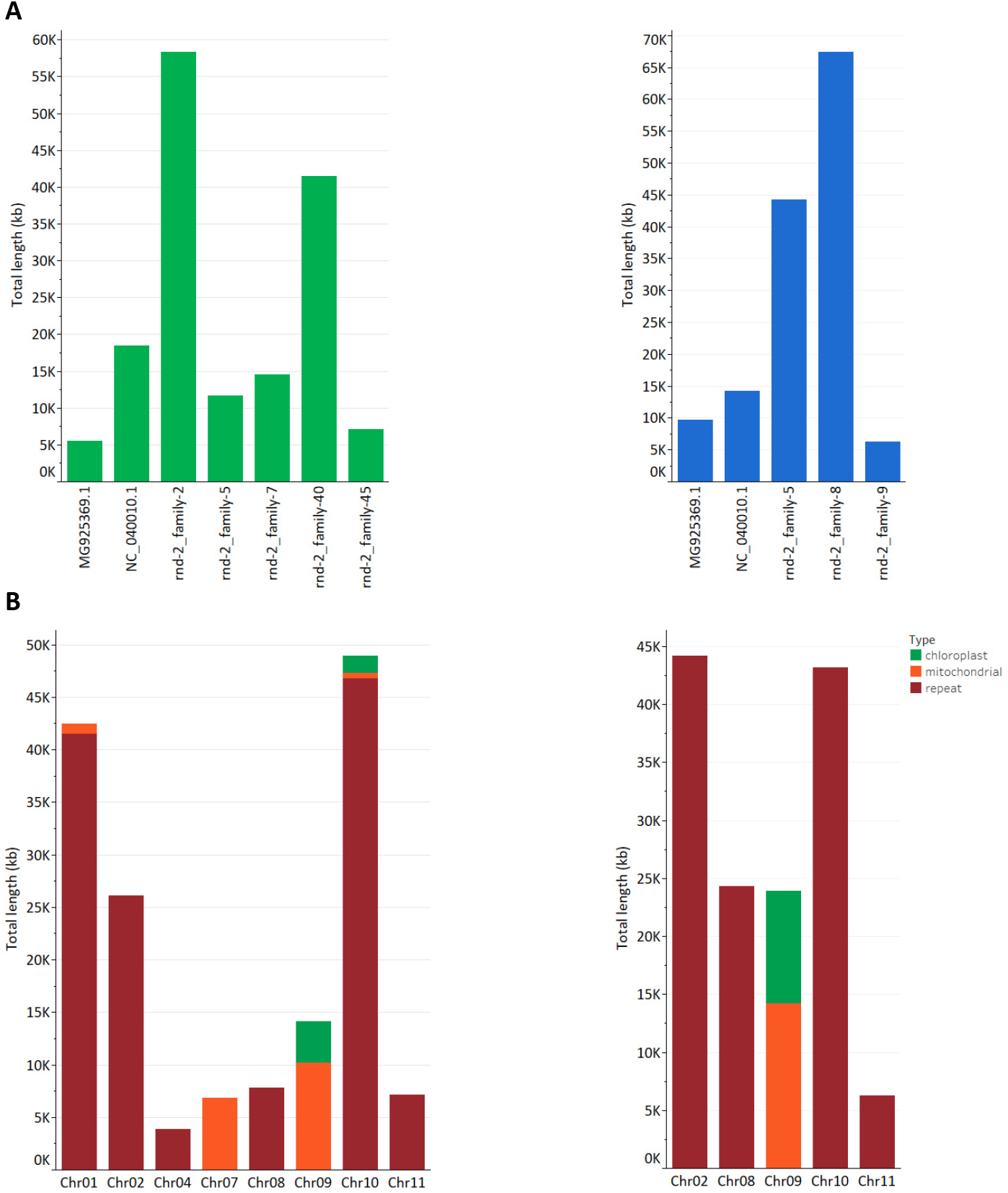
Summary of the total size and type of elements found in high genome coverage bins. Organellar introgression was identified through BLAST analysis to the *E. grandis* plastid genomes (Pinard *et al*., 2019), while repeat elements were identified with RepeatMasker. (A) The total size of different type of elements found in high coverage bins (see) for the *E. grandis* (green) and *E. urophylla* (blue) alignments. The element type is indicated on the x-axis as either mitochondrial (NC040010.1), chloroplast (MG925369.1) or repeat elements (rnd, the repeat family/class is given) and the total length the element contributes to all high coverage bins is given in kb (kilobases) on the y-axis. (B) The total length of different types of elements contributed per chromosome within high coverage bins. The chromosomes are indicated on the x-axis for *E. grandis* (left) and *E. urophylla* (right) and the total length contributed by each element is given on the y-axis. Contributions are either repetitive elements in red, mitochondrial introgression in orange or chloroplast introgression in green. Note that in both cases chromosome 9 only has organellar introgression, whereas the majority of other chromosomes have mostly repeat elements.

## Notes

### Competing Interest Statement

The authors have declared no competing interest.

